# Population genetics of active transposable elements: overdispersion arises naturally from transposition-deletion-selection balance

**DOI:** 10.1101/2025.11.27.691047

**Authors:** Adekanmi Daniel Omole, Peter Czuppon

## Abstract

Active transposable element (TE) families often show overdispersion (variance greater than the mean), which is in contrast to the predictions of classical Poisson-based models and related theories built on the same approximation. To address this gap, we develop a diploid stochastic model of TE dynamics under free recombination, based on a biparental Moran model with transposition, excision, purifying selection, and derive mean and variance dynamics of TE copy numbers at two stages of the life cycle: before and after transposition and excision. We show that overdispersion arises naturally through transposition generating positive linkage disequilibrium between TE insertion sites with the disequilibrium persisting even under free recombination. The higher the transposition rate, the stronger the resulting overdispersion. Underdispersion and Poisson-like variation are instead stage-dependent: neither is expected after transposition and excision, while both remain possible beforehand, governed by the curvature of the fitness function. We further show that maintaining positive equilibrium copy numbers, and thus sustaining overdispersion, requires the net transposition rate to remain below roughly `0.5` insertions per copy per generation, a constraint satisfied by observed natural populations to maintain genome stability. A qualitative comparison with the DPGP3 Zambian *Drosophila melanogaster* dataset, focusing on TE insertions in high-recombination euchromatic regions shows that most TE families are overdispersed and that many also exhibit the predicted right-skewness and excess-kurtosis patterns, which are predicted by our model. These results identify overdispersion as a natural outcome of active TE dynamics and provide a mechanistic framework for understanding the full distribution of TE copy numbers in highly recombining regions.

**Statement of significance:** This study addresses a mismatch between population genetic theory and empirical data on transposable elements (TEs). A detailed analysis of the stochastic model underlying the classical population genetics theory of TEs provides new predictions on the mean and variance of TE copy numbers that accurately match simulation results, and provide an explanation for the right-skewed, heavy-tailed distributions seen in real data. Importantly, we show that the pronounced variability in TE copy numbers, i.e. overdispersion, observed in natural populations emerges naturally from ongoing TE dynamics, even in highly recombining genomic regions. How much overdispersion is predicted further depends on the stage of the life cycle at which copy numbers are counted, before or after transposition and excision.

## 1 Introduction

Transposable elements (TEs) are abundant genomic sequences in eukaryotic genomes that influence the structure, function, and evolution of genomes (Bourque et al., 2018; Kazazian, 2004; Shapiro, 2012). These “jumping genes” replicate themselves and insert new copies throughout the genome, creating evolutionary tension between their selfish replication and their potentially deleterious effects, which create selective pressures acting on their hosts (Doolittle and Sapienza, 1980; Kelleher et al., 2020; Orgel and Crick, 1980). Understanding the TE copy number distribution in populations and species is therefore fundamental to comprehending genome evolution, adaptive processes, and the maintenance of genomic stability (Betancourt et al., 2024; Le Rouzic et al., 2007; Schrader and Schmitz, 2019).

The population genetics of TEs was first formalized by Charlesworth and Charlesworth (1983), who established a foundational model assuming free recombination and Poisson-distributed TE copy numbers, where the variance and mean of the TE copy number distribution are equal. This ‘equidispersion’ assumption facilitated analytical progress and provided an early prediction for equilibrium copy numbers under the balance between transposition and purifying selection (Charlesworth and Langley, 1986; Charlesworth and Charlesworth, 1983; Huang and Lee, 2024; Le Rouzic and Capy, 2006b; Scarpa and Kofler, 2023). However, empirical data show that TE copy number distributions often deviate from Poisson predictions, with active TE families typically exhibiting overdispersion (variance greater than the mean) along with higher-order distributional features such as right-skewness and heavy tails (kurtosis) (Lee, 2022; McGurk et al., 2021; Said et al., 2022). This mismatch between classical theory and empirical observations was pointed out by Smith et al. (2022). To explain overdispersion, their model relies on continuous immigration of TEs into the host genome to maintain positive TE copy numbers. However, biological observation suggests that horizontal transfer of TE families is rare (Kofler et al., 2015; Wierzbicki et al., 2025), which contrasts with the fundamental assumption of their model. In *Drosophila*, for instance, the estimated rate is approximately `0.04` horizontal transfer events per TE family per million years (Bartolomé et al., 2009).

Beyond that model, several biological explanations have been proposed to explain the mismatch between theory and data, including episodic bursts of TE activity rather than steady accumulation, insertion biases toward specific genomic regions, variation in the strength of selection across genomic locations, population structure, and positive linkage disequilibrium between TE insertions (Langmüller et al., 2023, 2025; McGurk et al., 2021; Oggenfuss and Croll, 2023; Roze, 2023). Importantly, the modeling study by Roze (2023) showed that positive linkage disequilibrium among insertion sites generated by transposition can inflate TE copy number variance above the Poisson baseline, especially for low recombination rates. A limitation of that study is that the mean and variance are not computed from the same model: the mean is approximated using the Poisson distribution assumption from Charlesworth and Charlesworth (1983), and then subsequently a correction to the variance is calculated. This inflates both moments, though under free recombination the effect is small at low transposition rates.

For higher transposition rates, however, recent theoretical work shows the limits of the Poisson assumption even against simulations. For example, Tomar et al. (2023) model TE regulation by piRNA silencing, which is triggered only once a copy inserts into a piRNA cluster and therefore requires a high transposition rate (*u* = `0.1`) for the family to amplify first. At that rate the Poisson assumption fails, since transposition adds copies preferentially to already TE-rich genomes, and they resort to an ad-hoc correction of the variance of TE copy numbers to match theoretical and simulated mean TE copy numbers. Similarly, simulation data from Huang and Lee (2024) also display observable overdispersion in TE copy numbers, i.e., the variance is larger than the mean. Together, these observed differences between theory, simulations and empirical data underscore the need for a joint derivation of mean and variance of TE copy numbers to explain overdispersion under free recombination from first principles.

To address this gap, we study the biparental Moran model of TE copy numbers under free recombination in two life-cycle stages (pre- and post-transposition/excision), which underlies the classical population genetics model (Charlesworth and Charlesworth, 1983). We demonstrate that overdispersion arises naturally in this model: the transposition process generates positive linkage disequilibrium among TE insertions, which free recombination weakens but does not remove completely, while excision and purifying selection set the mean copy number at equilibrium. Moreover, we find that this depends on the life-cycle stage: post-transposition/excision, active TEs are unlikely to be underdispersed or Poisson-distributed (though close to Poisson), since these outcomes require a delicate balance between transposition and selection that typically leads to TE loss. Contrarily, pre-transposition/excision, under-, equi- or overdispersion of TE copy numbers are all possible, which is governed by the curvature of the fitness function. Additionally, we derive the dynamics of the mean and variance of the TE copy number distribution, which show the impact of higher-order moments of the distribution, including skewness and excess kurtosis. More generally, our model predicts right-skewness and heavy-tailed distributions of TE copy numbers. Finally, we compare our predictions qualitatively to TE counts in the DPGP3 Zambian *Drosophila melanogaster* dataset of Lee (2022) in euchromatic regions of the genome. Most of the TE families are overdispersed and many also display the right-skewness and excess-kurtosis patterns predicted by our analysis. Together, these results establish that a low level of overdispersion is a natural outcome of active TE dynamics even under free recombination, while also indicating what remains to be explained in more strongly linked genomic settings.

## 2 Results

The Moran model underlying our analysis is detailed in section 4.1. Key ingredients are TE transposition at rate *u*, TE excision at rate *v*, and natural selection acting during reproduction of individuals. The fitness function, denoted *w*(*n*), depends on the total number of TEs *n*contained in the diploid genome of an individual. From this Moran model formulation, we can derive the mean *µ* and variance *σ*^2^ dynamics of the TE copy number per diploid genome as (see section S1 in the supplementary material for the full derivation):

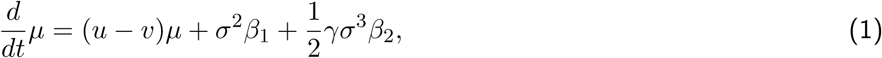

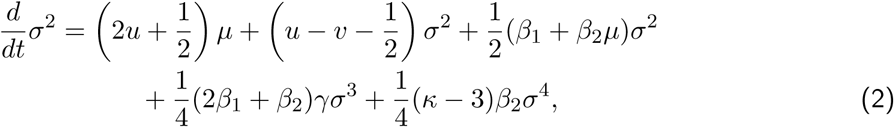

where *β*_1_ = *∂_µ_* ln *w*(*µ*) and 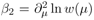 ln *w*(*µ*) capture how selection strength changes with TE copy number, *γ* and *κ −* 3 denote skewness and excess kurtosis of the TE distribution, respectively. Both moments are the copy number measured after a complete birth event, once parental sampling, recombination, and the subsequent transposition and excision have occurred. The earlier stage of the life cycle, immediately after recombination but before transposition and excision, is treated separately in Section 2.3.

To gain insight into how the TE copy number changes over time, we analyze these deterministic mean and variance dynamics derived from the stochastic model. As a central quantity to compare the variance with the mean, we will study the dispersion ratio 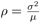, which quantifies the deviation of the variance from the Poisson expectation. Values of *ρ >* 1 indicate overdispersion; *ρ* = 1 corresponds to equidispersion, i.e., the Poisson case; and values of *ρ <* 1 indicate underdispersion.

We now present two methods for approximating the dispersion parameter. Both methods show that TE copy numbers tend to be naturally overdispersed, which is consistent with empirical observations, as we argue below. The first approach is based on an approximate moment closure. Here, we neglect the skewness *γ* and the excess kurtosis (*κ −* 3), i.e., we implicitly assume that the TE copy number is approximately normally distributed at all points in time. The second approach uses a parametric closure based on the negative binomial distribution. Unlike the approximation above, this method retains skewness and kurtosis by fitting the theoretical moments to expressions of the negative binomial distribution. This yields an exact expression for the dynamics that more fully captures the shape and variability of the TE copy-number distribution.

### 2.1 Approximate moment closure

In this section, we will assume that the terms involving skewness *γ* and excess kurtosis (*κ −* 3) are negligible compared to the other terms determining the dynamics of the mean and variance of TE copy numbers (Eq. (2)). To clarify that this is an approximation, we add a subscript, ‘app’ to the mean and variance. Similarly, we now also explicitly indicate that *β*_1_ and *β*_2_ depend on the mean TE copy number, as is typically the case. We then study the simplified system given by

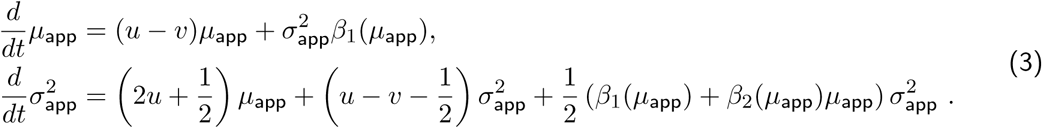

It is not possible to solve for the equilibrium values 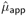 and 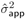 in general due to the dependence of *β*_1_ and *β*_2_ on the mean. However, it is possible to identify an equilibrium condition for the dispersion ratio 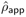 (see Appendix A):

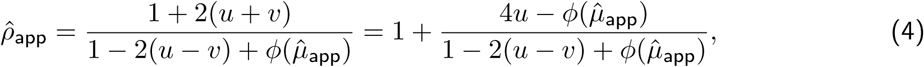

where 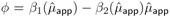. Rewriting 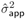 in the equilibrium equation for 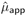, we then find the following condition for the transposition-deletion-selection balance:

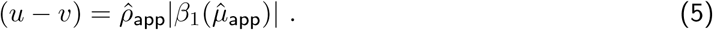

This states that in equilibrium, the overall purifying selection strength times the dispersion ratio, *ρ*_app_ (right-hand side) needs to balance the net transposition rate *u − v*. Since, as we will show below in Section 2.6, the dispersion ratio is typically greater than 1, this implies that purifying selection in equilibrium will be weaker than the net transposition rate.

To illustrate these general conclusions, we now study the Gaussian fitness function *w*(*n*) = exp(*−sn*^2^). In this case, the term *ϕ* in Eq. (4) is equal to zero (see Eq. (B.4)), resulting in 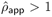, provided that the net transposition rate *u − v* is smaller than 1*/*2. Therefore, the overdispersion of TE copy numbers in equilibrium arises naturally from transposition-deletion-selection processes. Moreover, in this case the equilibrium mean, variance and dispersio ratio can be computed explicitly from Eq. (3) and are given by

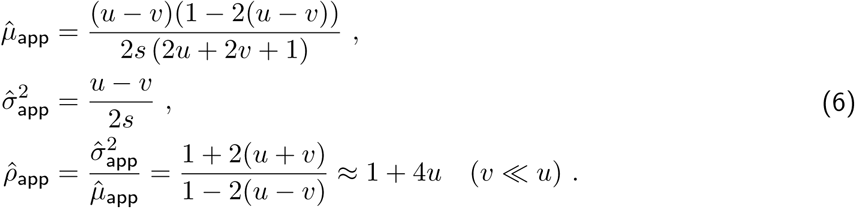

Note that for the mean TE copy number to remain positive at equilibrium, we again find that the net transposition rate must not exceed 1*/*2 new insertions per copy per generation. When *u − v >* 1*/*2, the variance in TE copy number increases without bound (see Eq. (3)), resulting in an unstable system in which the mean TE copy number eventually approaches zero and no stable equilibrium is maintained.

For comparison, we now briefly outline the classic mean dynamics described by Charlesworth and Charlesworth (1983) under the assumption of Poisson-distributed TE copy numbers, i.e., 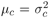. The index *c* indicates the classical TE models, and the dynamics of the mean TE copy number is given as (Charlesworth and Charlesworth, 1983)

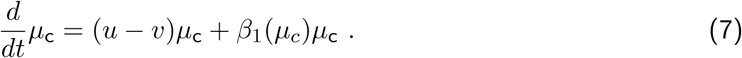

At equilibrium, assuming the Gaussian fitness function *w*(*n*) = exp(*−sn*^2^), the equilibrium mean and variance in the classical model are

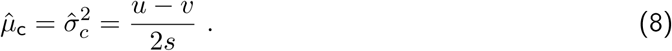

Comparing this to our analytical solution in Eq. (6), we see that the classic equilibrium mean copy number 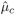 matches the approximate variance, 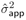, but not the mean. This pattern is illustrated and compared with stochastic simulations in Fig. 1a. Consistent with our theoretical results, we observe that the approximate variance, 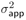, is greater than the approximate mean, μ_app_, showing the overdispersion in TE copy numbers. The high transposition rate used in Fig. 1a, *u* = 0.026 is chosen to make the overdispersion dynamics, that is, the separation between variance and mean, clearly visible. We consider a broader range of transposition rates later in Fig. 3. We also note that under the classical population genetics scaling for large population sizes, i.e., *u, v* and *s* of order 1*/N* and *N* going to infinity, our approximated mean 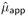 will tend to the approximated variance 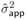 and hence the classical mean 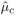 (see also Adeosun and Pfaffelhuber (2026) for a formal derivation).

**Figure 1:**
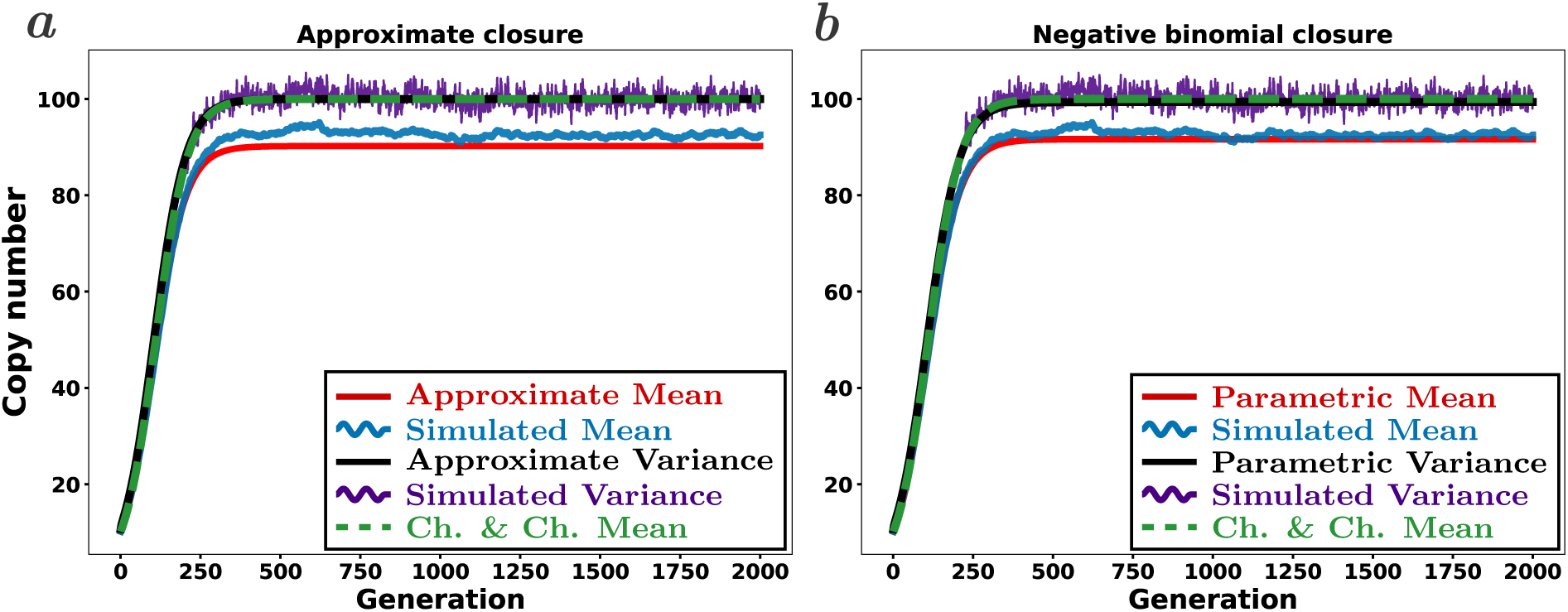
Comparison of theoretical dynamics of TE copy numbers to stochastic simulations, post-transposition/excision. The graphs show the a) approximate and b) parametric closure (straight lines) and stochastic simulation (wavy lines) of mean and variance dynamics of active TE copy numbers, including the Charlesworth and Charlesworth (1983) mean over time (dashed lines). The approximate dynamics (left) are computed using Eq. (3). The negative binomial closure dynamics (right) are computed using Eq. (12). Parameter values are *u* = 0.026*, v* = 0.006*, N* = 10^4^*, s* = 1*/N,* with free recombination and a Gaussian fitness function *w*(*n*) = exp(*−sn*^2^). The resulting mean and variance values are given in Supplementary Table S2.

**Figure 2:**
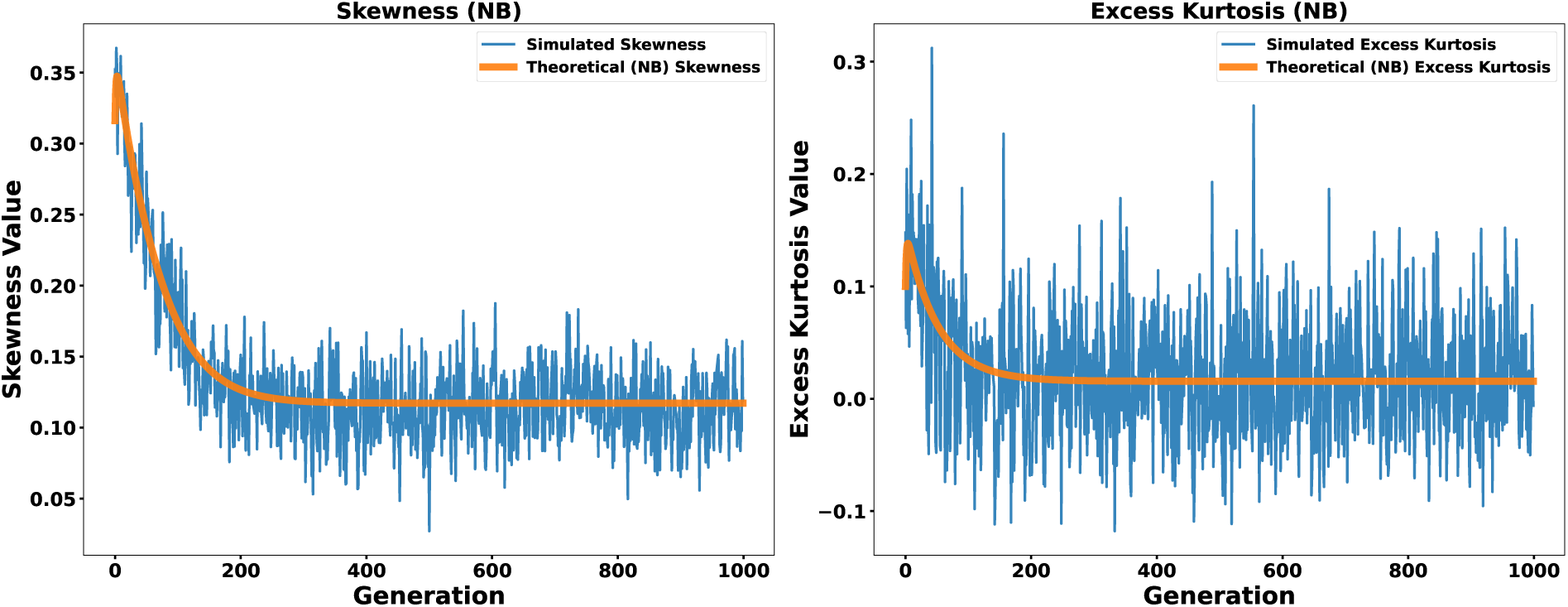
Comparison of theoretical (parametric closure) skewness and excess kurtosis dynamics of TE copy numbers to stochastic simulations, post-transposition/excision. The graph shows the parametric closure (straight lines) and stochastic simulation (wavy lines) for the skewness (left) and excess kurtosis (right) dynamics of active TE copy numbers over time using a Gaussian fitness function. Parameter values are as in Fig. 1. The resulting skewness and excess kurtosis values are given in Supplementary Table S2.

**Figure 3:**
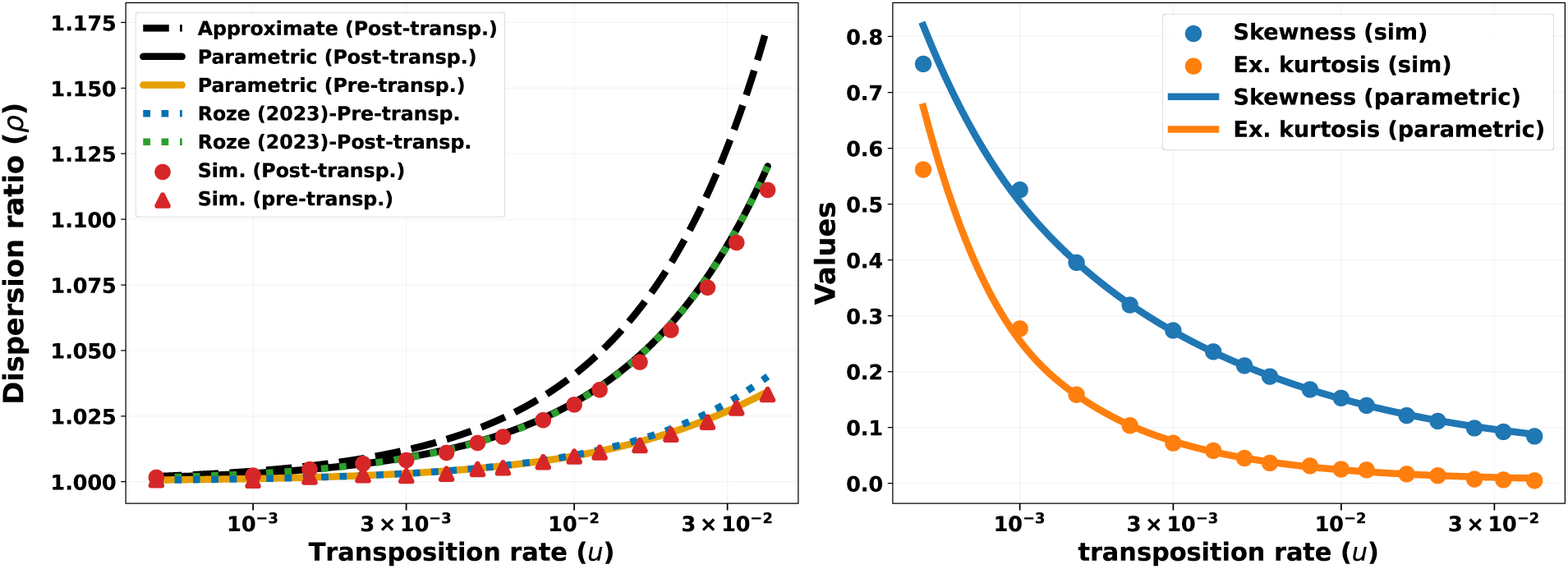
Comparison of theoretical predictions and stochastic simulations across transpo- sition rates. In the left panel, the dispersion parameter *ρ* is plotted as a function of the transposition rate *u* (on a log scale), comparing the approximate closure (black, dashed line), the parametric negativebinomial closure (black, solid line), the stochastic simulations (red points), and the prediction of Roze (2023) (purple line). In the right panel, skewness (blue line) and excess kurtosis (orange line) are shown for the parametric negative-binomial closure and compared to stochastic simulation results (points), at the post-transpositon/excision stage. Parameter values are *N* = 10^4^, *s* = 1*/N*, free (binomial) recombination, and *v* = 10*^−^*^4^, under the Gaussian fitness function, and *u* varying from 5 *×* 10*^−^*^4^ to 0.04, with *β* = 2*s* for Roze (2023). The predicted dispersion ratio is calculated from Eq. (4) for the approximate closure and from the closed-form equilibrium of the parametric closure, Eq. (14) post-transposition/excision and Eq. (23) pre-transposition/excision, with skewness and excess kurtosis from Eq. (16). Simulation results and the prediction from Roze (2023) are obtained as reported in the caption of Table 1.

**Table 1:** Comparison of stochastic simulation and theoretical values (post-transposition/excision stage). The table reports the mean, variance, dispersion parameter, skewness, and excess kurtosis, for transposition rates *u ∈ {*0.001, 0.005, 0.01, 0.016*}* and constant excision rate *v* = 10*^−^*^4^, as in Fig. 3 under the Gaussian fitness function. The stochastic model was simulated for 50,000 generations, and a single equilibrium value was obtained by averaging samples taken every 100 generations after a burn-in period of 9,000 generations. Theory (neg. bin.) denotes the parametric negative-binomial closure, evaluated from its closed-form equilibrium, Eq. (15) for the mean and variance and Eq. (14) for the dispersion ratio, with skewness and excess kurtosis from Eq. (16); the corresponding pre-transposition/excision values follow from Eq. (23). Theory (approx.) gives mean and variance from the approximate closure, Eq. (6); skewness and excess kurtosis are not reported because the approximate closure assumes the higher moments are zero. For Roze (2023), the mean follows from Eq. (11) in Roze (2023), and variance is derived by extending the framework of Roze (2023) through their supplementary material, the dispersion parameter *ρ* is then computed as the ratio of their variance to their mean; skewness and excess kurtosis are not reported. Supplementary Tables S5–S6 give the full set of post-transposition/excision equilibrium moment values across transposition rates, while the corresponding pre-transposition/excision values are given in Supplementary Tables S3–S4.

|  | u = 0.001 |  |  |  |  | u = 0.005 |  |  |  |  |
| --- | --- | --- | --- | --- | --- | --- | --- | --- | --- | --- |
| | Mean | Var. | $\rho$ | Skew. | Ex. kurt. | Mean | Var. | $\rho$ | Skew. | Ex. kurt. |
| Stochastic sim. | 4.06 | 4.07 | 1.0036 | 0.50 | 0.26 | 23.75 | 24.11 | 1.0152 | 0.21 | 0.04 |
| Theory (neg. bin.) | 3.98 | 4.00 | 1.0032 | 0.50 | 0.26 | 23.62 | 23.98 | 1.0152 | 0.21 | 0.05 |
| Theory (approx.) | 4.48 | 4.50 | 1.0040 | — | — | 24.01 | 24.50 | 1.0202 | — | — |
| Roze (2023) | 4.50 | 4.51 | 1.0031 | — | — | 24.50 | 24.87 | 1.0151 | — | — |

**Table 1: Comparison of stochastic simulation and theoretical values (post-transposition/excision stage).**
|  | u = 0.01 |  |  |  |  | u = 0.016 |  |  |  |  |
| --- | --- | --- | --- | --- | --- | --- | --- | --- | --- | --- |
| | Mean | Var. | $\rho$ | Skew. | Ex. kurt. | Mean | Var. | $\rho$ | Skew. | Ex. kurt. |
| Stochastic sim. | 47.80 | 49.25 | 1.0303 | 0.15 | 0.03 | 76.10 | 79.64 | 1.0465 | 0.12 | 0.02 |
| Theory (neg. bin.) | 47.52 | 48.95 | 1.0302 | 0.15 | 0.02 | 75.30 | 78.93 | 1.0482 | 0.12 | 0.02 |
| Theory (approx.) | 47.56 | 49.50 | 1.0408 | — | — | 74.57 | 79.50 | 1.0661 | — | — |
| Roze (2023) | 49.50 | 50.99 | 1.0301 | — | — | 79.50 | 83.32 | 1.0481 | — | — |

Nevertheless, Fig. 1a shows that the approximate mean does not fully capture the simulated mean trajectory, indicating a limitation of this closure. This mismatch is a consequence of the approximation introduced above, namely the neglect of the skewness and excess-kurtosis contributions in the variance dynamics, which effectively assumes an approximately normal TE copy-number distribution.

The simulations indicate that these higher-order moments, especially skewness, are not negligible, and suggest that a more accurate description requires a closure that retains information about the shape of the distribution beyond its mean and variance.

### 2.2 Parametric closure via the negative binomial distribution

To improve upon the earlier approximations of skewness and excess kurtosis, and to better capture the overall shape of the distribution; we adopt the negative binomial distribution as a parametric model. Note that this is an assumption to account for overdispersion, as is often done in statistical modeling. This choice of a negative binomial distribution takes advantage of the well-known relationships between mean, variance, skewness, and kurtosis in this family of distributions. By doing so, we can express the skewness and kurtosis in terms of the mean and variance, effectively closing the system of moment equations.

We obtain the parameters of the negative binomial distribution by moment matching. That is, we equate the mean and variance of the negative binomial with the mean *µ* and variance *σ*^2^ derived from our stochastic model. The dynamics for the mean, variance and higher moments, under this parameterization, are referred to as ‘nb’, which yields the following:

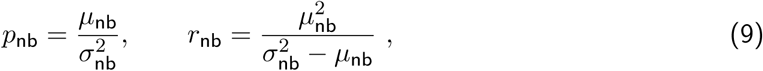

where *p* corresponds to the ‘success probability’ and *r* to the ‘number of successes’, the parameters of the negative binomial distribution. The inverse of this ‘success probability’ *p*_nb_ is equal to the dispersion ratio *ρ*_nb_, i.e., *ρ*_nb_ = 1*/p*_nb_ which we use as our dispersion measure. However, we note that the negative binomial distribution only allows us to study overdispersion and equidispersion. In contrast, underdispersion would correspond to *p >* 1 and *r <* 0, which are outside the valid parameter range. In the negative binomial distribution, the skew and excess kurtosis can then be written as

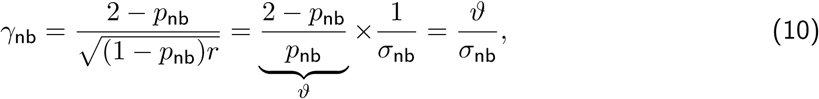

and

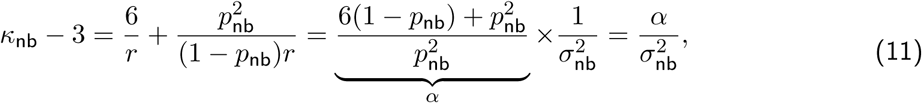

where *ϑ* and *α* are implicitly defined by the last respective equalities.

These parameterizations allow us to express the higher-order moments, skewness and excess kurtosis, in terms of the mean and variance, which results in a parametric moment closure of Eqs. (1) and (2):

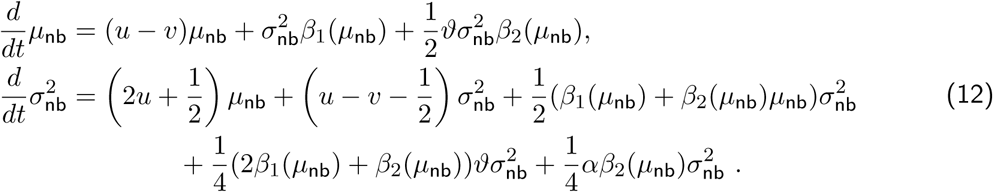

As before, it is not generally possible to solve the equilibrium conditions of these equations explicitly, as the parameters *β*_1_ and *β*_2_ may depend on the mean equilibrium copy number. Nevertheless, an equilibrium condition can be derived for the inverse dispersion ratio 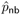 (see supplementary methods section S2.1 for the explicit expression). Moreover, the TE copy number reaches a positive equilibrium only when

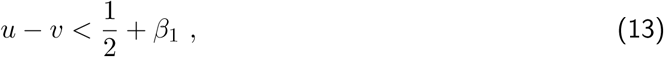

see Supplementary Eq. (S91) for the derivation. This is the counterpart of the previous condition on the net transposition rate being smaller than 1*/*2, obtained in the earlier approximation. Violation of this condition leads to unstable TE copy-number dynamics.

For the Gaussian fitness function *w*(*n*) = exp(*−sn*^2^), the equilibrium of Eq. (12) can be solved in closed form when *u ≪* 1 (see section S4 of the Supplementary Methods for the derivation). The dispersion ratio is then

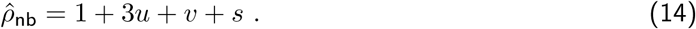

The dispersion ratio therefore depends on the selection coefficient, but only through the additive term *s*, so that the effect of selection is very small; this is consistent with the simulations across two orders of magnitude in *s* in Fig. 4, where *ρ* stays nearly constant. The term *s* enters solely through the skewness and excess kurtosis of the negative-binomial closure: removing their contributions from Eq. (12) leaves a dispersion ratio that is independent of *s* and recovers the approximate-closure result of Eq. (6) exactly. For *s, v ≪ u* this reduces further to 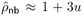, which coincides with the prediction extended from Roze (2023) at the same life-cycle stage, as we discuss in Section 2.4.

**Figure 4:**
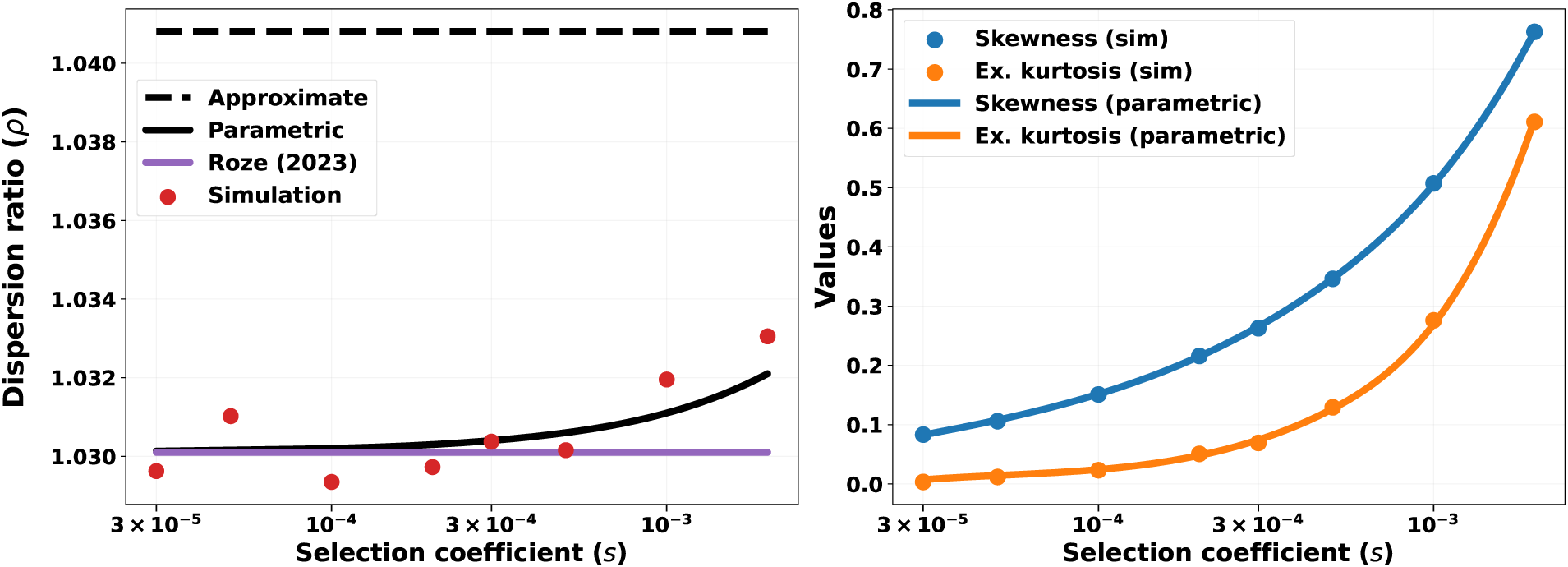
E**f**fect **of selection strength on TE copy-number distribution, post-transposition/excision.** Left panel: Dispersion ratio *ρ* as a function of selection coefficient *s* (log scale), comparing the approximate closure (black dashed line), the parametric negative-binomial closure (black solid line), the prediction of Roze (2023) (purple line), and stochastic simulations (red points). Right panel: Skewness (blue) and excess kurtosis (orange) from the parametric closure (lines) and stochastic simulations (points). Parameter values are *N* = 10^4^, *u* = 0.01, *v* = 10*^−^*^4^, free (binomial) recombination, Gaussian fitness *w*(*n*) = exp(*−sn*^2^), and *s* varying from 3 *×* 10*^−^*^5^ to 2 *×* 10*^−^*^3^, with *ρ* = 1 + 3*u* and *β* = 2*s* for Roze (2023). The parametric closure is evaluated from its closed-form equilibrium, Eqs. (14) and (16), and the approximate closure from Eq. (4); theoretical predictions are otherwise computed as in Fig. 3; see the caption of Table 1 for details, and the equilibrium moment values are given in Supplementary Table S9.

Neglecting the order-*s* term, the equilibrium mean and variance follow as

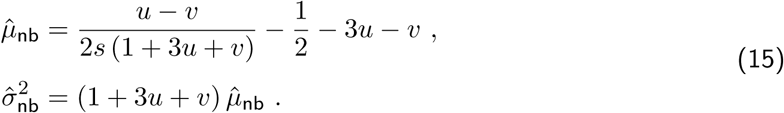

The corresponding higher moments are

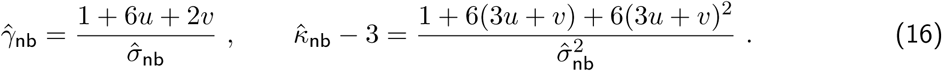

Since 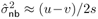, both moments are small and scale with the ratio of selection to transposition, 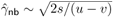 and 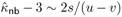, so that the copy-number distribution is right-skewed and heavy-tailed but approaches normality as selection weakens relative to transposition (see Fig. 4, right panel).

In Fig. 1b, we compare the equilibrium mean and variance under the parametric closure, Eq. (15), to simulated TE dynamics under the Gaussian fitness function. The parametric closure yields a slightly improved fit of mean and variance between the theory and simulations compared to the approximate moment closure, thereby correcting the inaccuracy introduced by neglecting skewness and excess kurtosis. It also predicts the mean more accurately than the classical expression of Charlesworth and Charlesworth (1983), Eq. (8), which tracks the simulated variance rather than the mean (Fig. 1b). This makes the negative-binomial closure the more accurate and informative approximation of the two. We additionally compared our theoretical skewness and excess kurtosis, computed from the analytic expressions in Eq. (16), with simulation results under the Gaussian fitness function (Fig. 2). The comparison with the simulation shows that the negative binomial parametrization is a good description of the TE copy number distribution even during the transient phase where the TE copy number has not yet equilibrated.

Beyond the fixed parameter set used so far (Figs. 1 and 2), in Fig. 3 we show results across a broader range of transposition rates while keeping the remaining parameters fixed. We observe that overdispersion (*ρ*) increases as transposition rate (*u*) increases, which is consistent with our theoretical expectation (Eqs. (4), (6), (14) and (S95)). Across these cases, all moments show good agreement with the stochastic simulations, demonstrating that the parametric closure via the negative binomial distribution provides a reliable approximation. Notably, we see that the parametric closure fits the simulated overdispersion values more closely than the approximate moment closure (left panel in Fig. 3). This is because the approximate closure neglects skewness and excess kurtosis in the mean-variance dynamics, whereas the parametric closure retains them through Eq. (12). The right panel of Fig. 3 shows that both the skewness and the excess kurtosis predicted by the negative binomial closure fit the simulation results well. Both quantities are positive, i.e., the TE copy number distribution is right-skewed and has positive excess kurtosis relative to the normal. Skewness is larger than excess kurtosis, so asymmetry is more pronounced than tail inflation in standardized terms. As transposition rate *u* increases, both quantities decrease, as predicted by Eq. (16): higher transposition raises the equilibrium variance, and both moments scale inversely with it. In particular, excess kurtosis becomes very close to zero at the highest transposition rates, while skewness stays small but positive. This is biologically relevant because it shows that increasing transposition alters not only the mean-variance relationship, but the entire shape of the TE copy-number distribution.

### 2.3 Pre-transposition/excision mean, variance, and dispersion ratio

So far, *µ* and *σ*^2^ describe the TE copy number distribution after a complete birth event, following parental sampling, recombination, and the subsequent transposition and excision events. It is also useful to compute the mean, variance, and dispersion ratio of the offspring’s TE copy number immediately after parental sampling and recombination, but before transposition and excision occur. At this earlier stage, the offspring carries a copy number that we denote *m^′^*; because *m^′^* is drawn from the same fitness-weighted parents that underlie Eqs. (1)–(2), its mean and variance follow from the same derivation by evaluating the process one step earlier in the life cycle, prior to transposition and excision (see section S5 of the Supplementary Methods for the full derivation):

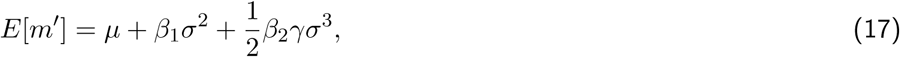

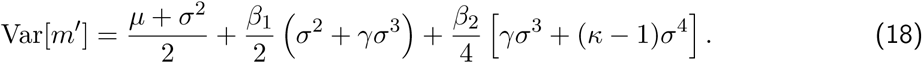

Specializing Eqs. (17)–(18) to the negative-binomial closure, i.e., substituting *γ*_nb_ = *ϑ/σ*_nb_ and 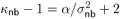 from Eqs. (10)–(11), gives

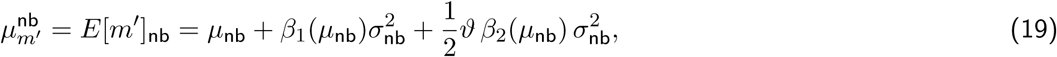

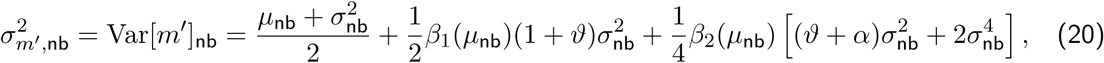

and the higher moments are given as

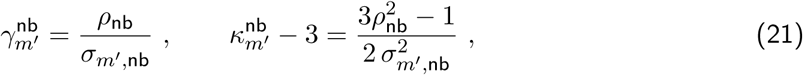

where *µ*_nb_ and 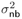 are the same equilibrium moments obtained from the parametric closure, Eq. (12). The same simplification carries through for the approximate closure, i.e., setting *γ* = 0 and *κ* = 3 in Eqs. (17)–(18); we give the resulting expressions in section S5.2 of the Supplementary Methods.

The resulting pre-transposition/excision dispersion ratio,

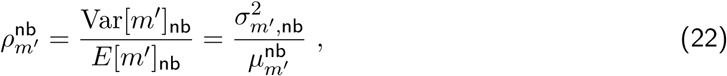

describes the same quantity as *ρ* in Eqs. (1)–(2) and (12), but measured one stage earlier in the life cycle, directly after recombination and before transposition and excision.

For the Gaussian fitness function *w*(*n*) = exp(*−sn*^2^), Eqs. (19)–(22) can again be evaluated in closed form at equilibrium (see section S5.3 of the Supplementary Methods), giving

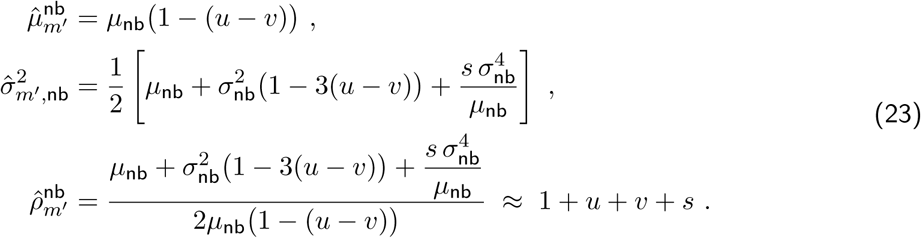

The higher moments are

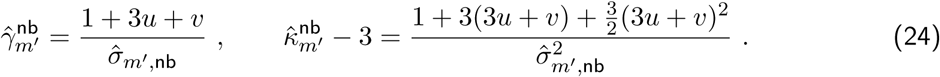

For *v, s ≪ u* the dispersion ratio is therefore approximately 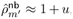, in contrast to 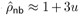 one step later in the life cycle. This turns out to matter for a fair comparison with Roze (2023), whose theory measures overdispersion at exactly this pre-transposition/excision stage, as we discuss next. We include 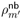 alongside our post-transposition predictions in Fig. 3 and Table 1.

### 2.4 Comparison to Roze (2023)

We further compare our predictions with those of Roze (2023), who derived the dispersion ratio, *ρ*, from linkage disequilibrium between TE insertions (Eq. (6) in Roze (2023)). Their result applies immediately after recombination, before transposition and excision, corresponding to the same life-cycle stage as our pre-transposition/excision quantity, *ρ_m_′* (Section 2.3). Under free recombination (with *v ≪ u*), their expression simplifies to *ρ ≈* 1 + *u*, which is exactly our 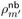 in Eq. (23). For the later life-cycle stage, extending the framework of Roze (2023) through their supplementary material gives *ρ ≈* 1 + 3*u*, matching our post-transposition/excision dispersion ratio in Eq. (14). As shown in Fig. 3 and Table 1, both theoretical predictions closely match the overdispersion observed in the simulations at their corresponding stage of the life cycle.

Although Roze (2023) accurately predict the overdispersion, their predictions for the mean and variance differ slightly from our simulations. This difference arises because they take the equilibrium mean from Charlesworth and Charlesworth (1983), which assumes a Poisson-distributed TE copy number, and derive the corresponding variance (Eq. (5) in Roze (2023)) under the same framework. As a result, both the mean and variance are slightly overestimated relative to our simulations (Table 1; Supplementary Tables S3–S4). Because these quantities are inflated by a similar proportion, the bias largely cancels when taking their ratio. The inflation lies in the mean: (*u − v*)*/*2*s* is in fact the equilibrium variance rather than the mean, and the mean is smaller than it by the factor 1*/*(1+3*u*+*v*) (Eq. (15)).

Taken together, these comparisons show that overdispersion is not a fixed property of the population, but depends on the stage of the life cycle at which it is measured, with the pre- and post-transposition/excision dispersion ratios differing roughly three times in their departure from the Poisson expectation, *u* against 3*u* (Fig. 3, Table 1).

### 2.5 Effect of selection strength on TE copy number distribution

Having examined how transposition rate affects overdispersion, we now investigate how varying selection strength *s* influences the shape of the TE copy-number distribution. We vary the selection coefficient *s* from 3 *×* 10*^−^*^5^ to 2 *×* 10*^−^*^3^ while holding the transposition rate fixed at *u* = 0.01 and the excision rate at *v* = 10*^−^*^4^ (Fig. 4); beyond *s ≈* 2 *×* 10*^−^*^3^, TE copy numbers become too small for meaningful comparison.

Increasing selection strength substantially reduces both the mean and variance of TE copy number (see Supplementary Table S9). As *s* increases across the range tested, the mean TE load declines from approximately 159 copies to about 2 copies, reflecting more efficient purging of deleterious TE insertions under stronger selection. Despite this large reduction in TE abundance, the dispersion ratio *ρ* remains relatively stable, showing only a slight increase beyond *s ≈* 3 *×* 10*^−^*^4^ (Fig. 4, left panel). This suggests that while selection modulates the absolute level of TE load, it does not strongly influence the overdispersed nature of the copy-number distribution.

In contrast to the dispersion ratio, the higher moments of the distribution respond more strongly to selection strength. The simulations reveal a clear increase in both skewness and excess kurtosis as selection becomes stronger (Fig. 4, right panel). Under weak selection, the TE copy-number distribution is only mildly right-skewed (*γ ≈* 0.06), but as selection increases, the distribution becomes increasingly right-skewed and heavy-tailed. Biologically, this pattern reflects the concentration of most genomes at low TE copy numbers under strong purifying selection, while a small fraction of genomes experiences (stochastic) TE bursts, producing rare high-copy-number outliers.

Importantly, the simulated skewness and excess kurtosis closely match the predictions of the negative-binomial (NB) parametric closure across all selection regimes (Fig. 4, right panel), confirming that this closure provides an accurate description of the stationary TE copy-number distribution even as selection varies over two orders of magnitude. This agreement demonstrates that our model captures not only the mean-variance relationship but also the full shape of the distribution under biologically relevant conditions.

### 2.6 Conditions for overdispersion, underdispersion and the Poisson case

In Figs. 3 and 4, we see that TE copy numbers are always overdispersed in simulations for both life-cycle stage and for our choice of parameters and fitness functions. We now identify general conditions for overdispersion (*ρ >* 1). For the post-transposition stage, we start by assuming that the TE copy numbers are Poisson distributed in equilibrium. From this, we then show that this results in a contradiction on the parameter constraints, thus showing that the equilibrium distribution is not a Poisson distribution.

We assume that in equilibrium we have *ρ* = 1. This condition is equivalent to

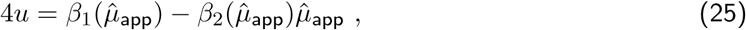

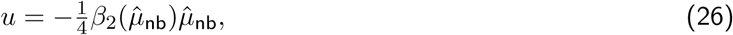

for the approximate (from Eq. (4)) and parametric closures (Eq. (S94)), respectively. Taking the example of a Gaussian fitness function (exp(*−sn^γ^*); *γ* = 2), the approximate closure directly yields a contradiction because the right-hand side in Eq. (25) is equal to zero. Therefore, we would have the condition that *u* = 0, and thus the TE would not invade in the first place. In the parametric closure, the condition in Eq. (26) can be rewritten using Eq. (12) (note that *ϑ* = 1 in the case of *ρ* = 1), which results in the condition

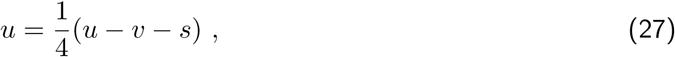

which again yields a contradiction. More general (and abstract) conditions for arbitrary fitness functions are studied in Section S3 in the Supplement. These can be used to infer whether for a specific fitness function, equidispersion in equilibrium is a possibility.

It is more involved to formally prove that underdispersion is impossible in equilibrium in our model, because the parametric closure via the negative binomial distribution is only defined for *ρ ≥* 1. We therefore test for underdispersion numerically, across a range of *γ* values in the Gaussian fitness function, and find none at this stage (Appendix E, right panel of Fig. E.1). We further provide a condition for the less accurate approximate closure in Section S3.2 in the Supplementary Information. Since neither the closure nor a finite set of simulations can establish impossibility, we conjecture that for active TEs, i.e. with *u >* 0, underdispersion in equilibrium is not possible under free recombination for the post-transposition/excision stage. However, in the pre-transposition/excision stage (*m^′^*, the copy number immediately after recombination but before transposition and excision occur; Section 2.3), all three outcomes remain possible, governed by the curvature of the log-fitness function, *β*_2_. Weak curvature, produced by a fitness function that declines only gradually with copy number (small *γ*, e.g., *γ* = 1.5), leaves the small restoring pull toward *ρ_m_′* = 1 in control, so *m^′^* stays overdispersed; sufficiently strong, concave curvature, produced instead by a steep, threshold-like decline (large *γ*, e.g., *γ* = 4; compare the fitness curves in the left panel of Fig. E.1), overwhelms this pull and drives *m^′^* below one, with equidispersion marking the exact boundary between the two regimes (Supplementary Section S5.4). We confirm both over- and underdispersion through simulations spanning this range of *γ* values for the Gaussian fitness function (Appendix E).

Taken together, these results indicate that whether TE copy numbers can be equidispersed or underdispersed depends on the life-cycle stage considered: both remain possible before transposition and excision take place, but neither is expected to persist once transposition and excision have occurred.

### 2.7 Model comparison with data

We now qualitatively compare the theoretical prediction mean, variance, skewness, and excess kurtosis to empirical TE data using the preprocessing described in subsection 4.3. Figure 5 summarizes the dispersion of per-strain TE copy numbers of 164 *D. melanogaster* strains from a Zambian population (Lee, 2022) in two complementary views. The TE counts are restricted to the euchromatic part of the genome, where recombination rates are higher than in the remaining heterochromatic region. In the left panel of Fig. 6, sample mean and variance are shown for each TE family and in the right panel, the same TE families are ordered by the dispersion ratio *ρ* = variance*/*mean; both showing that most families lie above the Poisson diagonal (variance = mean), thus being overdispersed with all the LTR families being overdispersed and ROO being the most overdispersed family. Interestingly, ROO is considered to be one of the most active TE families in *D. melanogaster* (Díaz-González and Domínguez, 2020). Moreover, four TE families are underdispersed (*ρ <* 1): BS, 1360, S, and TRANSIB2. This could reflect low or absent transposition for some families. For example, prior work on *D. melanogaster* associates BS, S, and TRANSIB2 with transpositional inactivity or near inactivity (Kapitonov and Jurka, 2003; Maside et al., 2003; Udomkit et al., 1995). In general, this pattern that the transposition rate positively correlates with the dispersion ratio is consistent with our theoretical analysis of overdispersion of TE copy-numbers under transposition-excision-selection TE processes in highly recombining regions.

**Figure 5:**
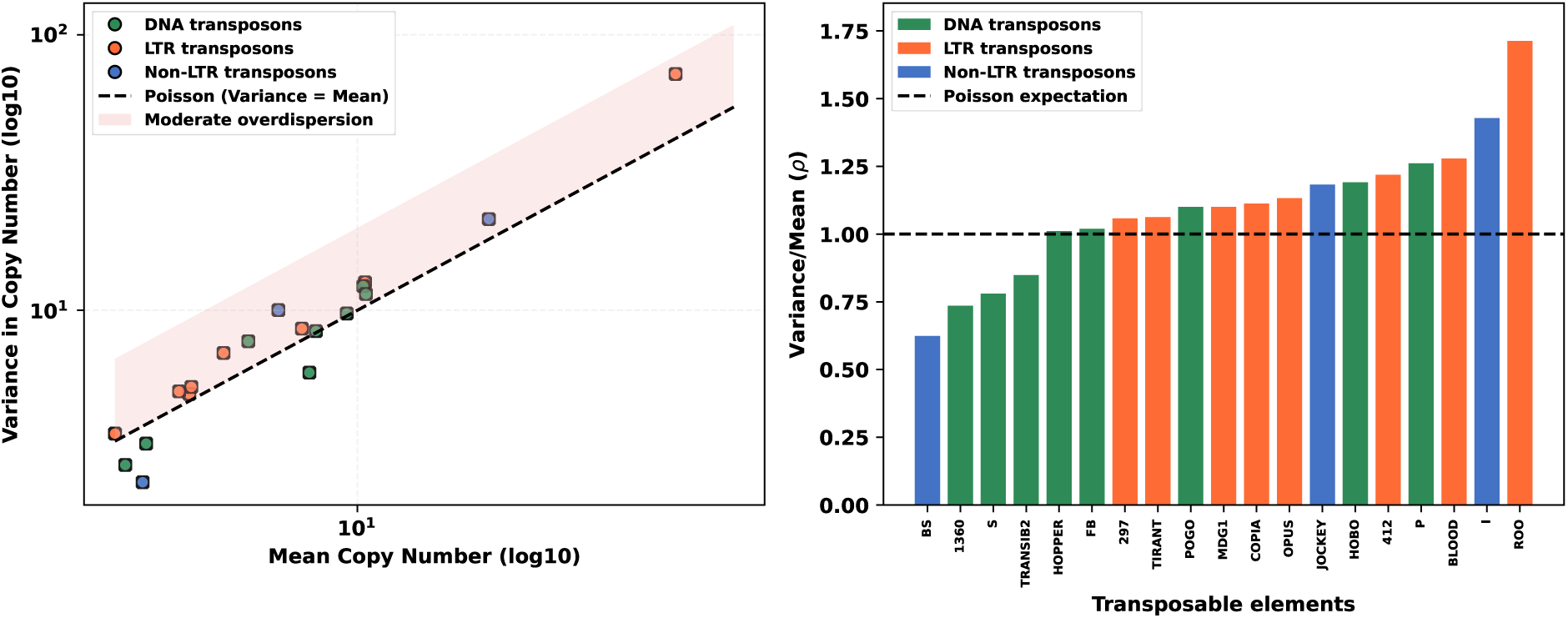
M**e**an**–variance relationship and dispersion across TE families. Left:** Sample mean and variance of per-strain TE burdens (sums of presence calls across sites) for each family (dots) passing the filters described in subsection 4.3, coloured by broad TE order (DNA, LTR, non-LTR). The dashed line is the Poisson prediction variance = mean; the shaded band marks our “moderate overdispersion” range, where the dispersion ratio *ρ* = variance*/*mean exceeds 1 but is not greater than 2. **Right:** Dispersion ratio *ρ* = variance*/*mean for the same families, ordered by *ρ*; the horizontal dashed line is the Poisson expectation *ρ* = 1.

**Figure 6:**
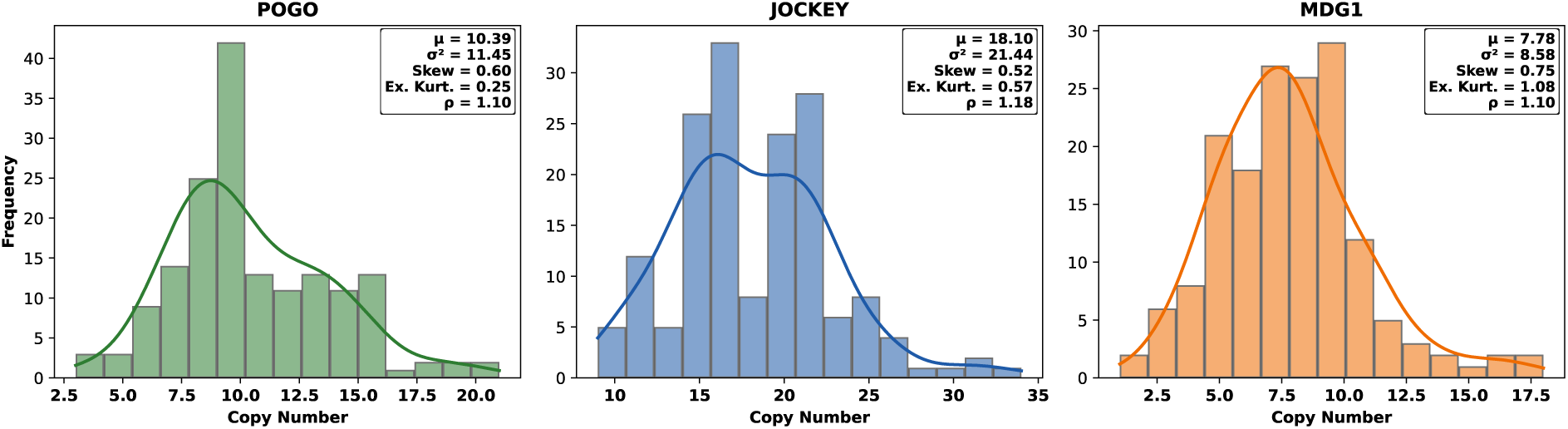
D**i**stribution **of per-strain TE burdens.** Histograms for three representative families from the filtered set of 19, one from each TE class: DNA (green), non-LTR (blue), and LTR (orange). Overlaid smooth curves are Gaussian kernel density estimates of the same per-strain copy numbers, with bin width and density scaled to the histogram frequency scale. In-panel summaries give sample mean, variance, skewness, excess kurtosis, and *ρ* = variance*/*mean; all three are overdispersed (*ρ >* 1) and right-skewed. Plots for the remaining filtered families are in Figure S1 in the Supplementary Material.

Figure 6 summarizes per-strain TE burden distributions for three representative families, one from each TE class (see Table 3 in Appendix C for the representative elements calculated moments and statistics); the distributions for all 19 families are shown in Supplementary Figure S1, and the corresponding moments and statistics are shown in Supplementary Table S1. Across all families, skewness and excess kurtosis are not always positive, but most families show positive values for both, which is consistent with our prediction from the negative-binomial moment closure, which implicitly assumes overdispersion (Eqs. (10)–(11)). For the majority of families, the empirical magnitudes of skewness and excess kurtosis lie in the range of values traced by the parametric closure and simulations in the right-hand panel of Fig. 3 (across the transposition rates shown there). ROO is an exception: its excess kurtosis is somewhat higher than that typical range. Overall, these observed TE burden distributions qualitatively are consistent with the distribution predicted by our parametric closure.

## 3 Discussion

Natural populations of active TEs typically show overdispersed copy-number distributions (variance exceeding the mean) and right-skewed burdens (Lee, 2022; McGurk et al., 2021; Said et al., 2022). In the classical free-recombination theory of Charlesworth and Charlesworth (1983), TE copy number is treated as Poisson-like (variance *≈* mean). As a result, the predicted mean (or variance) can be inaccurate, which has been noted before when theory was compared to simulations (Huang and Lee, 2024; Tomar et al., 2023). To address this problem, we derive mean and variance dynamics under free recombination directly from first principles, using a diploid biparental Moran process with transposition, excision, and selection (Eqs. (1)–(2)), that captures the distributional properties of TEs and is qualitatively consistent with empirical observations.

Our model goes beyond mean and variance to describe the shape of TE distributions through skewness (asymmetry) and excess kurtosis (tail heaviness) (Eqs. (10), (11)), which emerge from the underlying biological processes. Positive skewness within a population reflects ongoing transposition in our model; the variance in copy number scales (among other terms) with the transposition rate times copy number, so fluctuations above the equilibrium tend to be larger than below it (Eq. (12)), with most individuals with low/moderate TE load and a few individuals with high TE load, contributing to the characteristic right-skew seen in data (McGurk et al., 2021). Moreover, the right panel of Fig. 3 shows that as transposition increases, both skewness and excess kurtosis decrease; this means higher transposition raises TE load more broadly across individuals, so the distribution becomes less dominated by a few extreme high-copy genomes, even though it remains overdispersed. Excess kurtosis is typically small and near zero in our simulations, so extremely high copy numbers are not strongly overrepresented, consistent with purifying selection against very large TE loads (Lee, 2022).

We employed two approaches to close the moment equations: an approximate closure that neglects higher-order moments, and a parametric closure using the negative binomial distribution. Compared with simulations, Fig. 1a shows that the approximate closure captures the variance better than the mean, while the negative-binomial closure matches both mean and variance well (Fig. 1b). The same pattern appears in the left panel of Fig. 3; across transposition rates, the negative-binomial predictions remain close to the simulated dispersion ratios, while the approximate closure deviates more, especially at higher transposition rates. Overall, the negative-binomial closure is more accurate because the approximate closure neglects higher-order moment effects.

At equilibrium, we derive expressions for the dispersion ratio *ρ*, which quantifies how variance relates to mean copy number and frames the biological conditions for overdispersion. This ratio depends on exactly when in the life cycle it is measured, we therefore distinguish two stages throughout: immediately after recombination but before the current generation’s transposition and excision act (*ρ_m_′*, Section 2.3), and after a complete birth event (*ρ*, used elsewhere in this paper); Fig. 3 reports both together. Based on our model, overdispersion emerges when transposition exceeds selection strength multiplied by the average number of TE copies per individual to some power, allowing individual variation in TE activity to generate the observed distributional patterns (McGurk et al., 2021; Said et al., 2022). The underlying biological mechanism behind this condition is that transposition generates positive linkage disequilibrium (LD) between TE insertion sites (Roze, 2023), leading to the association between TE insertion sites. Recombination however, works in the opposite direction, reshuffling insertions between homologous chromosomes each generation and thereby eroding the very associations transposition builds up. Under free recombination this erosion is strong but not total: some LD from previous generations survives it, which is why *ρ_m_′*, measured just after recombination, already exceeds one; transposition then lays down fresh LD on top of this before the next round of recombination can act on it, which is why *ρ*, measured after a complete birth event, is noticeably larger still (Fig. 3). Thus, the level of overdispersion observed under free recombination depends on the positive LD among TE insertions generated by the transposition process.

Moreover, our analysis describes the effect of purifying selection on dispersion: stronger selection reduces mean TE load but the dispersion ratio remains relatively stable (Fig. 4, left panel). In contrast, stronger selection increases both skewness and excess kurtosis (Fig. 4, right panel), reflecting that most genomes are concentrated at low copy numbers while a few individuals still harbour transient TE amplifications. In addition, we show that higher transposition rates result in larger overdispersion (left panel of Fig. 3), which is consistent with observations in Tomar et al. (2023), where high transposition invalidates the Poisson approximation in the classical model.

Additionally, for the mean TE copy number to remain positive and thus sustain overdispersion, the net transposition rate must remain below roughly 0.5 insertions per copy per generation. This boundary represents a co-evolutionary constraint between hosts and TEs: a TE cannot be too ‘selfish,’ because if each copy produces more than half a surviving new copy per generation, this is likely to result in genome instability and therefore strong purifying selection. Empirically, TE transposition rates are well below this threshold (10*^−^*^5^ to 10*^−^*^2^ per copy per generation) (Ewing and Kazazian, 2010), consistent with the view that natural populations maintain TE activity within a range that avoids destabilizing the genome (Bourque et al., 2018; McGurk et al., 2021).

We further compare our overdispersion results with those of Roze (2023), who derived the dispersion ratio *ρ* from the linkage disequilibrium that transposition generates among TE insertions. Our analysis refines this comparison in the high-recombination regime (see Fig. D.1 with *R ≥* 1) by recognizing that *ρ* is not a fixed property of the population but depends on the stage of the life cycle at which it is measured. Roze (2023)’s original prediction applies immediately after recombination, before the current generation’s transposition and excision events, and matches our pre-transposition/excision ratio *ρ_m_′* closely, at leading order *ρ_m_′ ≈* 1 + *u*, as in Eq. (23). Extending their framework to the post-transposition/excision stage later in the life cycle instead gives *ρ≈* 1+3*u* (Denis Roze, personal communication), which matches our post-transposition/excision ratio *ρ* in Eq. (14) equally well (left panel of Fig. 3, Table 1).

This agreement in the dispersion ratio holds even though the individual moments do not match as closely. Because Roze (2023) take the equilibrium mean from Charlesworth and Charlesworth (1983), which assumes Poisson-distributed copy numbers, their mean and variance both sit slightly above the simulated values (Section 2.4, Table 1). What our closed form adds is an interpretation of that classical expression: once copy numbers are overdispersed, (*u − v*)*/*2*s* gives the equilibrium variance rather than the mean, and the mean is smaller by the factor 1*/*(1 + 3*u* + *v*) (Eq. (15)). Because the offset affects their mean and variance alike, it leaves their dispersion ratio intact, and the agreement between their prediction and ours is unaffected by the discrepancy in the moments themselves.

The same pattern appears more visibly, in the low-recombination regime (*R ∈* (10*^−^*^5^, 1] in Fig. D.1) considered by Roze (2023), although this regime is not the primary focus of our study. The gap between their moments and the simulations grows wider there, because a second problem is added to the one above: their mean assumes linkage equilibrium, following Charlesworth and Charlesworth (1983), while their variance includes linkage disequilibrium, and these two assumptions pull further apart as linkage becomes tighter (see Supplementary Tables S7–S8). The dispersion ratio is again unaffected, for the same reason as before: Eq. (14) in Roze (2023) does not carry the error in the mean, and it still follows the simulations closely in this regime (Fig. D.1).

We show in Section 2.6 that the possibility of equidispersion or underdispersion depends on where in the TE life cycle they are measured. At the post-transposition/excision stage, our theory rules out both as viable equilibria for actively transposing TEs: either would require the transposition rate to sit at a precise, unstable balance with selection, a balance our simulations of active TEs never sustain, so overdispersion emerges as the expected outcome whenever a stable TE equilibrium exists. Earlier in the life cycle, however, at the pre-transposition/excision stage, this restriction relaxes: which outcome arises instead depends on the curvature of the fitness function, *β*_2_, with steep, threshold-like declines in fitness favoring underdispersion over the milder, gradually declining fitness functions that favor overdispersion, a prediction we confirm both analytically (Supplementary Section S5.4) and by simulation (Appendix E). This stage-dependence suggests that the underdispersed or Poisson-like copy-number patterns seen in high-recombination regions more likely reflect inactive or low-activity TE families, in which deletion and drift dominate over transposition (McGurk et al., 2021), than a genuine pre-transposition signature; capturing such patterns explicitly would require extending our model to inactive TEs. In low-recombination regions, by contrast, Roze (2023) shows that even active TE copies can become underdispersed at the pre-transposition/excision stage, once the negative linkage disequilibrium generated by selection and drift (the Hill–Robertson effect) overtakes the positive disequilibrium generated by transposition. The same effect drives insertions to fixation through Muller’s ratchet in Dolgin and Charlesworth (2008): a site fixed in the population is carried by every individual and so contributes nothing to the variance in copy number, again pulling the variance below the mean. There, copy numbers are censused one stage later, after transposition and excision.

We compare our theory with TE burden data from the DPGP3 Zambian *D. melanogaster* dataset (Lee, 2022). The TE insertions used here lie in high-recombination euchromatin regions of the genome. After filtering strains by call rate and retaining TE families with at least 500 total presence calls, the 19 retained TE families across 164 *D. melanogaster* strains are mostly overdispersed, and many also show the right-skewness and positive excess kurtosis expected from the negative-binomial closure. In particular, all the LTR families are overdispersed; ROO is the most overdispersed family, which is consistent with its high transpositional activity and abundance in *D. melanogaster* (Díaz-González and Domínguez, 2020; de la Chaux and Wagner, 2009; García Guerreiro, 2012). At the same time, four families, BS, 1360, S, and TRANSIB2, are underdispersed, which may reflect low or absent transposition; in *D. melanogaster*, BS, S, and TRANSIB2 have been linked to lineages with little or no ongoing transposition (Kapitonov and Jurka, 2003; Maside et al., 2003; Udomkit et al., 1995). This supports the interpretation developed above that underdispersed or near-Poisson families may often be inactive or weakly active rather than representative of the active-transposition equilibrium described by our model. Overall, this single-population comparison provides qualitative support for the model, that active TEs (i.e., those undergoing transposition) should display an overdispersed copy number distribution, but it is not yet a quantitative test across all genomic contexts: uniform selection and free recombination likely underestimate the additional variability introduced by insertion bias, heterogeneity in selective effects, and recombination-rate variation across the genome (Langmüller et al., 2023, 2025; Roze, 2023).

More generally, there are various other factors that may change the pattern of overdispersion of TE copy numbers in populations. On the population level, spatial structure tends to increase overdispersion (McGurk et al., 2021). Within the genome, various host regulation processes such as piRNA clusters (Kofler, 2019; Tomar et al., 2023) control TE transposition, thus reducing the ‘effective’ transposition rate in equilibrium within the population, which we predict to lower the dispersion ratio, potentially even resulting in underdispersion, even under free recombination. Lastly, interaction of multiple TE families with a genome, e.g. autonomous-nonautonomous TE dynamics (Le Rouzic and Capy, 2006a; Omole and Czuppon, 2025), may also interfere with our prediction. Again, if this interaction results in a reduction of the transposition rate in equilibrium, we would expect a reduction in the dispersal ratio.

Future extensions of this study could therefore incorporate TE inactivation, e.g. through host regulation, variable selection strengths between insertion sites, and TE insertion site biases to capture the full spectrum of dispersion patterns observed across TE families (Langmüller et al., 2025; Tomar et al., 2023; Wierzbicki and Kofler, 2023). Such developments would further bridge the gap between theoretical predictions and the complex reality of TE evolution in natural genomes.

## 4 Model and methods

We develop a stochastic population genetics model to describe the dynamics of TEs in large diploid populations. Our framework extends the biparental Moran model of TE copy number evolution with free recombination introduced in Pfaffelhuber and Wakolbinger (2023), incorporating additional evolutionary forces relevant to TE evolution: purifying selection against deleterious insertions, transposition events that create new copies, and deletion events capturing the loss of TEs due to excision or genomic deletions. By explicitly modeling these processes, we can analytically derive the mean and variance dynamics of the TE copy number and examine how these forces interact to shape the TE copy number distribution within a population.

### 4.1 Moran model with transposable elements

We consider a diploid population of fixed size *N* evolving under a continuous-time Moran model with biparental reproduction. At each reproduction event, two individuals are chosen with probabilities proportional to their fitness values *w*(*l*) and *w*(*n*) (fertility selection), where *l* and *n* denote the TE copy numbers in their diploid genomes. The waiting time between events is exponentially distributed, and the rate at which events occur is 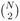, corresponding to the number of distinct unordered pairs in the population; for large *N*, this is approximately *N* ^2^*/*2 (Etheridge, 2011).

Following the biparental Moran framework of Pfaffelhuber and Wakolbinger (2023), the TEs of both parents are pooled and transmitted to the offspring under free recombination, i.e., each TE copy is inherited independently with probability 1*/*2. After recombination, the inherited number of TE copies, denoted *m^′^*, may change due to transposition or deletion events. We assume their rates are low enough that the probability of more than one event occurring in a single time step is negligible. With probability 1 *−* (*u* + *v*)*m^′^*, no event occurs and the copy number remains *m* = *m^′^*. A transposition event increases the TE count by one (*m* = *m^′^* + 1) and occurs with probability *um^′^*. Conversely, a deletion event decreases the count by one (*m* = *m^′^ −* 1) and occurs with probability *vm^′^*. Once the final TE count *m* is determined, the offspring replaces a random individual in the population. We measure overdispersion at two stages of this life cycle: immediately after recombination but before transposition and excision, where the copy number is *m^′^*, and after the complete birth event, where it is *m*. The mathematical formulation of both stages is presented in sections S1 and S5 of the supplementary material.

### 4.2 Stochastic simulation model

We implemented an individual-based forward-time Wright-Fisher simulation in a finite diploid population. We use an infinite-insertion-sites representation of the genome in which each diploid individual carries two homologous chromosomes stored as vectors of TE insertion positions on the interval [0, 1]. At initialization, the total number of TE copies per diploid is drawn from a Poisson distribution with mean *n*_init_ = 10, and each TE copy is assigned to one of the two homologs with equal probability and a uniformly random position in [0, 1]. In each generation, parents are sampled with probability proportional to fitness under the Gaussian function *w*(*n*) = exp(*−sn*^2^), where *n* is total diploid TE copy number.

For each selected parent, meiosis generates two gametes. In the free-recombination setting used for the main theory comparison, each TE copy is independently assigned to one gamete or the other (binomial assortment), matching the independent-inheritance assumption of our Moran model. We also implement a crossover-based mode, with a Poisson number of crossovers along a map of length *R*, for the purpose of comparison to other models. Offspring are formed by randomly choosing one gamete from each parent. TE content in the offspring is then updated by excision and transposition. Both events are modeled by binomial sampling at the copy level: each existing copy can transpose with probability *u*, and each existing copy can be excised with probability *v*. New insertions are placed uniformly at random positions in [0, 1] on randomly chosen homologs, and excision removes copies uniformly at random across those currently present.

The simulation tracks population statistics, including the mean, variance, dispersion ratio, skewness, and excess kurtosis of the TE copy number across multiple generations. To compare these results with theoretical predictions, the timescales of the simulated Wright-Fisher and analyzed Moran models must be aligned; specifically, *N/*2 Moran steps are treated as equivalent to one Wright-Fisher generation, which corresponds to the classical rescaling (Etheridge, 2011).

Table 2 summarises the simulation parameters. The simulations were implemented in Julia, while data analysis was performed using Python. All code and data files are publicly available at https://github.com/Ade-omole/TE_overdispersion.

**Table 2:** Overview of parameters. We summarize all parameter abbreviations, their biological interpretations, and default values used in the simulations and notation throughout the paper.

| Parameter | Description | Value (range) |
| --- | --- | --- |
| $N$ | population size (diploids) | $10^4$ |
| $s$ | selection coefficient | $1/N$ |
| $r / R$ | recombination setting (main results use binomial assortment of TE copies; crossover mode uses map length $R$) | binomial assortment; $R \in [10^{-5}, 10^3]$ |
| $u$ | transposition rate | $[5 \times 10^{-4}, 0.04]$ |
| $v$ | excision rate | $[10^{-4}, 0.006]$ |

### 4.3 Empirical TE data

To compare qualitatively the moments of our theoretical predictions; mean, variance, skewness, and excess kurtosis to data, we build per-strain TE *burden* distributions from the raw DPGP3 (*Drosophila* Population Genomics Project) presence–absence matrix for TE insertions in Zambian *D. melanogaster* strains (Lee, 2022). The raw data provided on https://github.com/jumpingTE-LeeLab/TE_insertion_DPGP3_raw shows TE insertions in euchromatic regions of the genome before the data manipulation explained in Lee (2022).

For our purposes, we removed the ambiguously assigned TEs (family equal to `multi`), since we are interested in family-specific burden distributions, which are the quantities that we compare to our theory. We then retain only strains that pass a minimum *call rate* threshold. For each strain, every site is either *callable* (assigned a definite `0` or `1`) or *uncallable* (coded `NA`), i.e., insertions that could not be reliably detected (called) at a genomic site. The *call rate* is therefore the fraction of sites that are not NA:

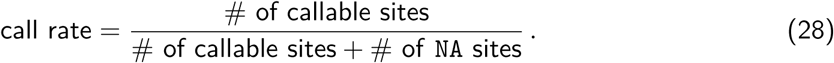

We set the minimum call rate to 0.8 and discard strains that fall below this threshold. Strains with many missing (NA) calls contribute fewer callable sites to each per-family burden sum, because NAs are omitted rather than counted as absences; mixing those strains with strains that are nearly fully called would compare burdens built from different total numbers of sites and could inflate among-strain variance in burdens for technical reasons (different effective numbers of loci summed) rather than reflect biological heterogeneity in TE load alone. Once strains are filtered in this way, we retain only TE families with at least 500 total presence calls summed across all sites and strains (*n*_present_ *≥* 500), because below that threshold families are too sparse for stable summaries: mean burdens lie near zero, and the dispersion ratio *ρ* = *σ*^2^*/µ* together with skewness and excess kurtosis are dominated by sampling noise rather than by interpretable among-strain variation. These filters leave 164 strains and 19 TE families, which we compare with our theoretical predictions; details of the retained TE families, their classes, and their corresponding statistical moments and dispersion ratios are listed in Supplementary Table S1.

## Data availability

The empirical TE dataset analyzed in this study is based on the DPGP3 *D. melanogaster* dataset of Lee (2022). In addition to the published dataset, the full dataset used for this reanalysis is included in our public repository, together with the simulated data files and the figures supporting the findings of this study, at https://github.com/Ade-omole/TE_overdispersion.

## Acknowledgement

AO acknowledges funding from the FlyInnovation project (CZ 294/1-1) within the GEvol priority program (SPP 2349), funded by the German Research Foundation (DFG). We thank Grace Yuh Chwen Lee for making the full DPGP3 dataset available for this reanalysis and for sharing her insights on TEs with us. Lastly, we are also grateful to an anonymous reviewer and Denis Roze for their constructive comments and suggestions, which substantially improved the manuscript. Specifically, Denis Roze pointed out the life-stage dependence of the dispersion ratio and provided the 1 + 3*u* approximation of the dispersion ratio derived in his modeling framework for meaningful comparison. We used generative AI tools for debugging simulation code and for improving the clarity and readability of the text. All derivations and analyses were carried out and independently checked by the authors.

## Appendix

## A Approximate dispersion ratio *ρ*_app_

At equilibrium, Eq. (3) becomes:

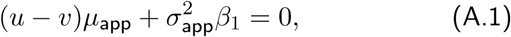

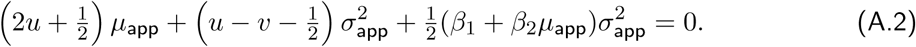

Subtracting Eq. (A.2) from Eq. (A.1):

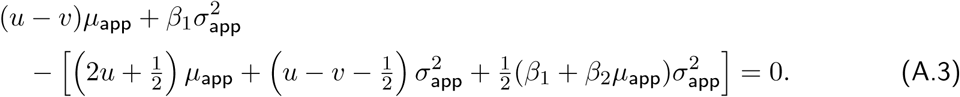

Simplifying and grouping terms:

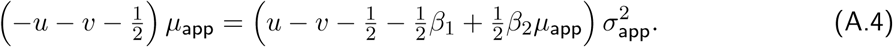

Solving for the dispersion ratio 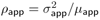:

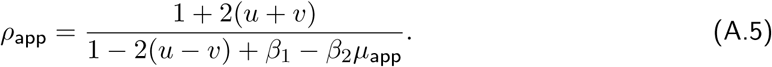

To study conditions for overdispersion, it is simpler to consider the inverse dispersion ratio, which simplifies to

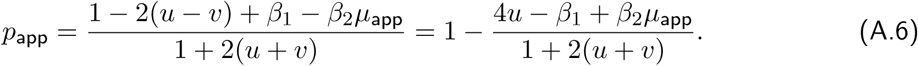

## B Selection function *ϕ* under two fitness functions

The function is given as:

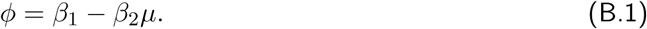

Using the Gaussian fitness function, *w*(*n*) = exp(*−n*^2^*s*),

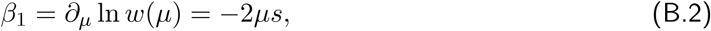

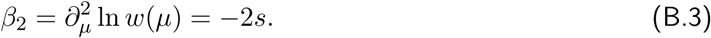

Leading to

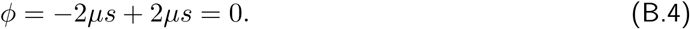

Using the synergistic fitness function, *w*(*n*) = 1 *− n*^2^*s*:

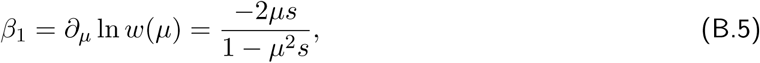

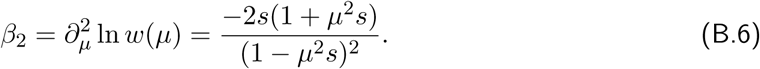

Substituting *β*_1_ and *β*_2_ into 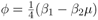:

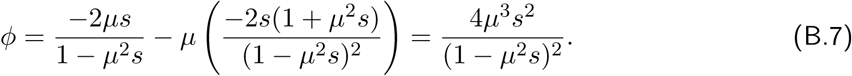

## C Summary statistics of representative TEs

**Table 3:** Summary statistics for three representative TE families (POGO, JOCKEY, MDG1) in the DPGP3 Zambian *D. melanogaster* dataset of Lee (2022) shown in Fig. 6. We report the total number of presence calls summed across sites and strains (*n*_present_), the mean and variance of per-strain TE burdens, the dispersion ratio *ρ*, and higher moments for each family.

| TE Family | Class | $n_{\text{present}}$ | Mean | Variance | Var./Mean ($\rho$) | Skew | Exc. kurtosis |
| --- | --- | --- | --- | --- | --- | --- | --- |
| POGO | DNA | 1704 | 10.39 | 11.45 | 1.1 | 0.6 | 0.25 |
| JOCKEY | non-LTR | 2968 | 18.1 | 21.44 | 1.19 | 0.52 | 0.57 |
| MDG1 | LTR | 1276 | 7.78 | 8.58 | 1.1 | 0.75 | 1.08 |

## D Dispersion ratio versus recombination map length *R*

The main text comparisons in Figs. 3 and 4 focus on free recombination. Figure D.1 extends that comparison across chromosome map lengths *R*, from tightly linked insertions to effectively free recombination, with all quantities measured at the pre-transposition/excision stage, the stage to which the prediction of Roze (2023) applies. We use the same parameter set as the top-right panel of Figure 1 in Roze (2023), i.e., on average 10 TEs per genome. Simulations are run with the Gaussian fitness function *w*(*n*) = exp(*−sn*^2^) from the main text and the ectopic-pair fitness of Roze (2023) *w*(*n*) = exp(*−βn_p_*), in which homologous TE copies at non-allelic sites can mispair during recombination. Thus, *n_p_* is the number of pairs of TEs present at different genomic locations. We observe that under lower recombination regime, *n*_p_ is not well approximated by *n*^2^*/*2.

**Figure D.1:**
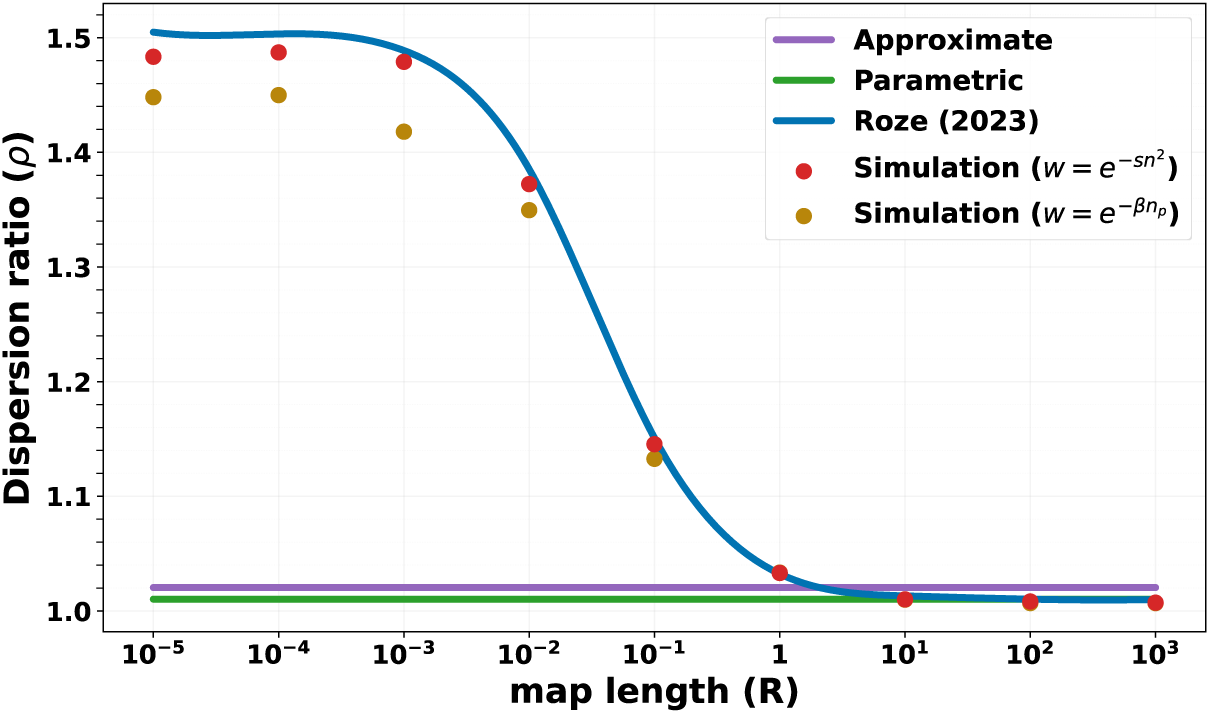
Dispersion ratio versus recombination map length. *R* **(Morgans), pre-transposition/excision.** The figure plots the equilibrium dispersion ratio *ρ* as a function of chromosome map length *R* (log scale), measured immediately after recombination and before transposition and excision. Smooth lines show the approximate moment closure (purple), from Eq. (S159) of the Supplementary Methods, and the parametric negative-binomial closure (green), from Eq. (23); the blue curve shows the dispersion ratio of Roze (2023), calculated from their Eq. (14). Red markers are stochastic simulations with fitness function *w*(*n*) = exp(*−sn*^2^) acting on total diploid copy number *n*, while the gold markers are simulations using the fitness function *w*(*n*_p_) = exp(*−βn*_p_) with ectopic TE pairs *n*_p_ (Roze, 2023), both rec orded at the same stage. Both sets of simulations were generated with parameters: *N* = 10^4^, *u* = 10*^−^*^2^, *v* = *u/*100, *s* = *u/*20, and *β* = *u/*10, which ran for 30 000 generations, and equilibrium summaries from piecewise means every 100 generations after burn-in from 9,000 generations. Numerical means, variances, dispersion ratios, and higher moments for these runs are reported in Supplementary Tables S7–S8.

## E Dispersion ratio under varying fitness curvature

The results in the main text use the Gaussian fitness function *w*(*n*) = exp(*−sn*^2^). Here we vary the exponent *γ* in *w*(*n*) = exp(*−sn^γ^*), which tunes how sharply fitness declines with copy number and hence the curvature *β*_2_ of the log-fitness function. Small *γ* gives a gradual decline, while large *γ* produces a steep, threshold-like drop close to a reference copy number (Fig. E.1, left panel). This lets us test the prediction of Section 2.6 that the pre-transposition/excision copy number *m^′^* can be either over- or underdispersed depending on *β*_2_, whereas the post-transposition/excision copy number remains overdispersed throughout.

**Figure E.1:**
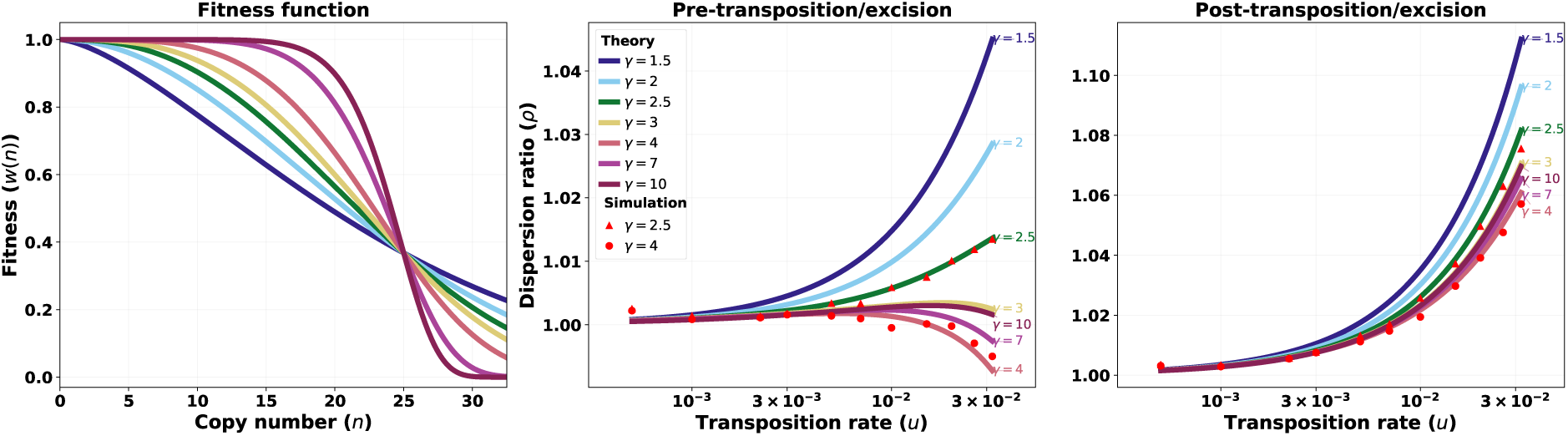
Effect of fitness curvature on the dispersion ratio at the two life-cycle stages. Left panel: the fitness function *w*(*n*) = exp(*−sn^γ^*)*, γ ∈* [1.5, 10]. Middle and right panels: the equilibrium dispersion ratio at the pre-transposition/excision stage (*ρ_m_′*, Eqs. (19)–(20)) and at the post-transposition/excision stage (*ρ*, Eq. (12)), as a function of the transposition rate *u* (log scale). Lines are the parametric negative-binomial closure, evaluated; red markers are stochastic simulations for *γ* = 2.5 (triangles) and *γ* = 4 (circles). At the post-transposition/excision stage all curves stay above *ρ* = 1, whereas at the pre-transposition/excision stage sufficiently strong curvature (e.g., *γ* = 4) drives *ρ_m_′* below one, confirming that underdispersion is possible at this earlier stage (Supplementary Section S5.4). Simulation parameters: *N* = 10^4^, *s* = 1*/N*, *v* = 10*^−^*^4^, free (binomial) recombination, run for 50 000 generations with equilibrium summaries taken from piecewise means every 100 generations after a burn-in of 9 000 generations.

## References

Adeosun, S. A. and Pfaffelhuber, P. Markov processes forced on a subspace by a large drift, with applications to population genetics. *arXiv pre-print*, 2026. doi: 10.48550/arxiv.2602.16342.

Bartolomé, C., Bello, X., and Maside, X. Widespread evidence for horizontal transfer of transposable elements across drosophila genomes. Genome biology, 10(2):R22, 2009. doi: 10.1186/gb-2011-12-11-411.

Betancourt, A. J., Wei, K. H.-C., Huang, Y., and Lee, Y. C. G. Causes and consequences of varying transposable element activity: An evolutionary perspective. Annual review of genomics and human genetics, 25(1):1–25, 2024. doi: 10.1146/annurev-genom-120822-105708.

Bourque, G., Burns, K. H., Gehring, M., Gorbunova, V., Seluanov, A., et al. Ten things you should know about transposable elements. Genome Biology, 19:1–12, 2018. doi: 10.1186/s13059-018-1577-z.

Charlesworth, B. and Langley, C. The evolution of self-regulated transposition of transposable elements. Genetics, 112(2):359, 1986. doi: 10.1093/genetics/112.2.359.

Charlesworth, B. and Charlesworth, D. The population dynamics of transposable elements. Genetics Research, 42(1):1–27, 1983. doi: 10.1017/S0016672300021455.

de la Chaux, N. and Wagner, A. Evolutionary dynamics of the LTR retrotransposons *roo* and *rooA* inferred from twelve complete *Drosophila* genomes. BMC Evolutionary Biology, 9:205, 2009. doi: 10.1186/1471-2148-9-205.

Díaz-González, J. and Domínguez, A. Different structural variants of *roo* retrotransposon are active in *Drosophila melanogaster*. Gene, 741:144546, 2020. doi: 10.1016/j.gene.2020.144546.

Dolgin, E. S. and Charlesworth, B. The effects of recombination rate on the distribution and abundance of transposable elements. Genetics, 178(4):2169–2177, 2008.

Doolittle, W. F. and Sapienza, C. Selfish genes, the phenotype paradigm and genome evolution. Nature, 284(5757):601–603, 1980. doi: 10.1038/284601a0.

Etheridge, A. M. *Some Mathematical Models from Population Genetics: École d’Été de Probabilités de Saint-Flour XXXIX-*2009, volume 2012 of *Lecture Notes in Mathematics*. Springer-Verlag, Berlin, Heidelberg, 2011. ISBN 978-3-642-16631-0. doi: 10.1007/978-3-642-16632-7.

Ewing, A. D. and Kazazian, H. H. High-throughput sequencing reveals extensive variation in human-specific L1 content in individual human genomes. Genome research, 20(9):1262–1270, 2010. doi: 10.1101/gr.106419.110.

García Guerreiro, M. P. What makes transposable elements move in the *Drosophila* genome? Heredity, 108(5):461–468, 2012. doi: 10.1038/hdy.2011.89.

Huang, Y. and Lee, Y. C. G. Blessing or curse: how the epigenetic resolution of host-transposable element conflicts shapes their evolutionary dynamics. Proceedings of the Royal Society B, 291 (2020):20232775, 2024. doi: 10.1098/rspb.2023.2775.

Kapitonov, V. V. and Jurka, J. Molecular paleontology of transposable elements in the drosophila melanogaster genome. Proceedings of the National Academy of Sciences, 100(11):6569–6574, 2003. doi: 10.1073/pnas.0732024100.

Kazazian, H. H. Mobile elements: Drivers of genome evolution. Science, 303(5664):1626–1632, 2004. doi: 10.1126/science.1089670.

Kelleher, E. S., Barbash, D. A., and Blumenstiel, J. P. Taming the turmoil within: New insights on the containment of transposable elements. Trends in Genetics, 36(7):474—-489, 2020. doi: 10.1016/j.tig.2020.04.007.

Kofler, R. Dynamics of transposable element invasions with piRNA clusters. Molecular Biology and Evolution, 36(7):1457–1472, 2019. doi: 10.1093/molbev/msz079.

Kofler, R., Hill, T., Nolte, V., Betancourt, A. J., and Schlötterer, C. The recent invasion of natural *Drosophila simulans* populations by the P-element. Proceedings of the National Academy of Sciences, 112(21):6659–6663, 2015. doi: 10.1073/pnas.1500758112.

Langmüller, A. M., Nolte, V., Dolezal, M., and Schlötterer, C. The genomic distribution of transposable elements is driven by spatially variable purifying selection. Nucleic Acids Research, 51(17): 9203–9213, 2023. doi: 10.1093/nar/gkad635.

Langmüller, A. M., Haller, B. C., Nolte, V., and Schlötterer, C. Purifying selection shapes the dynamics of P-element invasion in *Drosophila simulans* populations. Genome Biology, 26(1):221, 2025. doi: 10.5281/zenodo.15710171.

Le Rouzic, A. and Capy, P. Population genetics models of competition between transposable element subfamilies. Genetics, 174(2):785–793, 2006a. doi: 10.1534/genetics.105.052241.

Le Rouzic, A. and Capy, P. Population genetics models of competition between transposable element subfamilies. Genetics, 174(2):785–793, 2006b. doi: 10.1534/genetics.105.052241.

Le Rouzic, A., Boutin, T. S., and Capy, P. Long-term evolution of transposable elements. Proceedings of the National Academy of Sciences, 104(49):19375–19380, 2007. doi: 10.1073/pnas.0705238104.

Lee, Y. C. G. Synergistic epistasis of the deleterious effects of transposable elements. Genetics, 220 (2):iyab211, 2022. doi: 10.1093/genetics/iyab211.

Maside, X., Bartolomé, C., and Charlesworth, B. Inferences on the evolutionary history of the S-element family of *Drosophila melanogaster*. Molecular Biology and Evolution, 20(8):1183–1187, 2003. doi: 10.1093/molbev/msg120.

McGurk, M. P., Dion-Côté, A.-M., and Barbash, D. A. Rapid evolution at the *Drosophila* telomere: transposable element dynamics at an intrinsically unstable locus. Genetics, 217(2):iyaa027, 2021. doi: 10.1093/genetics/iyaa027.

Oggenfuss, U. and Croll, D. Recent transposable element bursts are associated with the proximity to genes in a fungal plant pathogen. PLoS Pathogens, 19(2):e1011130, 2023. doi: 10.1371/journal.ppat.1011130.

Omole, A. D. and Czuppon, P. Maintenance of long-term transposable element activity through regulation by nonautonomous elements. Genetics, 229(2):iyae209, 2025. doi: 10.1093/genetics/iyae209.

Orgel, L. E. and Crick, F. H. C. Selfish DNA: the ultimate parasite. Nature, 284(5757):604–607, 1980. doi: 10.1038/284604a0.

Pfaffelhuber, P. and Wakolbinger, A. A diploid population model for copy number variation of genetic elements. Electronic Journal of Probability, 28:1–15, 2023. doi: 10.1214/23-EJP934.

Roze, D. Causes and consequences of linkage disequilibrium among transposable elements within eukaryotic genomes. Genetics, 224(2):iyad058, 2023. doi: 10.1093/genetics/iyad058.

Said, I., McGurk, M. P., Clark, A. G., and Barbash, D. A. Patterns of piRNA regulation in *Drosophila* revealed through transposable element clade inference. Molecular Biology and Evolution, 39(1): msab336, 2022. doi: 10.1093/molbev/msab336.

Scarpa, A. and Kofler, R. The impact of paramutations on the invasion dynamics of transposable elements. Genetics, 225(4):iyad181, 2023. doi: 10.1093/genetics/iyad181.

Schrader, L. and Schmitz, J. The impact of transposable elements in adaptive evolution. Molecular Ecology, 28(6):1537–1549, 2019. doi: 10.1111/mec.14794.

Shapiro, J. Mobile genetic elements. Elsevier, 2012.

Smith, R. D., Puzey, J. R., and Conradi Smith, G. D. Population genetics of transposable element load: A mechanistic account of observed overdispersion. Plos one, 17(7):e0270839, 2022. doi: 10.1371/journal.pone.0270839.

Tomar, S. S., Hua-Van, A., and Le Rouzic, A. A population genetics theory for piRNA-regulated transposable elements. Theoretical Population Biology, 150:1–13, 2023. doi: 10.1016/j.tpb.2023.02.001.

Udomkit, A., Forbes, S., Dalgleish, G., and Finnegan, D. BS a novel LINE-like element in *Drosophila melanogaster*. Nucleic Acids Research, 23(8):1354–1358, 1995. doi: 10.1093/nar/23.8.1354.

Wierzbicki, F. and Kofler, R. The composition of piRNA clusters in *Drosophila melanogaster* deviates from expectations under the trap model. BMC biology, 21(1):224, 2023. doi: 10.1186/s12915-023-01727-7.

Wierzbicki, F., Pianezza, R., Selvaraju, D., Eller, M. M., and Kofler, R. On the origin of the P-element invasion in *Drosophila simulans*. Mobile DNA, 16(1):1–10, 2025. doi: 10.1186/s13100-025-00345-0.

