## Supplementary_material for "Population genetics of active transposable elements: overdispersion arises naturally from transposition-deletion-selection balance"

### Supplementary Material: A population genetics model explaining overdispersion in active transposable elements

#### S1 Derivation of TE mean and variance dynamics

Following from the model section of the main text, at each reproduction event of the bi-parent Moran model, two individuals are chosen with probabilities proportional to their fitness values $w(l)$ and $w(n)$ (fertility selection), where $l$ and $n$ denote the TE copy numbers in their diploid genomes. Each TE copy transmitted from the parent to the offspring is inherited independently with probability $1/2$ (free recombination). If the parents together carry $l+n$ elements, the offspring's TE count before transposition/deletion, $m'$, follows a binomial distribution with success probability 1/2, reflecting independent segregation:

$$\mathbb{P}_{\text{recomb}}(m' \mid l, n) = \binom{l+n}{m'} 2^{-(l+n)} \ . \tag{S1}$$

After recombination, the inherited number of TE copies, denoted $m'$, may change. We denote by $u$ the transposition rate per TE and by $v$ the excision rate. We assume that these rates are low enough that the probability of more than one event occurring in a single time step, i.e., a generation, is negligible. In this case, the exponential terms of the transition probabilities of the associated Poisson processes may be linearized as follows. With a probability of $1-(u+v)m'$, the number of TEs remains unchanged at $m = m'$. A transposition event, which increases the TE count by one ($m = m'+1$), occurs with probability $um'$. Conversely, a deletion event, which decreases the TE count by one ($m = m'-1$), occurs with probability $vm'$. Once the final TE count $m$ is determined, the offspring replaces a random individual in the population.

The state of the TE population of the bi-parent Moran model is described by the vector $x = (x_0, x_1, \ldots)$, where $x_m$ is the fraction of individuals with $m$ elements. For any test function $f(x)$, the infinitesimal generator $\mathcal{G}f(x)$ gives the expected instantaneous rate of change of $f$:

$$\mathcal{G}f(x) = \frac{N^2}{2} \sum_k x_k \sum_m \left[ f\left(x + \frac{e_m}{N} - \frac{e_k}{N}\right) - f(x) \right] \times \Big[ R_0(m) + R_+(m) + R_-(m) \Big], \tag{S2}$$

where $e_m$ and $e_k$ denote the unit vector that add one individual with $m$ elements and remove one individual with $k$ elements, respectively. The terms $R_0(m)$, $R_+(m)$, and $R_-(m)$ are the probabilities of producing an offspring with $m$ elements via no change, transposition, or deletion, respectively:

$$R_0(m) = \sum_{l,n} \frac{x_l\, w(l)}{\bar{w}(x)} \cdot \frac{x_n\, w(n)}{\bar{w}(x)} \cdot \binom{l+n}{m} 2^{-(l+n)} \cdot [1-(u+v)m]\,,$$
$$R_+(m) = \sum_{l,n} \frac{x_l\, w(l)}{\bar{w}(x)} \cdot \frac{x_n\, w(n)}{\bar{w}(x)} \cdot \binom{l+n}{m-1} 2^{-(l+n)} \cdot u(m-1), \tag{S3}$$
$$R_-(m) = \sum_{l,n} \frac{x_l\, w(l)}{\bar{w}(x)} \cdot \frac{x_n\, w(n)}{\bar{w}(x)} \cdot \binom{l+n}{m+1} 2^{-(l+n)} \cdot v(m+1),$$

where $\bar{w}(x) = \sum_j x_j\, w(j)$ denotes the mean fitness of the population. This is the basis for the derivation of the TE mean and variance dynamics in the following sections.

#### S1.1 Derivation of mean copy number dynamics

To derive the dynamics of the mean TE copy number, we apply the infinitesimal generator framework with the test function $f(x) = g(\mu(x))$, where $\mu(x) = \sum_k kx_k$ represents the mean copy number of TEs per individual in the population.

The population state change from a single reproduction-replacement event is:

$$\mu\left(x + \frac{e_m}{N} - \frac{e_k}{N}\right) - \mu(x) = \frac{m-k}{N}. \tag{S4}$$

Using a Taylor expansions around the current population mean yields:

$$g\left(\mu\left(x + \frac{e_m}{N} - \frac{e_k}{N}\right)\right) - g(\mu(x)) = \frac{1}{N} g'(\mu(x))(m-k) + \frac{1}{2N^2} g''(\mu(x))(m-k)^2 + \mathcal{O}(N^{-3}). \tag{S5}$$

Substituting this expansion into the infinitesimal generator from Eq. (S2), we can rewrite it by

$$\mathcal{G}g(\mu(x)) = G_1(g(\mu(x))) + G_2(g(\mu(x))) + \mathcal{O}(N^{-1}), \tag{S6}$$

where $G_1$ captures the first-order deterministic dynamics evolving on a $\mathcal{O}(N)$ timescale, while $G_2$ represents second-order stochastic fluctuations that, though scaled by $1/N^2$, still contribute on the same $N$-generation timescale, at least in finite populations. In what follows we retain only $G_1$, i.e. the leading deterministic drift in $\mu$.

The first-order term is

$$G_1(g(\mu(x))) = \frac{N}{2} g'(\mu(x)) \sum_k x_k \sum_m (m-k) \left[R_0(m) + R_+(m) + R_-(m)\right]. \tag{S7}$$

We now compute each reproduction term separately. For $R_0(m)$, representing offspring with an unchanged TE count after transposition-deletion events we have

$$\begin{aligned}\sum_m (m-k)R_0(m) &= \sum_m (m-k)\sum_{l,n}\frac{x_l x_n w(l)w(n)}{\bar{w}^2(x)}\binom{l+n}{m}2^{-(l+n)}(1-(u+v)m)\\ &= \frac{1}{\bar{w}^2(x)}\sum_{l,n}x_l x_n w(l)w(n)2^{-(l+n)}\sum_m (m-k)\binom{l+n}{m}(1-(u+v)m)\\ &= \frac{1}{\bar{w}^2(x)}\sum_{l,n}x_l x_n w(l)w(n)2^{-(l+n)}\\ &\quad\times\left[\sum_m m\binom{l+n}{m} - (u+v)\sum_m m^2\binom{l+n}{m} - k\sum_m\binom{l+n}{m} + k(u+v)\sum_m m\binom{l+n}{m}\right].\end{aligned} \tag{S8}$$

For the transposition term $R_+(m)$, we need to account for the index shift (offspring with $m$ TEs arose from $m-1$ TEs after transposition):

$$\begin{aligned}\sum_m (m-k)R_+(m) &= \sum_m (m-k)\sum_{l,n}\frac{x_l x_n w(l)w(n)}{\bar{w}^2(x)}\binom{l+n}{m-1}2^{-(l+n)}u(m-1)\\ &= \frac{u}{\bar{w}^2(x)}\sum_{l,n}x_l x_n w(l)w(n)2^{-(l+n)}\sum_m (m-k)(m-1)\binom{l+n}{m-1}.\end{aligned} \tag{S9}$$

Let $j=m-1$, so $m=j+1$ and the sum becomes:

$$\begin{aligned}\sum_m (m-k)(m-1)\binom{l+n}{m-1} &= \sum_j (j+1-k)j\binom{l+n}{j}\\ &= \sum_j j(j+1)\binom{l+n}{j} - k\sum_j j\binom{l+n}{j}\\ &= \sum_j j^2\binom{l+n}{j} + \sum_j j\binom{l+n}{j} - k\sum_j j\binom{l+n}{j}\\ &= \sum_j j^2\binom{l+n}{j} + (1-k)\sum_j j\binom{l+n}{j}.\end{aligned} \tag{S10}$$

Therefore:

$$\sum_m (m-k)R_+(m) = \frac{u}{\bar{w}^2(x)}\sum_{l,n}x_l x_n w(l)w(n)2^{-(l+n)}\left[\sum_j j^2\binom{l+n}{j} + (1-k)\sum_j j\binom{l+n}{j}\right]. \tag{S11}$$

Similarly, for the deletion term $R_-(m)$, offspring with $m$ TEs arose from $m+1$ TEs after deletion:

$$\begin{aligned}\sum_m (m-k)R_-(m) &= \sum_m (m-k)\sum_{l,n}\frac{x_l x_n w(l)w(n)}{\bar{w}^2(x)}\binom{l+n}{m+1}2^{-(l+n)}v(m+1)\\ &= \frac{v}{\bar{w}^2(x)}\sum_{l,n}x_l x_n w(l)w(n)2^{-(l+n)}\sum_m (m-k)(m+1)\binom{l+n}{m+1}.\end{aligned} \tag{S12}$$

Let $j = m + 1$, so $m = j - 1$ and the sum becomes:

$$\begin{aligned}\sum_m (m-k)(m+1)\binom{l+n}{m+1} &= \sum_j (j-1-k)j\binom{l+n}{j} \\ &= \sum_j j(j-1)\binom{l+n}{j} - k\sum_j j\binom{l+n}{j} \\ &= \sum_j j^2\binom{l+n}{j} - \sum_j j\binom{l+n}{j} - k\sum_j j\binom{l+n}{j} \\ &= \sum_j j^2\binom{l+n}{j} - (1+k)\sum_j j\binom{l+n}{j}.\end{aligned} \tag{S13}$$

Therefore:

$$\sum_m (m-k)R_-(m) = \frac{v}{\bar{w}^2(x)}\sum_{l,n} x_l x_n w(l)w(n)2^{-(l+n)}\left[\sum_j j^2\binom{l+n}{j} - (1+k)\sum_j j\binom{l+n}{j}\right]. \tag{S14}$$

Combining all three terms:

$$\begin{aligned}&\sum_m (m-k)[R_0 + R_+ + R_-] \\ &= \frac{1}{\bar{w}^2(x)}\sum_{l,n} x_l x_n w(l)w(n)2^{-(l+n)} \\ &\times \left[\sum_m m\binom{l+n}{m} - (u+v)\sum_m m^2\binom{l+n}{m} - k\sum_m \binom{l+n}{m} + k(u+v)\sum_m m\binom{l+n}{m}\right. \\ &\quad + u\sum_m m^2\binom{l+n}{m} + u(1-k)\sum_m m\binom{l+n}{m} \\ &\quad \left. + v\sum_m m^2\binom{l+n}{m} - v(1+k)\sum_m m\binom{l+n}{m}\right].\end{aligned} \tag{S15}$$

Collecting like terms:

- Terms with $\sum_m m\binom{l+n}{m}$: $1 + k(u+v) + u(1-k) - v(1+k) = 1 + u - v$
- Terms with $\sum_m m^2\binom{l+n}{m}$: $-(u+v) + u + v = 0$
- Terms with $\sum_m \binom{l+n}{m}$: $-k$

This simplifies to:

$$\begin{aligned}&\sum_m (m-k)[R_0 + R_+ + R_-] \\ &= \frac{1}{\bar{w}^2(x)}\sum_{l,n} x_l x_n w(l)w(n)2^{-(l+n)}\left[(1+u-v)\sum_m m\binom{l+n}{m} - k\sum_m \binom{l+n}{m}\right].\end{aligned} \tag{S16}$$

Using the binomial moment identities $\sum_m \binom{l+n}{m} 2^{-(l+n)} = 1$ and $\sum_m m\binom{l+n}{m} 2^{-(l+n)} = \frac{l+n}{2}$:

$$\sum_m (m-k)[R_0 + R_+ + R_-] = \frac{1}{\bar{w}^2(x)} \sum_{l,n} x_l x_n w(l) w(n) \left[(1+u-v)\frac{l+n}{2} - k\right]. \tag{S17}$$

To proceed, we employ a second-order Taylor expansion of the fitness function around the population mean $\mu$:

$$w(l) \approx w(\mu) + w'(\mu)(l-\mu) + \frac{w''(\mu)}{2}(l-\mu)^2. \tag{S18}$$

For products of fitness functions, neglecting terms of order $(w')^2$, $w'w''$, and $(w'')^2$ (which for typically used fitness functions are of order $N^{-2}$ in the weak selection regime and under classical selection scaling, i.e., $s \in O(N^{-1})$):

$$\begin{aligned} w(l)\cdot w(n) &\approx \left[w(\mu) + w'(\mu)(l-\mu) + \frac{w''(\mu)}{2}(l-\mu)^2\right]\left[w(\mu) + w'(\mu)(n-\mu) + \frac{w''(\mu)}{2}(n-\mu)^2\right] \\ &\approx w(\mu)^2 + w(\mu)w'(\mu)[(l-\mu)+(n-\mu)] + w(\mu)\frac{w''(\mu)}{2}[(l-\mu)^2 + (n-\mu)^2]. \end{aligned} \tag{S19}$$

Using Eqs. (S16) and (S17), and substituting the fitness approximation into $G_1$ (Eq. (S7)), we find:

$$\begin{aligned} G_1(g(\mu(x))) &= \frac{N}{2} g'(\mu(x)) \frac{1}{\bar{w}^2(x)} \sum_{k,l,n} x_k x_l x_n w(l) w(n) \left[(1+u-v)\frac{l+n}{2} - k\right] \\ &= \frac{N}{2} g'(\mu(x)) \frac{1}{\bar{w}^2(x)} \sum_{k,l,n} x_k x_l x_n \\ &\quad \times \left[w(\mu)^2 + w(\mu)w'(\mu)[(l-\mu)+(n-\mu)] + w(\mu)\frac{w''(\mu)}{2}[(l-\mu)^2 + (n-\mu)^2]\right] \\ &\quad \times \left[(1+u-v)\frac{l+n}{2} - k\right]. \end{aligned} \tag{S20}$$

We now evaluate each term systematically:

*Terms with $w(\mu)^2$:*

$$\begin{aligned} &w(\mu)^2 \sum_{k,l,n} x_k x_l x_n \left[(1+u-v)\frac{l+n}{2} - k\right] \\ &= w(\mu)^2 \left[(1+u-v)\frac{1}{2}\sum_{k,l,n} x_k x_l x_n (l+n) - \sum_{k,l,n} x_k x_l x_n k\right] \\ &= w(\mu)^2 \left[(1+u-v)\frac{1}{2}\cdot 2\mu - \mu\right] \quad \text{(since } \sum_k x_k = \sum_l x_l = \sum_n x_n = 1\text{)} \\ &= w(\mu)^2[(1+u-v)\mu - \mu] \\ &= w(\mu)^2(u-v)\mu. \end{aligned} \tag{S21}$$

*Terms with $w'(\mu)$:*

$$\begin{aligned}
&w(\mu)w'(\mu)\sum_{k,l,n} x_k x_l x_n[(l-\mu)+(n-\mu)]\left[(1+u-v)\frac{l+n}{2}-k\right] \\
&\approx w(\mu)w'(\mu)\sum_{k,l,n} x_k x_l x_n[(l-\mu)+(n-\mu)]\left[\frac{l+n}{2}-k\right] \\
&= 2w(\mu)w'(\mu)\sum_{k,l,n} x_k x_l x_n(l-\mu)\left[\frac{l+n}{2}-k\right] \quad \text{(by symmetry in } l \text{ and } n).
\end{aligned} \tag{S22}$$

The approximation is justified by assuming that transposition and deletion rates scale with the inverse population size, i.e., the product of $w'(\mu)(u-v)$ is of order $N^{-2}$ (or smaller).

Expanding the product:

$$\begin{aligned}
&2w(\mu)w'(\mu)\sum_{k,l,n} x_k x_l x_n(l-\mu)\left[\frac{l+n}{2}-k\right] \\
&= w(\mu)w'(\mu)\sum_{k,l,n} x_k x_l x_n\left[l^2+ln-\mu l-\mu n-2kl+2k\mu\right].
\end{aligned} \tag{S23}$$

Evaluating each sum using $\sum_k x_k = 1$, $\sum_k x_k k = \mu$, and $\sum_k x_k k^2 = \mu^2+\sigma^2$:

$$\begin{aligned}
&= w(\mu)w'(\mu)\left[(\mu^2+\sigma^2)+\mu^2-\mu^2-\mu^2-2\mu^2+2\mu^2\right] \\
&= w(\mu)w'(\mu)\sigma^2 \\
&= w(\mu)^2\beta_1\sigma^2.
\end{aligned} \tag{S24}$$

In the last equality, we have rewritten the fitness derivative using logarithmic derivatives:

$$\begin{aligned}
\beta_1 &= \frac{w'(\mu)}{w(\mu)} = \frac{\partial}{\partial\mu}\ln w(\mu), \\
\beta_2 &= \frac{w''(\mu)}{w(\mu)} - \left(\frac{w'(\mu)}{w(\mu)}\right)^2 = \frac{\partial^2}{\partial\mu^2}\ln w(\mu).
\end{aligned} \tag{S25}$$

This gives us $w'(\mu) = w(\mu)\beta_1$ and $w''(\mu) = w(\mu)(\beta_1^2+\beta_2)$, thus explaining the last equality.

*Terms with $w''(\mu)$:* Following similar steps and using the assumption that $u, v \in \mathcal{O}(1/N)$, $\beta_1, \beta_2 \in$

$\mathcal{O}(s)$, $s \in \mathcal{O}(1/N)$ and dropping terms of $\mathcal{O}(1/N^2)$:

$$\begin{aligned}
&\frac{w(\mu)w''(\mu)}{2}\sum_{k,l,n} x_k x_l x_n \left[(l-\mu)^2 + (n-\mu)^2\right]\left((1+u-v)\left(\frac{l+n}{2}\right)-k\right)\\
&\approx w(\mu)w''(\mu)\sum_{k,l,n} x_k x_l x_n (l-\mu)^2\left(\frac{l+n}{2}-k\right)\\
&= w(\mu)w''(\mu)\sum_{k,l,n} x_k x_l x_n \left(\left(\frac{l+n}{2}\right)(l-\mu)^2 - k(l-\mu)^2\right)\\
&= w(\mu)w''(\mu)\left(\frac{1}{2}(\sigma^3\gamma+\mu\sigma^2)+\frac{1}{2}\mu\sigma^2-\mu\sigma^2\right)\\
&= w(\mu)w''(\mu)\left(\frac{1}{2}\sigma^3\gamma\right)\\
&= \frac{w(\mu)^2\beta_2}{2}\sigma^3\gamma.
\end{aligned} \tag{S26}$$

where $\gamma = \mathbb{E}[(X-\mu)^3]/\sigma^3$ is the population skewness and the third moment is rewritten as $\mathbb{E}[X^3] = \sigma^3\gamma + 3\mu\sigma^2 + \mu^3$.

The population mean fitness is:

$$\begin{aligned}
\bar{w}(x) &= \sum_j x_j w(j) \approx \sum_j x_j \left[w(\mu) + w'(\mu)(j-\mu) + \frac{w''(\mu)}{2}(j-\mu)^2\right]\\
&= w(\mu) + 0 + \frac{w''(\mu)}{2}\sigma^2 = w(\mu)\left(1+\frac{\beta_2}{2}\sigma^2\right).
\end{aligned} \tag{S27}$$

Therefore: $\bar{w}^2(x) \approx w^2(\mu)\left(1+\beta_2\sigma^2\right)$.

Combining all contributions into Eq. (S20), we have:

$$\begin{aligned}
G_1(g(\mu)) &= \frac{N}{2}g'(\mu)\frac{w^2(\mu)}{w^2(\mu)(1+\beta_2\sigma^2)}\left[(u-v)\mu + \beta_1\sigma^2 + \frac{\beta_2}{2}\sigma^3\gamma\right]\\
&\approx g'(\mu)\frac{N}{2}\left[(u-v)\mu + \beta_1\sigma^2 + \frac{\beta_2}{2}\sigma^3\gamma\right],
\end{aligned} \tag{S28}$$

where the approximation is again justified by dropping terms of order $N^{-2}$.

Converting from the discrete Moran timescale (events per $N/2$ time units) to continuous time steps by rescaling time, we find the following deterministic dynamics of the mean TE copy number per genome:

$$\frac{d\mu}{dt} = (u-v)\mu + \beta_1\sigma^2 + \frac{\beta_2}{2}\sigma^3\gamma, \tag{S29}$$

or equivalently

$$\frac{d\mu(x)}{dt} = (u-v)\mu(x) + \sigma^2(x)\frac{\partial}{\partial\mu}\ln w(\mu(x)) + \frac{1}{2}\gamma(x)\sigma^3(x)\frac{\partial^2}{\partial\mu^2}\ln w(\mu(x)). \tag{S30}$$

#### S1.2 Derivation of variance dynamics

We derive the dynamics of the variance $\sigma^2(x) = \psi(x) - \mu(x)^2$, where $\psi(x) = \sum_m m^2 x_m$ is the second moment and $\mu(x) = \sum_m m x_m$ is the mean. The generator $\mathcal{G}$ acts on $\sigma^2(x)$ as:

$$\mathcal{G}g(\sigma^2(x)) = \frac{N^2}{2}\sum_k x_k \sum_m \left[ g\Big(\sigma^2\left(x + \frac{e_m}{N} - \frac{e_k}{N}\right)\Big) - g(\sigma^2(x))\right] \cdot [R_0(m) + R_+(m) + R_-(m)] \,. \tag{S31}$$

First, we compute the change in $\sigma^2(x)$ under the perturbation $x \mapsto x + \frac{e_m}{N} - \frac{e_k}{N}$:

$$\begin{aligned}\sigma^2\left(x + \frac{e_m}{N} - \frac{e_k}{N}\right) - \sigma^2(x) &= \left(\psi\left(x + \frac{e_m}{N} - \frac{e_k}{N}\right) - \mu\left(x + \frac{e_m}{N} - \frac{e_k}{N}\right)^2\right) - \left(\psi(x) - \mu(x)^2\right)\\ &= \left(\psi(x) + \frac{m^2-k^2}{N} - \left(\mu(x) + \frac{m-k}{N}\right)^2\right) - \left(\psi(x) - \mu(x)^2\right)\\ &= \frac{m^2-k^2}{N} - \frac{2\mu(x)(m-k)}{N} - \frac{(m-k)^2}{N^2}.\end{aligned} \tag{S32}$$

The generator applied to $\sigma^2(x)$ becomes (only keeping the first derivative of $g$):

$$\begin{aligned}G_1(g(\sigma^2(x))) = \frac{N^2}{2} g'(\sigma^2(x))) \sum_k x_k \sum_m &\left[\frac{m^2-k^2}{N} - \frac{2\mu(x)(m-k)}{N} - \frac{(m-k)^2}{N^2}\right]\\ &\times [R_0(m) + R_+(m) + R_-(m)]\\ = \frac{N}{2} g'(\sigma^2(x)))\Big(\sum_k x_k \sum_m &\left[(m^2-k^2) - 2\mu(x)(m-k)\right][R_0 + R_+ + R_-]\\ - \frac{1}{N}\sum_k x_k &\sum_m (m-k)^2 [R_0 + R_+ + R_-]\Big)\end{aligned} \tag{S33}$$

We now expand the terms one after the other. For the first term, using the same index shifts as applied in Eqs. (S10) and (S13), we find:

$$\begin{aligned}\sum_m (m^2-k^2)[R_0 + R_+ + R_-] = \frac{1}{\bar{w}(x)^2}\sum_{l,n} x_l x_n w(l) w(n) 2^{-(l+n)} &\Bigg[\sum_m m^2 \binom{l+n}{m}(1-(u+v)m)\\ - k^2 \sum_m \binom{l+n}{m}(1-(u+v)m) + u\sum_m (m^3 + 2m^2 + m)&\binom{l+n}{m}\\ - uk^2 \sum_m m\binom{l+n}{m} + v\sum_m (m^3 - 2m^2 + m)\binom{l+n}{m} - vk^2 &\sum_m m\binom{l+n}{m}\Bigg]\end{aligned} \tag{S34}$$

After cancellation of $m^3$ terms and collection:

$$= \frac{1}{\bar{w}(x)^2}\sum_{l,n} x_l x_n w(l) w(n)\left[(1+2u-2v)\sum_m m^2\binom{l+n}{m}2^{-(l+n)} + (u+v)\sum_m m\binom{l+n}{m}2^{-(l+n)} - k^2\right] \tag{S35}$$

For the second term in Eq. (S33), we have from Eq. (S17)

$$\sum_m (m-k)[R_0 + R_+ + R_-] = \frac{1}{\bar{w}^2(x)} \sum_{l,n} x_l x_n w(l) w(n) \Big[(1+u-v)\left(\frac{l+n}{2}\right) - k\Big] \tag{S36}$$

For the third term in Eq. (S33) we have:

$$\begin{aligned}\sum_m (m-k)^2[R_0 + R_+ + R_-] = &\sum_m (m^2 - 2km + k^2) R_0(m) + \sum_m (m^2 - 2km + k^2) R_+(m) \\ &+ \sum_m (m^2 - 2km + k^2) R_-(m)\end{aligned} \tag{S37}$$

For the term $R_0(m)$, we substitute its exact form (Eq. (S3)) and expand:

$$\begin{aligned}\sum_m (m-k)^2 R_0(m) &= \frac{1}{\bar{w}(x)^2} \sum_{l,n} x_l x_n w(l) w(n) 2^{-(l+n)} \sum_m (m^2 - 2km + k^2) \binom{l+n}{m} (1-(u+v)m) \\ &= \frac{1}{\bar{w}(x)^2} \sum_{l,n} x_l x_n w(l) w(n) 2^{-(l+n)} \Bigg[\sum_m m^2 \binom{l+n}{m} - (u+v) \sum_m m^3 \binom{l+n}{m} \\ &\quad - 2k \sum_m m \binom{l+n}{m} + 2k(u+v) \sum_m m^2 \binom{l+n}{m} \\ &\quad + k^2 \sum_m \binom{l+n}{m} - k^2(u+v) \sum_m m \binom{l+n}{m}\Bigg]\end{aligned} \tag{S38}$$

For the TE transposition term $R_+(m)$, we begin with the unshifted expression:

$$\sum_m (m^2 - 2km + k^2) R_+(m) = \frac{u}{\bar{w}(x)^2} \sum_{l,n} x_l x_n w(l) w(n) 2^{-(l+n)} \sum_m (m^2 - 2km + k^2)(m-1) \binom{l+n}{m-1} \tag{S39}$$

Before shifting indices, we expand the polynomial factor:

$$= \frac{u}{\bar{w}(x)^2} \sum_{l,n} x_l x_n w(l) w(n) 2^{-(l+n)} \sum_m \Big[m^2(m-1) - 2km(m-1) + k^2(m-1)\Big] \binom{l+n}{m-1} \tag{S40}$$

Now perform the index shift $m \to m+1$:

$$= \frac{u}{\bar{w}(x)^2} \sum_{l,n} x_l x_n w(l) w(n) 2^{-(l+n)} \sum_m \Big[(m+1)^2 m - 2k(m+1)m + k^2 m\Big] \binom{l+n}{m} \tag{S41}$$

Expanding the polynomials after shifting:

$$\begin{aligned}
&= \frac{u}{\bar{w}(x)^2}\sum_{l,n} x_l x_n w(l) w(n) 2^{-(l+n)} \sum_m \Big[(m^3+2m^2+m)-(2km^2+2km)+k^2 m\Big]\binom{l+n}{m} \\
&= \frac{u}{\bar{w}(x)^2}\sum_{l,n} x_l x_n w(l) w(n) 2^{-(l+n)} \Bigg[\sum_m m^3\binom{l+n}{m} + (2-2k)\sum_m m^2\binom{l+n}{m} \\
&\qquad + (1-2k+k^2)\sum_m m\binom{l+n}{m}\Bigg]
\end{aligned} \tag{S42}$$

For the TE deletion term $R_-(m)$, we similarly start with the unshifted form:

$$\sum_m (m^2-2km+k^2)R_-(m) = \frac{v}{\bar{w}(x)^2}\sum_{l,n} x_l x_n w(l) w(n) 2^{-(l+n)} \sum_m (m^2-2km+k^2)(m+1)\binom{l+n}{m+1} \tag{S43}$$

Expanding before shifting:

$$= \frac{v}{\bar{w}(x)^2}\sum_{l,n} x_l x_n w(l) w(n) 2^{-(l+n)} \sum_m \Big[m^2(m+1) - 2km(m+1) + k^2(m+1)\Big]\binom{l+n}{m+1} \tag{S44}$$

Performing the index shift $m \to m-1$:

$$= \frac{v}{\bar{w}(x)^2}\sum_{l,n} x_l x_n w(l) w(n) 2^{-(l+n)} \sum_m \Big[(m-1)^2 m - 2k(m-1)m + k^2 m\Big]\binom{l+n}{m} \tag{S45}$$

Expanding after shifting:

$$\begin{aligned}
&= \frac{v}{\bar{w}(x)^2}\sum_{l,n} x_l x_n w(l) w(n) 2^{-(l+n)} \sum_m \Big[(m^3-2m^2+m)-(2km^2-2km)+k^2 m\Big]\binom{l+n}{m} \\
&= \frac{v}{\bar{w}(x)^2}\sum_{l,n} x_l x_n w(l) w(n) 2^{-(l+n)} \Bigg[\sum_m m^3\binom{l+n}{m} + (-2-2k)\sum_m m^2\binom{l+n}{m} \\
&\qquad + (1+2k+k^2)\sum_m m\binom{l+n}{m}\Bigg]
\end{aligned} \tag{S46}$$

Now combining Eqs. (S38), (S42) and (S46) and inserting into Eq. (S37), and simplifying:

$$
\begin{aligned}
&\sum_m (m-k)^2[R_0+R_+ +R_-] = \\
&\qquad \frac{1}{\bar{w}(x)^2}\sum_{l,n} x_l x_n w(l)w(n)2^{-(l+n)}\Bigg[ \\
&\qquad\quad \sum_m m^2\binom{l+n}{m} - (u+v)\sum_m m^3\binom{l+n}{m} \\
&\qquad\quad - 2k\sum_m m\binom{l+n}{m} + 2k(u+v)\sum_m m^2\binom{l+n}{m} \\
&\qquad\quad + k^2\sum_m \binom{l+n}{m} - k^2(u+v)\sum_m m\binom{l+n}{m} \\
&\qquad\quad + u\sum_m m^3\binom{l+n}{m} + u(2-2k)\sum_m m^2\binom{l+n}{m} \\
&\qquad\quad + u(1-2k+k^2)\sum_m m\binom{l+n}{m} \\
&\qquad\quad + v\sum_m m^3\binom{l+n}{m} + v(-2-2k)\sum_m m^2\binom{l+n}{m} \\
&\qquad\quad + v(1+2k+k^2)\sum_m m\binom{l+n}{m}\Bigg]
\end{aligned} \tag{S47}
$$

The final simplified form is:

$$
\begin{aligned}
\sum_m (m-k)^2[R_0+R_+ +R_-] = \frac{1}{\bar{w}(x)^2}\sum_{l,n} x_l x_n w(l)w(n)\Bigg[&(1+2u-2v)\sum_m m^2\binom{l+n}{m}2^{-(l+n)} \\
&+(u+v-2k(1+u-v))\left(\frac{l+n}{2}\right)+k^2\Bigg]
\end{aligned} \tag{S48}
$$

Substituting Eqs. (S35), (S36) and (S48) into the generator in Eq. (S33) and combining terms, we obtain:

$$
\begin{aligned}
G_1(g(\sigma^2(x))) = \frac{N}{2}\sum_k x_k \frac{g'(\sigma^2(x))}{\bar{w}(x)^2}\sum_{l,n} x_l x_n w(l)w(n)\Bigg[&(1+2u-2v\sum_m m^2\binom{l+n}{m}2^{-(l+n)} \\
+(u+v)\left(\frac{l+n}{2}\right) - k^2 - 2\mu(x)(1+u-v)\left(\frac{l+n}{2}\right)+2\mu(x)k\Bigg]& \\
-\frac{1}{2}\sum_k x_k\frac{g'(\sigma^2(x))}{\bar{w}(x)^2}\sum_{l,n} x_l x_n w(l)w(n)\Bigg[&(1+2u-2v)\sum_m m^2\binom{l+n}{m}2^{-(l+n)} \\
+(u+v-2k(1+u-v))\left(\frac{l+n}{2}\right)+k^2\Bigg]&
\end{aligned} \tag{S49}
$$

109 Combining terms with the same coefficients:

$$
\begin{aligned}
G_1(g(\sigma^2(x))) = &\frac{(N-1)}{2\bar{w}(x)^2} g'(\sigma^2(x)) \sum_{k,l,n} x_k x_l x_n w(l) w(n) (1+2u-2v) \sum_m m^2 \binom{l+n}{m} 2^{-(l+n)} \\
&+ \frac{N}{2\bar{w}(x)^2} g'(\sigma^2(x)) \sum_{k,l,n} x_k x_l x_n w(l) w(n) \Big[ (u+v) - 2\mu(x)(1+u-v) \Big] \left( \frac{l+n}{2} \right) \\
&- \frac{1}{2\bar{w}(x)^2} g'(\sigma^2(x)) \sum_{k,l,n} x_k x_l x_n w(l) w(n) \Big[ (u+v) - 2k(1+u-v) \Big] \left( \frac{l+n}{2} \right) \\
&+ \frac{N}{2\bar{w}(x)^2} g'(\sigma^2(x)) \sum_{k,l,n} x_k x_l x_n w(l) w(n) \Big[ -k^2 + 2\mu(x) k \Big] \\
&- \frac{1}{2\bar{w}(x)^2} g'(\sigma^2(x)) \sum_{k,l,n} x_k x_l x_n w(l) w(n) k^2 \ .
\end{aligned}
\tag{S50}
$$

110 The generator $G_1(g(\sigma^2(x)))$ simplifies by summation over $k$:

$$
\begin{aligned}
G_1(g(\sigma^2(x))) = &\, g'(\sigma^2(x)) \frac{(N-1)}{2\bar{w}(x)^2} (1+2u-2v) \sum_{l,n} x_l x_n w(l) w(n) \sum_m m^2 \binom{l+n}{m} 2^{-(l+n)} \\
&+ g'(\sigma^2(x)) \frac{(N-1)}{2\bar{w}(x)^2} \sum_{l,n} x_l x_n w(l) w(n) \mathbb{E}[M|l,n] \Big[ (u+v) - 2\mu(x)(1+u-v) \Big] \\
&+ g'(\sigma^2(x)) \frac{1}{2\bar{w}(x)^2} \sum_{l,n} x_l x_n w(l) w(n) \Big[ -(N+1)\psi(x) + 2N\mu(x)^2 \Big] \ .
\end{aligned}
\tag{S51}
$$

111 Simplifying the last term, we find

$$
\begin{aligned}
-(N+1)\psi(x) + 2N\mu(x)^2 &= -(N+1)(\sigma^2(x) + \mu(x)^2) + 2N\mu(x)^2 \\
&= -(N+1)\sigma^2(x) + (N-1)\mu(x)^2 \ .
\end{aligned}
\tag{S52}
$$

The generator then becomes

$$
\begin{aligned}
G_1(g(\sigma^2(x))) = &\, g'(\sigma^2(x)) \frac{(N-1)}{2\bar{w}(x)^2} (1+2u-2v) \sum_{l,n} x_l x_n w(l) w(n) \sum_m m^2 \binom{l+n}{m} 2^{-(l+n)} \\
&+ g'(\sigma^2(x)) \frac{(N-1)}{2\bar{w}(x)^2} \sum_{l,n} x_l x_n w(l) w(n) \left( \frac{l+n}{2} \right) \Big[ (u+v) - 2\mu(x)(1+u-v) \Big] \\
&+ g'(\sigma^2(x)) \frac{1}{2\bar{w}(x)^2} \sum_{l,n} x_l x_n w(l) w(n) \Big[ -(N+1)\sigma^2(x) + (N-1)\mu(x)^2 \Big] \\
= &\, g'(\sigma^2(x)) \frac{(N-1)}{2\bar{w}(x)^2} \sum_{l,n} x_l x_n w(l) w(n) \Big[ (1+2u-2v) \sum_m m^2 \binom{l+n}{m} 2^{-(l+n)} \\
&+ \left( \frac{l+n}{2} \right) \Big( (u+v) - 2\mu(x)(1+u-v) \Big) + \Big( -\frac{N+1}{N-1} \sigma^2(x) + \mu(x)^2 \Big) \Big] \ .
\end{aligned}
\tag{S53}
$$

For large population size, $(N+1)/(N-1) \approx 1$ and $(N-1) \approx N$, and inserting the binomial second moment and mean, we have

$$\begin{aligned} G_1(g(\sigma^2(x))) \approx g'(\sigma^2) \frac{N}{2\bar{w}(x)^2} \Bigg[ &(1+2u-2v) \sum_{l,n} x_l x_n w(l) w(n) \left( \frac{l+n}{4} + \frac{(l+n)^2}{4} \right) \\ &+ (u+v-2\mu(x)(1+u-v)) \sum_{l,n} x_l x_n w(l) w(n) \left( \frac{l+n}{2} \right) \\ &+ \left( \mu(x)^2 - \sigma(x)^2 \right) \sum_{l,n} x_l x_n w(l) w(n) \Bigg] . \end{aligned} \tag{S54}$$

We now solve each line separately.

**First line in Eq. (S54):**

Using Eq. (S19) for $w(l)w(n)$ and by symmetry of $l, n$, we have

$$(1+2u-2v) w(\mu) \sum_{l,n} x_l x_n \left[ w(\mu) + 2w'(\mu)(n-\mu) + w''(\mu)(n-\mu)^2 \right] \left( \frac{l+n}{4} + \frac{(l+n)^2}{4} \right) . \tag{S55}$$

*Terms with* $w(\mu)$: For the zero-th order derivative terms in $w$ we have

$$\sum_{l,n} x_l x_n \left( \frac{l+n}{4} + \frac{(l+n)^2}{4} \right) = \frac{\mu}{2} + \frac{1}{4} \sum_{l,n} x_l x_n (l^2 + 2ln + n^2) = \frac{\mu}{2} + \frac{1}{2}\sigma^2 + \mu^2 . \tag{S56}$$

*Terms with* $w'(\mu)$: For the terms with the first derivative of $w$, we have

$$\begin{aligned} 2 \sum_{l,n} x_l x_n (n-\mu) \left( \frac{l+n}{4} + \frac{(l+n)^2}{4} \right) &= \frac{1}{2} \sum_{l,n} x_l x_n \left( nl + n^2 - \mu(l+n) + nl^2 + 2ln^2 + n^3 - \mu(l^2 + 2ln + n^2) \right) \\ &= \frac{1}{2} \Big( \mu^2 + \sigma^2 + \mu^2 - 2\mu^2 + \mu(\sigma^2 + \mu^2) + 2\mu(\sigma^2 + \mu^2) \\ &\quad + \sigma^3 \gamma + 3\mu\sigma^2 + \mu^3 - \mu(\sigma^2 + \mu^2) - 2\mu^3 - \mu(\sigma^2 + \mu^2) \Big) \\ &= \frac{1}{2} \left( \sigma^2 + 4\mu\sigma^2 + \sigma^3 \gamma \right) , \end{aligned} \tag{S57}$$

where we used the identity $\mathbb{E}[X^3] = \sigma^3\gamma + 3\mu\sigma^2 + \mu^3$.

*Terms with $w''(\mu)$*: The terms corresponding to the second derivative of $w$ simplify to

$$\begin{aligned}
\sum_{l,n} x_l x_n (n-\mu)^2 \left(\frac{l+n}{4} + \frac{(l+n)^2}{4}\right) &= \frac{1}{4}\sum_{l,n} x_l x_n \left(n^2 l + n^3 - 2\mu n(l+n) + \mu^2(l+n) + (n-\mu)^2(l+n)^2\right) \\
&= \frac{1}{4}\Big[\mu(\sigma^2+\mu^2) + \sigma^3\gamma + 3\mu\sigma^2 + \mu^3 - 2\mu^3 - 2\mu(\sigma^2+\mu^2) + 2\mu^3 \\
&\quad + \sum_{l,n} x_l x_n \Big(n^2 l^2 + 2n^3 l + n^4 - 2\mu(nl^2 + 2ln^2 + n^3) \\
&\qquad + \mu^2(l^2 + 2nl + n^2)\Big)\Big] \\
&= \frac{1}{4}\Big(\sigma^3\gamma + 2\mu\sigma^2 \\
&\qquad + (\sigma^2+\mu^2)^2 + 2\mu\sigma^3\gamma + 6\mu^2\sigma^2 + 2\mu^4 + \\
&\qquad + \sigma^4\kappa + 4\mu\sigma^3\gamma + 12\mu^2\sigma^2 + 4\mu^4 - 6\mu^2(\mu^2+\sigma^2) + 3\mu^4 \\
&\qquad - 2\mu^2(\sigma^2+\mu^2) - 4\mu^2(\sigma^2+\mu^2) \\
&\qquad - 2\mu\sigma^3\gamma - 6\mu^2\sigma^2 - 2\mu^4 + 2\mu^2(\sigma^2+\mu^2) + 2\mu^4\Big) \\
&= \frac{1}{4}\left(\sigma^4(1+\kappa) + \sigma^3\gamma(1+4\mu) + 2\mu\sigma^2 + 4\mu^2\sigma^2\right) \ ,
\end{aligned} \tag{S58}$$

where the fourth moment can be rewritten as $\mathbb{E}[X^4] = \sigma^4\kappa + 4\mu\mathbb{E}[X^3] - 6\mu^2\mathbb{E}[X^2] + 3\mu^4$.

**Second line in Eq. (S54):**

Using Eq. (S19) for $w(l)w(n)$ and by symmetry of $l, n$, we have

$$(u + v - 2\mu(1+u-v))\, w(\mu) \sum_{l,n} x_l x_n \left(w(\mu) + 2w'(\mu)(l-\mu) + w''(\mu)(l-\mu)^2\right)\left(\frac{l+n}{2}\right) \tag{S59}$$

We resolve the sum as follows

$$\begin{aligned}
w(\mu)\mu &+ w'(\mu)\sum_{l,n} x_l x_n (l^2 + ln - \mu l - \mu n) + \frac{w''(\mu)}{2}\sum_{l,n} x_l x_n (l^3 + l^2 n - 2\mu(l^2 + ln) + \mu^2(l+n)) \\
&= w(\mu)\mu + w'(\mu)(\sigma^2 + \mu^2 + \mu^2 - 2\mu^2) \\
&\quad + \frac{w''(\mu)}{2}\left(\sigma^3\gamma + 3\mu\sigma^2 + \mu^3 + \mu(\sigma^2+\mu^2) - 2\mu(\sigma^2+\mu^2) - 2\mu^3 + 2\mu^3\right) \\
&= w(\mu)\mu + w'(\mu)\sigma^2 + \frac{w''(\mu)}{2}\left(\sigma^3\gamma + 2\mu\sigma^2\right)
\end{aligned} \tag{S60}$$

**Third line in Eq. (S54):**

Here, we have

$$(\mu^2 - \sigma^2) w(\mu) \sum_{l,n} x_l x_n (w(\mu) + 2w'(\mu)(l-\mu) + w''(\mu)(l-\mu)^2) = (\mu^2 - \sigma^2) w(\mu)\left(w(\mu) + w''(\mu)\sigma^2\right) \tag{S61}$$

Collecting terms, ignoring products of $u, v$ and derivatives of $w$, rewriting the selection function in terms of logarithmic derivatives (Eq. (S25), and inserting into Eq. (S54), we find

$$
\begin{aligned}
G_1(g(\sigma^2(x)) &\approx g'(\sigma^2)\frac{Nw(\mu)}{2\bar{w}^2}\left[w(\mu)\left((1+2u-2v)\left(\frac{\mu+\sigma^2}{2}+\mu^2\right)+(u+v-2\mu(1+u-v))\mu+\mu^2-\sigma^2\right)\right.\\
&\quad +w'(\mu)\left(\frac{\sigma^2}{2}+2\mu\sigma^2+\frac{\sigma^3\gamma}{2}-2\mu\sigma^2\right)\\
&\quad \left.+w''(\mu)\left(\frac{\sigma^4(1+\kappa)}{4}+\frac{\sigma^3\gamma}{4}+\sigma^3\gamma\mu+\frac{\mu\sigma^2}{2}+\mu^2\sigma^2-\mu\sigma^3\gamma-2\mu^2\sigma^2+\mu^2\sigma^2-\sigma^4\right)\right]\\
&= g'(\sigma^2)\frac{Nw(\mu)}{2\bar{w}^2}\left[\frac{w(\mu)}{2}\Big(\mu+4u\mu+\sigma^2(2u-2v-1)\Big)+\frac{w'(\mu)}{2}\sigma^2(1+\sigma\gamma)\right.\\
&\quad \left.\frac{w''(\mu)}{4}\sigma^2(2\mu+\sigma\gamma+\sigma^2(\kappa-3))\right]\\
&\approx g'(\sigma^2)\frac{Nw(\mu)^2}{2w(\mu)^2(1+\beta_2\sigma^2)}\left[\mu\left(\frac{1}{2}+2u\right)+\sigma^2\left(u-v-\frac{1}{2}\right)+\frac{\beta_1}{2}\sigma^2(1+\sigma\gamma)\right.\\
&\quad \left.+\frac{\beta_2}{4}\sigma^2\Big(2\mu+\sigma\gamma+\sigma^2(\kappa-3)\Big)\right]\\
&\approx \frac{g'(\sigma^2N}{2}\left[\mu\left(\frac{1}{2}+2u\right)+\sigma^2\left(u-v-\frac{1}{2}(1-\beta_1-\beta_2\mu)\right)+\frac{\gamma\sigma^3}{4}(2\beta_1+\beta_2)+\frac{\sigma^4}{4}\beta_2(\kappa-3)\right]\ .
\end{aligned}
\tag{S62}
$$

Rescaling time by $N/2$, we find the following differential equation:

$$
\begin{aligned}
\frac{d}{dt}\sigma^2(x) = \left(2u+\frac{1}{2}\right)\mu+\left(u-v-\frac{1}{2}\right)\sigma^2+\frac{1}{2}\left(\beta_1+\beta_2\mu\right)\sigma^2\\
+\frac{1}{4}\left(2\beta_1+\beta_2\right)\gamma\sigma^3+\frac{1}{4}(\kappa-3)\beta_2\sigma^4\ .
\end{aligned}
\tag{S63}
$$

#### S2 Parameteric (negative binomial) closure of dispersion parameter $p_{\mathsf{nb}}$

##### S2.1 Solution of $p_{\mathsf{nb}}$

From the main text (Eq. (12)), the mean and variance at equilibrium are given by

$$(u-v)\mu_{\mathsf{nb}} + \sigma^2_{\mathsf{nb}}\beta_1 + \frac{1}{2}\vartheta\sigma^2_{\mathsf{nb}}\beta_2 = 0, \tag{S64}$$

$$\begin{aligned}\left(2u+\frac{1}{2}\right)\mu_{\mathsf{nb}} + \left(u-v-\frac{1}{2}\right)\sigma^2_{\mathsf{nb}} + \frac{1}{2}(\beta_1+\beta_2\mu_{\mathsf{nb}})\sigma^2_{\mathsf{nb}} \\ + \frac{1}{4}(2\beta_1+\beta_2)\vartheta\sigma^2_{\mathsf{nb}} + \frac{1}{4}\alpha\beta_2\sigma^2_{\mathsf{nb}} = 0.\end{aligned} \tag{S65}$$

Subtracting Eq. (S65) from Eq. (S64) eliminates shared terms and yields:

$$\left(-u-v-\frac{1}{2}\right)\mu_{\mathsf{nb}} + \left(-u+v+\frac{1}{2}(1+\beta_1-\beta_2\mu_{\mathsf{nb}})\right)\sigma^2_{\mathsf{nb}} + \frac{1}{2}\left(-\beta_1+\frac{1}{2}\beta_2\right)\rho\sigma^2_{\mathsf{nb}} - \frac{1}{4}\alpha\beta_2\sigma^2_{\mathsf{nb}} = 0. \tag{S66}$$

Defining $p_{\mathsf{nb}} = \mu_{\mathsf{nb}}/\sigma^2_{\mathsf{nb}}$, inserting the expressions of $\vartheta = \frac{2-p_{\mathsf{nb}}}{p_{\mathsf{nb}}}$ and $\alpha = \frac{6(1-p_{\mathsf{nb}})}{p^2_{\mathsf{nb}}} + 1$, and dividing by $\sigma^2_{\mathsf{nb}}$ gives:

$$\left(-u-v-\frac{1}{2}\right)p_{\mathsf{nb}} = \left(u-v-\frac{1}{2}(1+\beta_1-\beta_2\mu_{\mathsf{nb}})\right) + \frac{2-p_{\mathsf{nb}}}{2p_{\mathsf{nb}}}\left(\beta_1-\frac{1}{2}\beta_2\right) + \frac{1}{4}\beta_2\left(\frac{6(1-p_{\mathsf{nb}})+p^2_{\mathsf{nb}}}{p^2_{\mathsf{nb}}}\right). \tag{S67}$$

Multiplying both sides by $p^2_{\mathsf{nb}}$ gives:

$$\begin{aligned}\left(-u-v-\frac{1}{2}\right)p^3_{\mathsf{nb}} = \left(u-v-\frac{1}{2}(1+\beta_1-\beta_2\mu_{\mathsf{nb}})\right)p^2_{\mathsf{nb}} \\ + \frac{1}{2}\left(\beta_1-\frac{1}{2}\beta_2\right)(2-p_{\mathsf{nb}})p_{\mathsf{nb}} + \frac{1}{4}\beta_2\left[6(1-p_{\mathsf{nb}})+p^2_{\mathsf{nb}}\right].\end{aligned} \tag{S68}$$

Expanding and simplifying leads to the cubic form:

$$0 = \left(u+v+\frac{1}{2}\right)p^3_{\mathsf{nb}} + \left(u-v-\beta_1-\frac{1}{2}\left(1-\beta_2\mu_{\mathsf{nb}}-\frac{1}{2}\beta_2\right)\right)p^2_{\mathsf{nb}} + (\beta_1-2\beta_2)p_{\mathsf{nb}} + \frac{3}{2}\beta_2. \tag{S69}$$

This gives a cubic equation in $p_{\mathsf{nb}}$ of the form:

$$Ap^3_{\mathsf{nb}} + Bp^2_{\mathsf{nb}} + Cp_{\mathsf{nb}} + D = 0, \tag{S70}$$

with coefficients:

$$\begin{aligned}A &= u+v+\frac{1}{2}, \\ B &= \left(u-v-\frac{1}{2}-\beta_1+\frac{1}{2}\beta_2\mu_{\mathsf{nb}}+\frac{1}{2}\beta_2\right), \\ C &= (\beta_1-2\beta_2), \\ D &= \frac{3}{2}\beta_2.\end{aligned} \tag{S71}$$

The real solution for $p_{\mathsf{nb}}$ is given by (see Section S2.1.1 for a justification):

$$p_{\mathsf{nb}} = d + e - \frac{a}{3}, \tag{S72}$$

where

$$\begin{aligned} s &= \sqrt{\frac{Q^2}{4} + \frac{P^3}{27}}, \\ d &= \sqrt[3]{-\frac{Q}{2} + s}, \\ e &= \sqrt[3]{-\frac{Q}{2} - s}. \end{aligned} \tag{S73}$$

The parameters $a$, $P$, and $Q$ are computed from Eq. (S71) as:

$$\begin{aligned} a &= \frac{B}{A} = \frac{u - v - \frac{1}{2} - \beta_1 + \frac{1}{2}\beta_2\mu_{\mathsf{nb}} + \frac{1}{2}\beta_2}{u + v + \frac{1}{2}}, \\ P &= \frac{C}{A} - \frac{a^2}{3} = \frac{\beta_1 - 2\beta_2}{u + v + \frac{1}{2}} - \frac{a^2}{3}, \\ Q &= \frac{D}{A} + \frac{2a^3}{27} - \frac{aC}{3A} = \frac{\frac{3}{2}\beta_2}{u + v + \frac{1}{2}} + \frac{2a^3}{27} - \frac{a(\beta_1 - 2\beta_2)}{3(u + v + \frac{1}{2})}. \end{aligned} \tag{S74}$$

##### S2.1.1 Proof of Eq. S72

From the cubic equation given by Eq. (S70), there are three possible cases of solution depending on the discriminant:

$$\Delta = 4P^3 + 27Q^2 \quad \text{(Cardano's formula)}, \tag{S75}$$

- if $\Delta > 0$ (one real root),
- if $\Delta = 0$ (repeated roots),
- if $\Delta < 0$ (three distinct real roots),

where $P$ and $Q$ are given as in Eq. (S74).

Together, Eq. (S72) and Eq. (S73) give $p_{\mathsf{nb}}$ as Cardano's real root for (S70) in depressed form with $P$ and $Q$. Below we show that $\Delta > 0$ under our assumptions; Eq. (S75) then classifies the cubic as having a single real root.

The parameters involved satisfy the following constraints: $\mu > 0$, $u > 0$, $v > 0$, the constants $\beta_1, \beta_2 < 0$, where we additionally assume that $\beta_1 - 2\beta_2 < 0$, and $u - v < \frac{1}{2} + \beta_1$ (realistic biological conditions).

The common positive denominator is given by

$$A = u + v + \frac{1}{2}. \tag{S76}$$

Then, the parameter $a$ is expressed as

$$a = \frac{u - v - \frac{1}{2} - \beta_1 + \frac{1}{2}\beta_2(\mu + 1)}{A}. \tag{S77}$$

To determine the sign of $a$, observe that since $u - v < \frac{1}{2} + \beta_1$, the term $u - v - \frac{1}{2} - \beta_1$ is negative. Moreover, $\beta_2 < 0 \Rightarrow \frac{1}{2}\beta_2(\mu + 1) < 0$. Thus, the numerator is negative. Since $A > 0$, we conclude that $a < 0$.

From Eq. (S74) we have the depressed-cubic coefficients

$$\begin{aligned} P &= \frac{\beta_1 - 2\beta_2}{A} - \frac{a^2}{3}, \\ Q &= \frac{\frac{3}{2}\beta_2}{A} + \frac{2a^3}{27} - \frac{a(\beta_1 - 2\beta_2)}{3A}. \end{aligned} \tag{S78}$$

To deterimine the sign of the discriminant $\Delta = 4P^3 + 27Q^2$ in Eq. (S75), expand $P$ and $Q$ from Eq. (S74) around their dominant $a$-dependent parts by writing

$$P = -\frac{a^2}{3} + p_1, \qquad Q = \frac{2a^3}{27} + q_1, \tag{S79}$$

where

$$p_1 := \frac{\beta_1 - 2\beta_2}{A}, \qquad q_1 := \frac{\frac{3}{2}\beta_2}{A} - \frac{a(\beta_1 - 2\beta_2)}{3A}. \tag{S80}$$

Note that $p_1$ and $q_1$ are the terms proportional to the selection coefficients; in a weak-selection scaling $\beta_1, \beta_2 = O(1/N)$ with $a, A = O(1)$, one has $p_1, q_1 = O(1/N)$.

Expand $P^3$ and $Q^2$ with binomial coefficients:

$$P^3 = \left(-\frac{a^2}{3}\right)^3 + 3\left(-\frac{a^2}{3}\right)^2 p_1 + 3\left(-\frac{a^2}{3}\right) p_1^2 + p_1^3 = -\frac{a^6}{27} + \frac{a^4}{3} p_1 - a^2 p_1^2 + p_1^3, \tag{S81}$$

$$Q^2 = \left(\frac{2a^3}{27}\right)^2 + 2\left(\frac{2a^3}{27}\right) q_1 + q_1^2 = \frac{4a^6}{729} + \frac{4a^3}{27} q_1 + q_1^2. \tag{S82}$$

Therefore

$$\begin{aligned} \Delta &= 4P^3 + 27Q^2 \\ &= 4\left(-\frac{a^6}{27} + \frac{a^4}{3} p_1 - a^2 p_1^2 + p_1^3\right) + 27\left(\frac{4a^6}{729} + \frac{4a^3}{27} q_1 + q_1^2\right) \\ &= \underbrace{\left(-\frac{4a^6}{27} + \frac{4a^6}{27}\right)}_{=0} + \frac{4a^4}{3} p_1 + 4a^3 q_1 + \underbrace{\left(-4a^2 p_1^2 + 4p_1^3 + 27q_1^2\right)}_{=:\mathcal{R}}. \end{aligned} \tag{S83}$$

For small $p_1, q_1$ and bounded $a$, $\mathcal{R} = O(p_1^2 + q_1^2)$ as $p_1, q_1 \to 0$, hence negligible next to the linear terms in $\Delta$.

Factor the linear part as $\Delta = \frac{4a^4}{3} p_1 + 4a^3 q_1 + \mathcal{R} = 4a^3(\frac{a}{3} p_1 + q_1) + \mathcal{R}$. Substituting the definitions of $p_1$ and $q_1$, the two contributions proportional to $a(\beta_1 - 2\beta_2)/A$ cancel:

$$\frac{a}{3} p_1 + q_1 = \frac{a(\beta_1 - 2\beta_2)}{3A} + \left(\frac{\frac{3}{2}\beta_2}{A} - \frac{a(\beta_1 - 2\beta_2)}{3A}\right) = \frac{3\beta_2}{2A}. \tag{S84}$$

Hence the identity

$$\Delta = 4a^3\left(\frac{3\beta_2}{2A}\right) + \mathcal{R} = \frac{6a^3\beta_2}{A} + \mathcal{R}, \tag{S85}$$

with $\mathcal{R} = -4a^2 p_1^2 + 4p_1^3 + 27q_1^2$ as above; we set $\beta_{\max} := \max\{|\beta_1|, |\beta_2|\}$.

Under the assumptions $A > 0$, $a < 0$ (hence $a^3 < 0$), and $\beta_2 < 0$, the leading term follows

$$\frac{6a^3\beta_2}{A} > 0. \tag{S86}$$

If $|\beta_1|$ and $|\beta_2|$ are small enough that $|\mathcal{R}| < \frac{6a^3\beta_2}{A}$ (equivalently, in a weak-selection expansion, $\mathcal{R} = O(\beta_{\max}^2)$), then Eq. (S85) implies

$$\Delta > 0, \tag{S87}$$

so the cubic falls in the one-real-root case of Cardano's formula consistent with Eq. (S75).

#### S2.2 Positive root, $p_{\text{nb}} > 0$

Under $\beta_1 - 2\beta_2 < 0$, $\beta_2 < 0$, and $a < 0$ as above, one checks $P < 0$ (both summands in $P$ are negative) and $Q < 0$ (each summand in $Q$ is negative). The assumption $u - v < \frac{1}{2} + \beta_1$ was used to obtain $a < 0$. We now use $(P, Q, a)$ with these signs to show $d > 0$, $e > 0$, and hence $p_{\text{nb}} > 0$.

We proceed to show that this implies $d = \sqrt[3]{-\frac{Q}{2} + s} > 0$ and $e = \sqrt[3]{-\frac{Q}{2} - s} > 0$. For both $d > 0$ and $e > 0$ to hold simultaneously, we require the arguments of both cube roots to be positive, yielding the conditions

$$-\frac{Q}{2} + s > 0 \tag{S88}$$

and

$$-\frac{Q}{2} - s > 0. \tag{S89}$$

Eq. (S88) is satisfied since $Q < 0$ as shown above and $s > 0$ by definition. Eq. (S89) can be rewritten as follows

$$s < -\frac{Q}{2} \quad \Longleftrightarrow \quad \sqrt{\frac{Q^2}{4} + \frac{P^3}{27}} < -\frac{Q}{2} \quad \Longleftrightarrow \quad \frac{P^3}{27} < 0\,, \tag{S90}$$

which is satisfied since $P < 0$ as shown before.

We conclude that under the assumption $u - v < \frac{1}{2} + \beta_1$, the resulting conditions $P < 0$, $Q < 0$, and $a < 0$ collectively ensure that $d > 0$, $e > 0$, and consequently $p_{\text{nb}} > 0$. Thus, $p_{\text{nb}} > 0$ if and only if the condition

$$u - v < \frac{1}{2} + \beta_1, \tag{S91}$$

is satisfied.

Therefore, the real positive root is given by Equation (S72).

#### S2.3 Poisson case, $p_{\text{nb}} = 1$

For the Poisson case, that is $p_{\text{nb}} = 1$, we have from Eq. (S70):

$$Ap_{\text{nb}}^3 + Bp_{\text{nb}}^2 + Cp_{\text{nb}} + D = A + B + C + D = 0, \tag{S92}$$

where $A$, $B$, $C$, and $D$ are given in Eq. (S71).

Substituting the coefficients $A$, $B$, $C$ and $D$ from Eq. (S71) gives

$$\left(u + v + \frac{1}{2}\right) + \left(u - v - \frac{1}{2} - \beta_1 + \frac{1}{2}\beta_2\mu + \frac{1}{2}\beta_2\right) + (\beta_1 - 2\beta_2) + \frac{3}{2}\beta_2 = 0. \tag{S93}$$

Simplifying, we obtain:

$$2u + \frac{1}{2}\beta_2\mu = 0 \quad \Longleftrightarrow \quad u = -\frac{1}{4}\beta_2\mu \ , \tag{S94}$$

which shows that for the distribution to be Poisson, the transposition rate in equilibrium needs to be equal to the (scaled) product of (a function of the) selection coefficient and mean TE copy number per genome.

##### S2.4 Overdispersion case, $p_{\text{nb}} < 1$

For the condition $p_{\text{nb}} < 1$, we require:

$$p_{\text{nb}} = d + e - \frac{a}{3} < 1. \tag{S95}$$

With $\mu > 0$, $u > 0$, $v > 0$, and $\beta_1, \beta_2 < 0$, we know from Eq. (S94) that the boundary where $p_{\text{nb}} = 1$ is $u = -\frac{1}{4}\beta_2\mu$. This boundary separates two regimes in the parameter space. Since $\beta_2 < 0$ and $\mu > 0$, the expression $-\frac{1}{4}\beta_2\mu$ is positive. When $u$ exceeds the critical value in Eq. (S94), the cubic root $p_{\text{nb}}$ falls below one. Thus:

$$u > -\frac{1}{4}\beta_2\mu. \tag{S96}$$

#### S3 Proofs for equidispersion ($p_{\text{nb}} = 1$) and underdispersion ($p_{\text{nb}} > 1$) compatibility

Throughout this section, $\mu_{\text{nb}}$, $\sigma^2_{\text{nb}}$, $p_{\text{nb}}$, $\hat{\mu}_{\text{app}}$, and $\hat{p}_{\text{app}}$ refer to the standing population distribution, i.e., the TE copy number measured after a complete birth event, following transposition and excision (the quantity denoted $m$ and $\rho$ in the main text). The corresponding pre-transposition/excision quantities, $m'$ and $\rho_{m'}$, measured immediately after recombination but before the current generation's transposition and excision, are treated separately in Section S5.

##### S3.1 Equidispersion ($p_{\text{nb}} = 1$)

**Parametric (negative-binomial) closure.** Equation (S94) (Eq. (26) in the main text) gives the transposition rate $u$ at which the cubic equation for $p_{\text{nb}}$ has $p_{\text{nb}} = 1$ as a root, for a given equilibrium mean $\mu_{\text{nb}}$ and given $\beta_2(\mu_{\text{nb}})$. By itself, it does not decide whether the negative-binomial closed dynamics admit a steady state with $\sigma^2_{\text{nb}} = \mu_{\text{nb}} > 0$. To link the Poisson root to a putative equilibrium, we combine it with the mean equation at equilibrium.

Suppose there were an equilibrium with $\mu_{\text{nb}} > 0$, equidispersion $\sigma^2_{\text{nb}} = \mu_{\text{nb}}$, so $p_{\text{nb}} = 1$ and hence $\vartheta = 1$. Setting Eq. (S64) to zero and dividing by $\mu_{\text{nb}}$ gives

$$(u - v) + \beta_1 + \tfrac{1}{2}\beta_2 = 0. \tag{S97}$$

If in addition $p_{\text{nb}} = 1$ is a root of the cubic, then Eq. (S94) holds. Eliminating $u$ between (S97) and (S94) yields the compatibility condition

$$v = \beta_1 + \frac{\beta_2}{4}(2 - \mu_{\text{nb}}) \quad \text{equivalently} \quad 4(\beta_1 - v) + \beta_2(2 - \mu_{\text{nb}}) = 0. \tag{S98}$$

Defining $r(\mu) = \mu\beta_2(\mu)/\beta_1(\mu)$, we can then rewrite the condition by

$$v = \beta_1 + \frac{\beta_2}{2} - \frac{\beta_2\mu_{\text{nb}}}{4} = \beta_1\left(1 + \frac{r}{2\mu_{\text{nb}}} - \frac{r}{4}\right) \ . \tag{S99}$$

We know that $\beta_1 < 0$ since the strength of purifying selection increases with the number of TE copies. Thus, for condition Eq. (S99) to have a possible value of $v \geq 0$, we need

$$1 + \frac{r}{2\mu_{\text{nb}}} - \frac{r}{4} < 0 \qquad \Leftrightarrow \qquad 1 < r\left(\frac{\mu - 2}{4\mu}\right) \ , \tag{S100}$$

The fraction on the right-hand side is positive for $\mu_{\text{nb}} > 2$, a reasonable assumption for our theory because otherwise the loss of TEs in the population is very likely. Thus, for this inequality to be true, we have $r > 4\mu/(\mu - 2) \approx 4$ for large enough $\mu$.

If we assume that TE costs are dominated by independent per-copy effects and pairwise TE interactions (e.g. ectopic recombination), so that locally $g$ is no more than quadratic, $g(\mu) \approx c - a\mu - b\mu^2$ with $a, b \geq 0$, this implies

$$r(\mu) := \frac{\mu\beta_2(\mu)}{\beta_1(\mu)} = \frac{2b\mu}{a + 2b\mu} \in [0, 1]. \tag{S101}$$

Alternatively, and more generally, we allow mildly super-quadratic curvature at the relevant mean by assuming that for some small $\delta \geq 0$,

$$0 \leq r(\mu_{\text{nb}}) \leq 1 + \delta. \tag{S102}$$

Thus, if $r \leq 1 + \delta < 4$, then the bracket is positive, so $v$ has the sign of $\beta_1$ and thus $v < 0$. Hence under these conditions there is no Poisson-compatible equilibrium with $v \geq 0$.

**Approximate closure.** In the approximate closure (Eq. (25) in the main text), equidispersion imposes

$$4u = \phi(\hat{\mu}_{\text{app}}). \tag{S103}$$

Write $g(\mu) := \ln w(\mu)$ so $\beta_1 = g'(\mu)$ and $\beta_2 = g''(\mu)$, and define

$$\phi(\mu) := \beta_1(\mu) - \mu\beta_2(\mu), \qquad r(\mu) := \frac{\mu\beta_2(\mu)}{\beta_1(\mu)}. \tag{S104}$$

Assume $\hat{\mu}_{\text{app}} > 0$, $\beta_1(\hat{\mu}_{\text{app}}) < 0$, $\beta_2(\hat{\mu}_{\text{app}}) \leq 0$, and $0 \leq r(\hat{\mu}_{\text{app}}) \leq 1 + \delta$. Then

$$\phi(\hat{\mu}_{\text{app}}) = \beta_1(\hat{\mu}_{\text{app}})(1 - r(\hat{\mu}_{\text{app}})) \leq |\beta_1(\hat{\mu}_{\text{app}})|\,\delta, \qquad \Rightarrow \qquad u = \frac{\phi(\hat{\mu}_{\text{app}})}{4} \leq \frac{|\beta_1(\hat{\mu}_{\text{app}})|\,\delta}{4}, \tag{S105}$$

Again, in the strictly pairwise-limited case where TE costs are dominated by independent per-copy effects and pairwise TE interactions (e.g. ectopic recombination), so that locally $g$ is no more than quadratic, $g(\mu) \approx c - a\mu - b\mu^2$ with $a, b \geq 0$, we have $\delta = 0$. Then, the condition reduces to $u \leq 0$, leading to a contradiction that equidispersion cannot occur for active TEs. In general, under weak selection, the equidispersion constraint implies that the transposition rate must be extremely small whenever the curvature excess $\delta$ is not too large (mildly super-quadratic curvature); this makes successful invasion unlikely in biologically realistic parameter regimes.

#### S3.2 Underdispersion (approximate closure)

In the approximate closure (Eq. (25) in the main text), underdispersion corresponds to $\rho_{\mathsf{app}} < 1$ (equivalently $p_{\mathsf{app}} > 1$) and hence reverses the overdispersion condition to

$$4u < \phi(\hat{\mu}_{\mathsf{app}}), \qquad \phi(\mu) := \beta_1(\mu) - \mu\beta_2(\mu). \tag{S106}$$

Assume

$$\hat{\mu}_{\mathsf{app}} > 0, \qquad \beta_1(\hat{\mu}_{\mathsf{app}}) < 0, \qquad \beta_2(\hat{\mu}_{\mathsf{app}}) \leq 0, \qquad 0 \leq r(\hat{\mu}_{\mathsf{app}}) \leq 1 + \delta, \tag{S107}$$

where $r(\mu) := \mu\beta_2(\mu)/\beta_1(\mu)$ and the bound $r \leq 1 + \delta$ allows mildly super-quadratic curvature at the equilibrium mean. Then

$$\phi(\hat{\mu}_{\mathsf{app}}) = \beta_1(\hat{\mu}_{\mathsf{app}})(1 - r(\hat{\mu}_{\mathsf{app}})) \leq |\beta_1(\hat{\mu}_{\mathsf{app}})|\,\delta. \tag{S108}$$

Hence $4u < \phi(\hat{\mu}_{\mathsf{app}}) \leq |\beta_1(\hat{\mu}_{\mathsf{app}})|\,\delta$, so

$$u < \frac{|\beta_1(\hat{\mu}_{\mathsf{app}})|\,\delta}{4}. \tag{S109}$$

In particular, underdispersion with $u > 0$ forces $\phi(\hat{\mu}_{\mathsf{app}}) > 0$ and hence $r(\hat{\mu}_{\mathsf{app}}) > 1$. In the strictly pairwise-limited case $\delta = 0$, this reduces to the contradiction $u \leq 0$.

However, we cannot give an analogous underdispersion proof for the parametric negative-binomial closure because that derivation only supports equidispersion and overdispersion (it enforces $0 < p_{\mathsf{nb}} \leq 1$, i.e. $\sigma^2_{\mathsf{nb}} \geq \mu_{\mathsf{nb}}$).

### S4 Closed-form equilibrium under Gaussian fitness function

For the Gaussian fitness function $w(n) = \exp(-sn^2)$ the equilibrium can be obtained in closed form. Replacing the variance by the dispersion ratio, $\sigma^2_{\mathsf{nb}} = \mu_{\mathsf{nb}}\rho_{\mathsf{nb}}$ with $\rho_{\mathsf{nb}} = 1/p_{\mathsf{nb}}$. And substituting the negative-binomial shape terms $\vartheta = 2\rho_{\mathsf{nb}} - 1$ and $\alpha = 6\rho_{\mathsf{nb}}(\rho_{\mathsf{nb}} - 1) + 1$ into Eq. (12) closes the dynamics in $(\mu_{\mathsf{nb}}, \rho_{\mathsf{nb}})$,

$$\begin{aligned} \frac{d}{dt}\mu_{\mathsf{nb}} &= (u - v)\mu_{\mathsf{nb}} + \mu_{\mathsf{nb}}\rho_{\mathsf{nb}}\beta_1 + \tfrac{1}{2}(2\rho_{\mathsf{nb}} - 1)\mu_{\mathsf{nb}}\rho_{\mathsf{nb}}\beta_2 \ , \\ \frac{d}{dt}\rho_{\mathsf{nb}} &= \tfrac{1}{2}\Big[1 - \rho_{\mathsf{nb}} + 4u + \rho_{\mathsf{nb}}(\mu_{\mathsf{nb}} - \rho_{\mathsf{nb}} + \rho^2_{\mathsf{nb}})\beta_2\Big] \ . \end{aligned} \tag{S110}$$

For $w(n) = \exp(-sn^2)$ we have $\beta_1 = -2s\mu_{\mathsf{nb}}$ and $\beta_2 = -2s$. Setting the right-hand sides of Eq. (S110) to zero and dividing the first by $\hat{\mu}_{\mathsf{nb}} > 0$ gives the two equilibrium conditions

$$2s\hat{\mu}_{\mathsf{nb}}\hat{\rho}_{\mathsf{nb}} = (u - v) - 2s\hat{\rho}^2_{\mathsf{nb}} + s\hat{\rho}_{\mathsf{nb}} \ , \tag{S111}$$

$$\hat{\rho}_{\mathsf{nb}} = 1 + 4u - 2s\hat{\rho}_{\mathsf{nb}}\hat{\mu}_{\mathsf{nb}} + 2s\hat{\rho}^2_{\mathsf{nb}} - 2s\hat{\rho}^3_{\mathsf{nb}} \ . \tag{S112}$$

**Dispersion ratio.** Substituting Eq. (S111) into Eq. (S112) through the product $s\hat{\rho}_{\mathsf{nb}}\hat{\mu}_{\mathsf{nb}}$ yields

$$\hat{\rho}_{\mathsf{nb}} = 1 + 3u + v + s\big(4\hat{\rho}^2_{\mathsf{nb}} - \hat{\rho}_{\mathsf{nb}} - 2\hat{\rho}^3_{\mathsf{nb}}\big) \ . \tag{S113}$$

Equation (S113) is exact, but implicit. Writing $\hat{\rho}_{\mathsf{nb}} = 1+\varepsilon$ and expanding $f(\hat{\rho}_{\mathsf{nb}}) = 4\hat{\rho}^2_{\mathsf{nb}} - \hat{\rho}_{\mathsf{nb}} - 2\hat{\rho}^3_{\mathsf{nb}}$ about $\hat{\rho}_{\mathsf{nb}} = 1$,

$$f(1+\varepsilon) = (4-1-2) + (8-1-6)\,\varepsilon - 2\varepsilon^2 + O(\varepsilon^3) = 1 + \varepsilon - 2\varepsilon^2 + O(\varepsilon^3)\ , \tag{S114}$$

so that $\varepsilon = 3u + v + s\,(1 + \varepsilon - 2\varepsilon^2 + \dots)$. Since $\varepsilon = 3u + O(u^2)$ at leading order, the terms $s\varepsilon$ and $s\varepsilon^2$ which are of order $su$ and $su^2$ are neglected, that is, one and two powers of $\varepsilon$ beyond those retained, and are therefore negligible whenever $u \ll 1$. This gives

$$\hat{\rho}_{\mathsf{nb}} = 1 + 3u + v + s + O(su)\ . \tag{S115}$$

The order-$s$ term originates solely from the skewness and excess-kurtosis contributions: setting $\vartheta = \alpha = 0$ in Eq. (12) of the main text, which removes them while leaving selection everywhere else, gives $\rho = (1 + 2(u+v))/(1 - 2(u-v)) \approx 1 + 4u$. This is independent of $s$, and is exactly the approximate-closure result of Eq. (6) of the main text. Selection curvature therefore lowers the dispersion ratio from $1 + 4u$ to $1 + 3u$ by exactly the net transposition rate. Because $v$ and $s$ enter Eq. (S115) additively, they may be dropped together when $v, s \ll u$, giving the leading-order estimate $\hat{\rho}_{\mathsf{nb}} \approx 1 + 3u$.

**Mean and variance.** With $\hat{\rho}_{\mathsf{nb}}$ known, the mean follows by rearranging Eq. (S111), and the variance is recovered from the definition of the dispersion ratio,

$$\hat{\mu}_{\mathsf{nb}} = \frac{u-v}{2s\,\hat{\rho}_{\mathsf{nb}}} - \hat{\rho}_{\mathsf{nb}} + \frac{1}{2}\ , \qquad \hat{\sigma}^2_{\mathsf{nb}} = \hat{\rho}_{\mathsf{nb}}\,\hat{\mu}_{\mathsf{nb}} = \frac{u-v}{2s} - \hat{\rho}^2_{\mathsf{nb}} + \frac{\hat{\rho}_{\mathsf{nb}}}{2}\ . \tag{S116}$$

Substituting $\hat{\rho}_{\mathsf{nb}}$ from Eq. (S115), retaining $v$ and dropping only the order-$s$ term, $\hat{\rho}_{\mathsf{nb}} = 1 + 3u + v$, gives

$$\hat{\mu}_{\mathsf{nb}} = \frac{u-v}{2s\,(1+3u+v)} - \frac{1}{2} - 3u - v\ , \qquad \hat{\sigma}^2_{\mathsf{nb}} = (1+3u+v)\,\hat{\mu}_{\mathsf{nb}}\ . \tag{S117}$$

**Skewness and excess kurtosis.** The same equilibrium fixes the two higher moments. From Eqs. (10)–(11) of the main text, $\gamma_{\mathsf{nb}} = \vartheta/\sigma_{\mathsf{nb}}$ and $\kappa_{\mathsf{nb}} - 3 = \alpha/\sigma^2_{\mathsf{nb}}$. Writing them in terms of the dispersion ratio using $p_{\mathsf{nb}} = 1/\rho_{\mathsf{nb}}$ gives

$$\vartheta = \frac{2 - p_{\mathsf{nb}}}{p_{\mathsf{nb}}} = \frac{2}{p_{\mathsf{nb}}} - 1 = 2\rho_{\mathsf{nb}} - 1\ , \qquad \alpha = \frac{6(1-p_{\mathsf{nb}}) + p^2_{\mathsf{nb}}}{p^2_{\mathsf{nb}}} = \frac{6}{p^2_{\mathsf{nb}}} - \frac{6}{p_{\mathsf{nb}}} + 1 = 6\rho_{\mathsf{nb}}(\rho_{\mathsf{nb}} - 1) + 1\ . \tag{S118}$$

Substituting the equilibrium dispersion ratio for $s \ll u$, $\hat{\rho}_{\mathsf{nb}} = 1 + \varepsilon$ with $\varepsilon = 3u + v$ yields,

$$\vartheta = 1 + 2\varepsilon = 1 + 6u + 2v\ , \qquad \alpha = 1 + 6\varepsilon + 6\varepsilon^2 = 1 + 6(3u+v) + 6(3u+v)^2\ , \tag{S119}$$

so that, with $\hat{\sigma}^2_{\mathsf{nb}}$ from Eq. (S117),

$$\hat{\gamma}_{\mathsf{nb}} = \frac{1 + 6u + 2v}{\hat{\sigma}_{\mathsf{nb}}}\ , \qquad \hat{\kappa}_{\mathsf{nb}} - 3 = \frac{1 + 6(3u+v) + 6(3u+v)^2}{\hat{\sigma}^2_{\mathsf{nb}}}\ . \tag{S120}$$

#### S5 Pre-transposition/excision mean, variance, and dispersion condition

This section first derives the mean and variance of the pre-transposition/excision copy number $m'$ for a general weak-selection fitness function $w(\cdot)$, matching Eqs. (17)–(20) of the main text, and then uses those expressions to derive a general condition for whether $m'$ itself is over-, equi- or underdispersed, $\rho_{m'} = \mathrm{Var}[m']/E[m'] \gtrless 1$, under the negative-binomial and approximate closures.

##### S5.1 Derivation of the pre-transposition/excision mean and variance

As introduced in Section 1, the pre-transposition/excision copy number of an offspring drawn from parents with $l$ and $n$ TE copies is binomially distributed, reflecting free recombination: each of the $l+n$ parental copies segregates into the offspring independently with probability $\frac{1}{2}$,

$$m' \mid l, n \;\sim\; \mathrm{Binomial}(l+n, \tfrac{1}{2}). \tag{S121}$$

Within a single reproduction event of the Moran process, two parents are sampled according to fitness and $m'$ is the offspring's copy number immediately after segregation, before transposition and excision act on it. At a fixed population state $x$, its mean and variance therefore follow from Eq. (S121) together with the fitness-weighted distribution of the parents, whereas $\mu(x)$ and $\sigma^2(x)$ change as reproduction events accumulate and require the generator of the process.

Since $m' \mid l, n \sim \mathrm{Binomial}(l+n, \frac{1}{2})$,

$$E[m' \mid l, n] = \frac{l+n}{2}, \qquad \mathrm{Var}[m' \mid l, n] = \frac{l+n}{4}. \tag{S122}$$

The parents' copy numbers $l, n$ are two independent draws from the fitness-weighted distribution underlying Eq. (S3), $P(l = k) = x_k w(k)/\bar{w}(x)$. Their exact mean and variance are

$$a := E_w[l] = \sum_k k\,\frac{x_k w(k)}{\bar{w}(x)} = \frac{E[l\,w(l)]}{E[w(l)]}, \qquad b := \mathrm{Var}_w[l] = \sum_k k^2\,\frac{x_k w(k)}{\bar{w}(x)} - a^2 = \frac{E[l^2\,w(l)]}{E[w(l)]} - a^2, \tag{S123}$$

where $E[\cdot]$ denotes the unweighted expectation over $x$, e.g. $E[l\,w(l)] = \sum_k k\,x_k\,w(k)$; both parents are identically distributed, so the same $a, b$ apply to $n$. Equation (S123) is exact, for any fitness function and any population distribution.

By the law of total expectation,

$$E[m'] = E_{l,n}\Big[\frac{l+n}{2}\Big] = \frac{E_w[l] + E_w[n]}{2} = \frac{a+a}{2} = a. \tag{S124}$$

By the law of total variance,

$$\mathrm{Var}[m'] = \underbrace{E_{l,n}\big[\mathrm{Var}[m' \mid l, n]\big]}_{\text{within-pair binomial noise}} + \underbrace{\mathrm{Var}_{l,n}\big[E[m' \mid l, n]\big]}_{\text{between-pair variation}}. \tag{S125}$$

The first term is $E[(l+n)/4] = (a+a)/4 = a/2$. For the second term, since $l, n$ are independent, $\mathrm{Var}[(l+n)/2] = \frac{1}{4}(\mathrm{Var}[l] + \mathrm{Var}[n]) = \frac{1}{4}(b+b) = b/2$. Adding the two terms,

$$\mathrm{Var}[m'] = \frac{a}{2} + \frac{b}{2} = \frac{a+b}{2}. \tag{S126}$$

So

$$E[m'] = a = E_w[l], \qquad \mathrm{Var}[m'] = \frac{a+b}{2} = \frac{E_w[l] + \mathrm{Var}_w[l]}{2}. \tag{S127}$$

Everything reduces to finding $a$ and $b$: the mean and variance of a single parent, drawn with probability proportional to fitness from the population $x$.

Weak selection means $w(\cdot)$ varies slowly enough that $\ln w(l)$ can be Taylor-expanded around the population mean $\mu$ to second order,

$$\ln w(l) \approx \ln w(\mu) + \beta_1(l-\mu) + \frac{\beta_2}{2}(l-\mu)^2, \tag{S128}$$

with $\beta_1, \beta_2$ the log-fitness derivatives already defined in Eq. (S25), both $O(s)$ for a selection strength $s$. Exponentiating and keeping only terms linear in $\beta_1, \beta_2$ (equivalently, linear in $s$; quadratic terms such as $\beta_1^2$ are $O(s^2)$ and dropped),

$$w(l) \;\approx\; w(\mu)\Big[1 + \beta_1(l-\mu) + \frac{\beta_2}{2}(l-\mu)^2\Big]. \tag{S129}$$

Write $X := l - \mu$, so $E[X] = 0$, $E[X^2] = \sigma^2$, $E[X^3] = \gamma\sigma^3$, $E[X^4] = \kappa\sigma^4$ (central moments of the population). The constant $w(\mu)$ cancels between numerator and denominator of $E_w[\cdot] = E[\,\cdot\, w(l)]/E[w(l)]$, so it can be dropped from here on.

We now approximate $a = E_w[l] = E[l\, w(l)]/E[w(l)]$ (Eq. (S123)) to leading order in $s$, using the weak-selection expansion of $w(l)$ above. The denominator is

$$E\big[1 + \beta_1 X + \tfrac{\beta_2}{2}X^2\big] = 1 + \beta_1 \cdot 0 + \frac{\beta_2}{2}\sigma^2 = 1 + \frac{\beta_2}{2}\sigma^2. \tag{S130}$$

With $l = \mu + X$, the numerator is

$$\begin{aligned} E\big[l(1 + \beta_1 X + \tfrac{\beta_2}{2}X^2)\big] &= E\big[(\mu + X)(1 + \beta_1 X + \tfrac{\beta_2}{2}X^2)\big] \\ &= \mu + \mu\beta_1 E[X] + \mu\frac{\beta_2}{2}E[X^2] + E[X] + \beta_1 E[X^2] + \frac{\beta_2}{2}E[X^3] \\ &= \mu + \mu\frac{\beta_2}{2}\sigma^2 + \beta_1\sigma^2 + \frac{\beta_2}{2}\gamma\sigma^3. \end{aligned} \tag{S131}$$

To leading order in $s$, $\big(1 + \frac{\beta_2}{2}\sigma^2\big)^{-1} \approx 1 - \frac{\beta_2}{2}\sigma^2$, so dividing gives

$$E_w[l] \approx \Big[\mu + \mu\frac{\beta_2}{2}\sigma^2 + \beta_1\sigma^2 + \frac{\beta_2}{2}\gamma\sigma^3\Big]\Big[1 - \frac{\beta_2}{2}\sigma^2\Big]. \tag{S132}$$

Expanding and discarding every product of two $O(s)$ factors (e.g. $\beta_1\beta_2\sigma^4$), the $\mu\frac{\beta_2}{2}\sigma^2$ term cancels the corresponding term from the bracket, leaving

$$a = E_w[l] = \mu + \beta_1\sigma^2 + \frac{\beta_2}{2}\gamma\sigma^3 + O(s^2). \tag{S133}$$

Similarly, $b = \mathrm{Var}_w[l] = E_w[l^2] - (E_w[l])^2$ is approximated by first expanding $E_w[l^2] = E[l^2 w(l)]/E[w(l)]$ to leading order in $s$, then subtracting $(E_w[l])^2$. With $l^2 = (\mu+X)^2 = \mu^2 + 2\mu X + X^2$, the numerator of $E_w[l^2]$ is

$$E\big[l^2(1+\beta_1 X+\tfrac{\beta_2}{2}X^2)\big] = E\big[(\mu^2+2\mu X+X^2)(1+\beta_1 X+\tfrac{\beta_2}{2}X^2)\big] \\ = \mu^2+\mu^2\frac{\beta_2}{2}\sigma^2+2\mu\beta_1\sigma^2+2\mu\frac{\beta_2}{2}\gamma\sigma^3+\sigma^2+\beta_1\gamma\sigma^3+\frac{\beta_2}{2}\kappa\sigma^4, \tag{S134}$$

using $E[X]=0$, $E[X^2]=\sigma^2$, $E[X^3]=\gamma\sigma^3$, $E[X^4]=\kappa\sigma^4$ term by term as before. Dividing by the same denominator $1+\frac{\beta_2}{2}\sigma^2$, that is multiplying by $1-\frac{\beta_2}{2}\sigma^2$ and dropping $O(s^2)$ cross-terms, the $\mu^2\frac{\beta_2}{2}\sigma^2$ term again cancels against $-\mu^2\frac{\beta_2}{2}\sigma^2$ from the bracket, but a residual $-\sigma^2\cdot\frac{\beta_2}{2}\sigma^2$ survives, from multiplying the $\sigma^2$ term by the bracket, giving

$$E_w[l^2] = \mu^2+\sigma^2+2\mu\beta_1\sigma^2+\mu\beta_2\gamma\sigma^3+\beta_1\gamma\sigma^3+\frac{\beta_2}{2}(\kappa-1)\sigma^4+O(s^2). \tag{S135}$$

Squaring Eq. (S133), and keeping only $O(1)$ and $O(s)$ terms,

$$(E_w[l])^2 = \big[\mu+\beta_1\sigma^2+\tfrac{\beta_2}{2}\gamma\sigma^3\big]^2 = \mu^2+2\mu\beta_1\sigma^2+\mu\beta_2\gamma\sigma^3+O(s^2). \tag{S136}$$

Subtracting,

$$\mathrm{Var}_w[l] = E_w[l^2]-(E_w[l])^2 = [\mu^2+\sigma^2+2\mu\beta_1\sigma^2+\mu\beta_2\gamma\sigma^3+\beta_1\gamma\sigma^3+\tfrac{\beta_2}{2}(\kappa-1)\sigma^4]-[\mu^2+2\mu\beta_1\sigma^2+\mu\beta_2\gamma\sigma^3], \tag{S137}$$

and every $\mu$-dependent term cancels, leaving

$$b = \mathrm{Var}_w[l] = \sigma^2+\beta_1\gamma\sigma^3+\frac{\beta_2}{2}(\kappa-1)\sigma^4+O(s^2). \tag{S138}$$

Substituting Eqs. (S133) and (S138) (i.e. $a=E_w[l]$, $b=\mathrm{Var}_w[l]$) into Eq. (S127) gives the general pre-transposition/excision mean and variance, matching Eqs. (17)–(18) of the main text:

$$E[m'] = \mu+\beta_1\sigma^2+\frac{1}{2}\beta_2\gamma\sigma^3, \tag{S139}$$

$$\mathrm{Var}[m'] = \frac{\mu+\sigma^2}{2}+\frac{\beta_1}{2}(\sigma^2+\gamma\sigma^3)+\frac{\beta_2}{4}[\gamma\sigma^3+(\kappa-1)\sigma^4], \tag{S140}$$

valid for any weak-selection fitness function $w(\cdot)$ and any population moments $\mu,\sigma^2,\gamma,\kappa$. The pre-transposition/excision dispersion ratio is $\rho_{m'} := \mathrm{Var}[m']/E[m']$; Section S5.4 below evaluates it under the negative-binomial and approximate closures.

#### Skewness and excess kurtosis

The mean and variance of $m'$ are its first two cumulants, and the same construction delivers the next two, using cumulant generating functions (CGF).

Conditional on the parents, $m' \mid l,n \sim \mathrm{Binomial}(S,\frac{1}{2})$ with $S=l+n$, so $m'$ is a sum of $S$ independent Bernoulli($\frac{1}{2}$) variables. A Bernoulli variable $X$ takes the value 1 with probability $p$ and 0 with probability $1-p$, so at $p=\frac{1}{2}$ its moment generating function $M_X$, and hence its cumulant generating function $K_{\mathrm{B}}$, are

$$M_X(t) = E[\mathrm{e}^{tX}] = (1-p)\,\mathrm{e}^{t\cdot 0}+p\,\mathrm{e}^{t\cdot 1} = \frac{1}{2}+\frac{1}{2}\mathrm{e}^t = \frac{1+\mathrm{e}^t}{2}, \qquad K_{\mathrm{B}}(t) = \ln M_X(t) = \ln(1+\mathrm{e}^t)-\ln 2\,. \tag{S141}$$

Its derivatives are

$$K_{\mathrm{B}}'(t) = \frac{\mathrm{e}^t}{1+\mathrm{e}^t} , \qquad K_{\mathrm{B}}''(t) = \frac{\mathrm{e}^t}{(1+\mathrm{e}^t)^2} , \qquad K_{\mathrm{B}}'''(t) = \frac{\mathrm{e}^t(1-\mathrm{e}^t)}{(1+\mathrm{e}^t)^3} , \tag{S142}$$

so at $t = 0$ the first four cumulants are $\frac{1}{2}$, $\frac{1}{4}$, $0$, $-\frac{1}{8}$, the third vanishing because of the factor $1-\mathrm{e}^t$. Equivalently,

$$z := K_{\mathrm{B}}(t) = \frac{t}{2} + \frac{t^2}{8} + 0 \cdot t^3 - \frac{t^4}{192} + O(t^5) . \tag{S143}$$

Because $m'$ is a sum of $S$ independent Bernoulli($\frac{1}{2}$) variables with $S$ itself random, its cumulant generating function is the composition

$$K_{m'}(t) = K_S(K_{\mathrm{B}}(t)) = K_S(z) . \tag{S144}$$

The powers of $z$ needed below are

$$z^2 = \frac{t^2}{4} + \frac{t^3}{8} + \frac{t^4}{64}, \qquad z^3 = \frac{t^3}{8} + \frac{3t^4}{32}, \qquad z^4 = \frac{t^4}{16} . \tag{S145}$$

Write $a, b, c, d$ for the first four cumulants of a single fitness-weighted parent, so that $a = E_w[l]$ and $b = \mathrm{Var}_w[l]$ as above. The two parents are independent and identically distributed, so $\kappa_j(S) = 2\kappa_j(l)$ and

$$K_S(z) = 2a\,z + 2b\,\frac{z^2}{2} + 2c\,\frac{z^3}{6} + 2d\,\frac{z^4}{24} = 2a\,z + b\,z^2 + \frac{c}{3}z^3 + \frac{d}{12}z^4 + O(z^5) . \tag{S146}$$

Substituting Eqs. (S143)–(S145) into Eq. (S146) and collecting powers of $t$,

$$[t^1] = a, \qquad [t^2] = \frac{a+b}{4}, \qquad [t^3] = \frac{b}{8} + \frac{c}{24}, \qquad [t^4] = -\frac{a}{96} + \frac{b}{64} + \frac{c}{32} + \frac{d}{192} , \tag{S147}$$

where, for instance, the $t^4$ term collects $2a$ times the $-1/192$ of $z$, $b$ times the $1/64$ of $z^2$, $\frac{c}{3}$ times the $3/32$ of $z^3$, and $\frac{d}{12}$ times the $1/16$ of $z^4$. Since $\kappa_j(m') = j!\,[t^j]$,

$$\kappa_1(m') = a , \qquad \kappa_2(m') = \frac{a+b}{2} , \qquad \kappa_3(m') = \frac{3b+c}{4} , \qquad \kappa_4(m') = \frac{-2a+3b+6c+d}{8} , \tag{S148}$$

the first two reproducing Eq. (S127).

The fitness weighting leaves the parent cumulants at their population values, $a = \mu$, $b = \sigma^2$, $c = \gamma\sigma^3$ and $d = (\kappa-3)\sigma^4$, so that

$$\kappa_2(m') = \frac{\mu+\sigma^2}{2} , \qquad \kappa_3(m') = \frac{3\sigma^2+\gamma\sigma^3}{4} , \qquad \kappa_4(m') = \frac{-2\mu+3\sigma^2+6\gamma\sigma^3+(\kappa-3)\sigma^4}{8} , \tag{S149}$$

with $\gamma_{m'} = \kappa_3(m')/\kappa_2(m')^{3/2}$ and $\kappa_{m'} - 3 = \kappa_4(m')/\kappa_2(m')^2$.

Under the negative-binomial closure, $\sigma^2_{\mathsf{nb}} = \mu_{\mathsf{nb}}\rho_{\mathsf{nb}}$, $\gamma\sigma^3 = \vartheta\sigma^2_{\mathsf{nb}}$ and $(\kappa-3)\sigma^4 = \alpha\sigma^2_{\mathsf{nb}}$, with $\vartheta = 2\rho_{\mathsf{nb}} - 1$ and $\alpha = 6\rho_{\mathsf{nb}}(\rho_{\mathsf{nb}}-1)+1$. The third cumulant in Eq. (S149) then reads $(3+\vartheta)\sigma^2_{\mathsf{nb}}/4 = (1+\rho_{\mathsf{nb}})\sigma^2_{\mathsf{nb}}/2$, while the numerator of the fourth expands and factors as

$$-2\mu_{\mathsf{nb}} + (3+6\vartheta+\alpha)\mu_{\mathsf{nb}}\rho_{\mathsf{nb}} = \mu_{\mathsf{nb}}(6\rho^3_{\mathsf{nb}} + 6\rho^2_{\mathsf{nb}} - 2\rho_{\mathsf{nb}} - 2) = 2\mu_{\mathsf{nb}}(1+\rho_{\mathsf{nb}})(3\rho^2_{\mathsf{nb}} - 1) . \tag{S150}$$

Writing the first two cumulants as the mean and variance they are, Eq. (S149) collapses to

$$E[m']_{\mathsf{nb}} = \mu_{\mathsf{nb}} \;, \qquad \mathrm{Var}[m']_{\mathsf{nb}} = \frac{\mu_{\mathsf{nb}}(1+\rho_{\mathsf{nb}})}{2} =: \sigma^2_{m',\mathsf{nb}} \;, \tag{S151}$$

$$\kappa_3(m')_{\mathsf{nb}} = \frac{\mu_{\mathsf{nb}}\rho_{\mathsf{nb}}(1+\rho_{\mathsf{nb}})}{2} \;, \qquad \kappa_4(m')_{\mathsf{nb}} = \frac{\mu_{\mathsf{nb}}(1+\rho_{\mathsf{nb}})(3\rho^2_{\mathsf{nb}}-1)}{4} \;. \tag{S152}$$

Comparing Eq. (S152) with Eq. (S151), both higher cumulants are simple multiples of the variance,

$$\kappa_3(m')_{\mathsf{nb}} = \rho_{\mathsf{nb}}\,\sigma^2_{m',\mathsf{nb}} \;, \qquad \kappa_4(m')_{\mathsf{nb}} = \frac{3\rho^2_{\mathsf{nb}}-1}{2}\,\sigma^2_{m',\mathsf{nb}} \;, \tag{S153}$$

because each carries the same factor $\mu_{\mathsf{nb}}(1+\rho_{\mathsf{nb}})/2$. Dividing Eq. (S153) by the appropriate power of $\sigma_{m',\mathsf{nb}}$, the factor $\sigma^2_{m',\mathsf{nb}}$ cancels once in each case and leaves

$$\gamma^{\mathsf{nb}}_{m'} = \frac{\kappa_3(m')_{\mathsf{nb}}}{\sigma^3_{m',\mathsf{nb}}} = \frac{\vartheta_{m'}}{\sigma_{m',\mathsf{nb}}} \;, \qquad \kappa^{\mathsf{nb}}_{m'}-3 = \frac{\kappa_4(m')_{\mathsf{nb}}}{\sigma^4_{m',\mathsf{nb}}} = \frac{\alpha_{m'}}{\sigma^2_{m',\mathsf{nb}}} \;, \qquad \vartheta_{m'} = \rho_{\mathsf{nb}} \;, \qquad \alpha_{m'} = \frac{3\rho^2_{\mathsf{nb}}-1}{2} \;. \tag{S154}$$

#### S5.2 Approximate closure ($\gamma = 0$, $\kappa = 3$)

Setting $\gamma = 0$ and $\kappa = 3$ (the Gaussian moment assumption underlying the approximate closure) in Eqs. (S139)–(S140) gives the approximate-closure pre-transposition/excision mean and variance for *any* weak-selection fitness function, not yet specialized to a particular $w(\cdot)$, as noted in the main text:

$$E[m']_{\mathsf{app}} = \mu + \beta_1\sigma^2, \tag{S155}$$

$$\mathrm{Var}[m']_{\mathsf{app}} = \frac{\mu+\sigma^2}{2} + \frac{\beta_1}{2}\sigma^2 + \frac{\beta_2}{2}\sigma^4. \tag{S156}$$

Under the Gaussian fitness function $w(n) = e^{-sn^2}$, for which $\beta_1 = -2s\mu$ and $\beta_2 = -2s$, both expressions share the common factor $(1-2s\sigma^2)$,

$$E[m']_{\mathsf{app}} = \mu(1-2s\sigma^2), \qquad \mathrm{Var}[m']_{\mathsf{app}} = \frac{\mu+\sigma^2}{2}(1-2s\sigma^2), \tag{S157}$$

which cancels in the ratio to give

$$\rho^{\mathsf{app}}_{m'} = \frac{1}{2}\Big(1+\frac{\sigma^2}{\mu}\Big) = \frac{1+\hat\rho_{\mathsf{app}}}{2}, \tag{S158}$$

independent of $s$, where $\hat\rho_{\mathsf{app}} = \sigma^2/\mu$ is the population's own dispersion ratio. Substituting the known approximate-closure equilibrium, Eq. (6) of the main text ($\hat\mu_{\mathsf{app}} = (u-v)(1-2(u-v))/[2s(2u+2v+1)]$, $\hat\sigma^2_{\mathsf{app}} = (u-v)/(2s)$), gives $\hat\rho_{\mathsf{app}} = (1+2u+2v)/(1-2u+2v)$ and hence the closed form

$$\rho^{\mathsf{app}}_{m'} = \frac{1+2v}{1-2u+2v} \approx 1+2u \qquad (v \ll u). \tag{S159}$$

#### S5.3 Negative-binomial closure under Gaussian fitness

We evaluate the pre-transposition/excision moments, Eqs. (19)–(20) of the main text, for $w(n) = \exp(-sn^2)$, so that $\beta_1 = -2s\mu_{\mathsf{nb}}$ and $\beta_2 = -2s$. Here $\mu_{\mathsf{nb}}$ and $\sigma^2_{\mathsf{nb}} = \mu_{\mathsf{nb}}\rho_{\mathsf{nb}}$ are the post-transposition/excision equilibrium moments of the negative-binomial closure, with shape terms $\vartheta = 2\rho_{\mathsf{nb}} - 1$ and $\alpha = 6\rho_{\mathsf{nb}}(\rho_{\mathsf{nb}} - 1) + 1$. The derivation uses the equilibrium mean balance obtained in Section S4, Eq. (S111), written here as

$$2s\mu_{\mathsf{nb}}\rho_{\mathsf{nb}} + s\rho_{\mathsf{nb}}(2\rho_{\mathsf{nb}} - 1) = u - v \ , \tag{S160}$$

together with the shape sum $\vartheta + \alpha = (2\rho_{\mathsf{nb}} - 1) + (6\rho_{\mathsf{nb}}^2 - 6\rho_{\mathsf{nb}} + 1) = 2\rho_{\mathsf{nb}}(3\rho_{\mathsf{nb}} - 2)$.

Substituting $\beta_1$ and $\beta_2$ into Eq. (19) gives

$$E[m']_{\mathsf{nb}} = \mu_{\mathsf{nb}} - 2s\mu_{\mathsf{nb}}^2\rho_{\mathsf{nb}} - s(2\rho_{\mathsf{nb}} - 1)\mu_{\mathsf{nb}}\rho_{\mathsf{nb}} = \mu_{\mathsf{nb}}\Big[1 - \big(2s\mu_{\mathsf{nb}}\rho_{\mathsf{nb}} + s\rho_{\mathsf{nb}}(2\rho_{\mathsf{nb}} - 1)\big)\Big] \ , \tag{S161}$$

in which the bracketed group is the left-hand side of Eq. (S160), so that

$$E[m']_{\mathsf{nb}} = \mu_{\mathsf{nb}}\left(1 - (u - v)\right) \ . \tag{S162}$$

For the variance, Eq. (20) with $1 + \vartheta = 2\rho_{\mathsf{nb}}$ and $\sigma_{\mathsf{nb}}^2 = \mu_{\mathsf{nb}}\rho_{\mathsf{nb}}$ reads

$$\begin{aligned}\mathrm{Var}[m']_{\mathsf{nb}} &= \frac{\mu_{\mathsf{nb}} + \mu_{\mathsf{nb}}\rho_{\mathsf{nb}}}{2} + \frac{1}{2}(-2s\mu_{\mathsf{nb}})(2\rho_{\mathsf{nb}})\mu_{\mathsf{nb}}\rho_{\mathsf{nb}} + \frac{1}{4}(-2s)\Big[(\vartheta + \alpha)\mu_{\mathsf{nb}}\rho_{\mathsf{nb}} + 2\mu_{\mathsf{nb}}^2\rho_{\mathsf{nb}}^2\Big] \\ &= \frac{\mu_{\mathsf{nb}} + \mu_{\mathsf{nb}}\rho_{\mathsf{nb}}}{2} - 2s\mu_{\mathsf{nb}}^2\rho_{\mathsf{nb}}^2 - \frac{s}{2}(\vartheta + \alpha)\mu_{\mathsf{nb}}\rho_{\mathsf{nb}} - s\mu_{\mathsf{nb}}^2\rho_{\mathsf{nb}}^2 \\ &= \mu_{\mathsf{nb}}\left[\frac{1 + \rho_{\mathsf{nb}}}{2} - 3s\mu_{\mathsf{nb}}\rho_{\mathsf{nb}}^2 - s\rho_{\mathsf{nb}}^2(3\rho_{\mathsf{nb}} - 2)\right] \ ,\end{aligned} \tag{S163}$$

where the last term follows from $\frac{s}{2}(\vartheta + \alpha)\mu_{\mathsf{nb}}\rho_{\mathsf{nb}} = s\mu_{\mathsf{nb}}\rho_{\mathsf{nb}}^2(3\rho_{\mathsf{nb}} - 2)$. Solving Eq. (S160) for the product, $3s\mu_{\mathsf{nb}}\rho_{\mathsf{nb}}^2 = \frac{3\rho_{\mathsf{nb}}}{2}\left[(u - v) - s\rho_{\mathsf{nb}}(2\rho_{\mathsf{nb}} - 1)\right]$, and substituting into Eq. (S163), the terms in $s\rho_{\mathsf{nb}}^2$ combine as

$$\frac{3}{2}s\rho_{\mathsf{nb}}^2(2\rho_{\mathsf{nb}} - 1) - s\rho_{\mathsf{nb}}^2(3\rho_{\mathsf{nb}} - 2) = s\rho_{\mathsf{nb}}^2\Big[3\rho_{\mathsf{nb}} - \frac{3}{2} - 3\rho_{\mathsf{nb}} + 2\Big] = \frac{s\rho_{\mathsf{nb}}^2}{2} \ , \tag{S164}$$

leaving

$$\mathrm{Var}[m']_{\mathsf{nb}} = \mu_{\mathsf{nb}}\left[\frac{1 + \rho_{\mathsf{nb}}}{2} - \frac{3\rho_{\mathsf{nb}}}{2}(u - v) + \frac{s\rho_{\mathsf{nb}}^2}{2}\right] \ . \tag{S165}$$

Equivalently, substituting $\mu_{\mathsf{nb}}\rho_{\mathsf{nb}} = \sigma_{\mathsf{nb}}^2$ and $\mu_{\mathsf{nb}}\rho_{\mathsf{nb}}^2 = \sigma_{\mathsf{nb}}^4/\mu_{\mathsf{nb}}$, the mean and variance read

$$E[m']_{\mathsf{nb}} = \mu_{\mathsf{nb}}(1 - (u - v)) \ , \qquad \mathrm{Var}[m']_{\mathsf{nb}} = \frac{1}{2}\left[\mu_{\mathsf{nb}} + \sigma_{\mathsf{nb}}^2(1 - 3(u - v)) + \frac{s\,\sigma_{\mathsf{nb}}^4}{\mu_{\mathsf{nb}}}\right] \ . \tag{S166}$$

Their ratio gives the pre-transposition/excision dispersion ratio in terms of the equilibrium mean and variance alone,

$$\rho_{m'}^{\mathsf{nb}} = \frac{\mu_{\mathsf{nb}} + \sigma_{\mathsf{nb}}^2(1 - 3(u - v)) + \dfrac{s\,\sigma_{\mathsf{nb}}^4}{\mu_{\mathsf{nb}}}}{2\mu_{\mathsf{nb}}(1 - (u - v))} \ . \tag{S167}$$

Dividing Eq. (S165) by Eq. (S162) instead, the equilibrium mean cancels:

$$\rho_{m'}^{\mathsf{nb}} = \frac{\dfrac{1 + \rho_{\mathsf{nb}}}{2} - \dfrac{3\rho_{\mathsf{nb}}}{2}(u - v) + \dfrac{s\rho_{\mathsf{nb}}^2}{2}}{1 - (u - v)} \ . \tag{S168}$$

Substituting the equilibrium dispersion ratio $\rho_{\mathsf{nb}} = 1 + 3u + v + s$ from Eq. (S115), the numerator becomes

$$1+\frac{3u+v+s}{2}-\frac{3}{2}(u-v)+\frac{s}{2}+O(u^2)=1+2v+s+O(u^2)\ , \tag{S169}$$

the terms in $\frac{3}{2}u$ cancelling, and with $1/(1-u+v)=1+u-v+O(u^2)$,

$$\rho^{\mathsf{nb}}_{m'}=1+u+v+s+O(u^2)\ . \tag{S170}$$

The same equilibrium fixes the two higher moments at this stage. Writing $\varepsilon=3u+v$ and substituting $\rho_{\mathsf{nb}}=1+\varepsilon$ from Eq. (S115), with the order-$s$ term dropped as in Section S4, the shape terms of Eq. (S154) become

$$\vartheta_{m'}=1+\varepsilon=1+3u+v\ ,\qquad \alpha_{m'}=\frac{3(1+\varepsilon)^2-1}{2}=1+3\varepsilon+\frac{3}{2}\varepsilon^2=1+3(3u+v)+\frac{3}{2}(3u+v)^2\ , \tag{S171}$$

so that, with $\sigma^2_{m',\mathsf{nb}}=\mathrm{Var}[m']_{\mathsf{nb}}$ from Eq. (S166),

$$\gamma^{\mathsf{nb}}_{m'}=\frac{1+3u+v}{\sigma_{m',\mathsf{nb}}}\ ,\qquad \kappa^{\mathsf{nb}}_{m'}-3=\frac{1+3(3u+v)+\frac{3}{2}(3u+v)^2}{\sigma^2_{m',\mathsf{nb}}}\ . \tag{S172}$$

These are the pre-transposition/excision counterparts of Eq. (S120), whose shape terms are $1+6u+2v$ and $1+6(3u+v)+6(3u+v)^2$

##### S5.4 Condition for over-, equi- and underdispersion for pre-transposition/excision stage

Using the general moments derived above, this section obtains a closed-form condition for whether $m'$ is over- or underdispersed, under the negative-binomial closure and under the approximate closure. Throughout, $\mu$, $\sigma^2$, $\beta_1$, $\beta_2$ denote the equilibrium mean, variance, and log-fitness derivatives of the standing, post-transposition/excision population, and $\rho=\sigma^2/\mu$ is the corresponding post-transposition/excision dispersion ratio used elsewhere in the main text.

###### S5.4.1 Negative-binomial closure

Substituting the negative-binomial shape relations $\vartheta=(2-p)/p$ and $\alpha=6(1-p)+p^2$, where $p=\mu/\sigma^2$, into the general pre-transposition/excision moments, Eqs. (S139)–(S140), gives the negative-binomial pre-transposition/excision mean and variance:

$$E[m']_{\mathsf{nb}}=\mu+\beta_1\sigma^2+\frac{1}{2}\vartheta\,\beta_2\sigma^2, \tag{S173}$$

$$\mathrm{Var}[m']_{\mathsf{nb}}=\frac{\mu+\sigma^2}{2}+\frac{1}{2}\beta_1(1+\vartheta)\sigma^2+\frac{1}{4}\beta_2\left[(\vartheta+\alpha)\sigma^2+2\sigma^4\right], \tag{S174}$$

where $\vartheta$ and $\alpha$ are the negative-binomial shape terms defined in Eqs. (10)–(11) of the main text.

Define

$$\Delta:=\mathrm{Var}[m']_{\mathsf{nb}}-E[m']_{\mathsf{nb}}. \tag{S175}$$

Subtracting Eq. (S173) from Eq. (S174) gives

$$\Delta=\frac{\sigma^2-\mu}{2}+\frac{\beta_1}{2}(\vartheta-1)\sigma^2+\frac{\beta_2}{4}(\alpha-\vartheta)\sigma^2+\frac{\beta_2}{2}\sigma^4. \tag{S176}$$

Writing $p_{\mathsf{nb}} = \mu/\sigma^2$ and $\rho = \sigma^2/\mu = 1/p_{\mathsf{nb}}$ for the population's own (post-transposition/excision) dispersion ratio, the shape terms of Eqs. (10)–(11) of the main text become $\vartheta = 2\rho - 1$ and $\alpha = 6\rho^2 - 6\rho + 1$, so that

$$\vartheta - 1 = 2(\rho - 1), \qquad \alpha - \vartheta = 2(3\rho - 1)(\rho - 1). \tag{S177}$$

Substituting these, together with $\sigma^2 = \mu\rho$, into Eq. (S176) and collecting every term over the common factor $\mu/2$ gives

$$\Delta = \mathrm{Var}[m']_{\mathsf{nb}} - E[m']_{\mathsf{nb}} = \frac{\mu}{2}\Big\{(\rho - 1)(1 + 2\beta_1\rho) + \beta_2\,\rho\big[3\rho^2 + (\mu - 4)\rho + 1\big]\Big\}, \tag{S178}$$

evaluated at the negative-binomial closure's equilibrium $(\hat{\mu}_{\mathsf{nb}}, \hat{\sigma}^2_{\mathsf{nb}})$, with $\rho = \hat{\sigma}^2_{\mathsf{nb}}/\hat{\mu}_{\mathsf{nb}}$. Because $E[m']_{\mathsf{nb}} > 0$ at any admissible equilibrium, the sign of $\rho^{\mathsf{nb}}_{m'} - 1$ is exactly the sign of $\Delta$, so the pre-transposition/excision dispersion condition follows without ever computing $\rho^{\mathsf{nb}}_{m'}$ itself:

$$\rho^{\mathsf{nb}}_{m'} \gtrless 1 \quad\Longleftrightarrow\quad (\rho - 1)(1 + 2\beta_1\rho) + \beta_2\,\rho\big[3\rho^2 + (\mu - 4)\rho + 1\big] \gtrless 0. \tag{S179}$$

A valid negative-binomial shape requires $p_{\mathsf{nb}} = \mu/\sigma^2 \in (0, 1)$, i.e. $\rho \geq 1$ at any admissible closure equilibrium. Hence the first term in Eq. (S179), $(\rho - 1)(1 + 2\beta_1\rho)$, is never negative for the mild selection strengths considered here ($\beta_1\rho \gtrsim -\frac{1}{2}$).

The quadratic factor multiplying $\beta_2$, $3\rho^2 + (\mu - 4)\rho + 1$, is positive at $\rho = 1$, where it equals $3 + (\mu - 4) + 1 = \mu > 0$, and stays positive for every admissible $\rho$: its vertex lies at $\rho = (4 - \mu)/6 < 1$, so the factor is increasing on $\rho \geq 1$ and is therefore bounded below by $\mu$ there. Close to Poisson-like dispersion ($\rho \approx 1$), the sign of $\beta_2$, the curvature of $\ln w(\cdot)$, therefore decides the outcome: a concave $\ln w$ ($\beta_2 < 0$) pulls $\Delta$ downward, and pre-transposition/excision *underdispersion* ($\rho^{\mathsf{nb}}_{m'} < 1$) occurs once this curvature term outweighs the (small, since $\rho - 1 \ll 1$) first term, and vice versa.

Underdispersion also deepens as $u$ increases, because $\rho$ enters the two terms of Eq. (S179) at different orders: the restoring term is linear in $\rho$ (via $\rho - 1$), while the curvature term carries a further quadratic in $\rho$ on top of its own leading factor, effectively cubic overall. Raising $u$ therefore inflates the curvature term much faster than the restoring one. A second, independent channel reinforces this for any fitness function with $\gamma > 2$ (i.e. $\beta_2 \propto \mu^{\gamma-2}$): $\beta_2$ itself grows in magnitude as $\mu$ rises with $u$. Once the curvature term overtakes the restoring term, further increases in $u$ compound the underdispersion rather than merely sustain it.

##### S5.4.2 Approximate closure

Recall the general approximate-closure pre-transposition/excision mean and variance, Eqs. (S155)–(S156) above. Subtracting Eq. (S155) from Eq. (S156) and substituting $\sigma^2 = \mu\rho$, exactly as in the negative-binomial case above, gives

$$\Delta_{\mathsf{app}} = \mathrm{Var}[m']_{\mathsf{app}} - E[m']_{\mathsf{app}} = \frac{\mu}{2}\Big\{(\rho - 1) - \beta_1\rho + \beta_2\mu\rho^2\Big\}, \tag{S180}$$

so that

$$\rho^{\mathsf{app}}_{m'} \gtrless 1 \quad\Longleftrightarrow\quad (\rho - 1) - \beta_1\rho + \beta_2\mu\rho^2 \gtrless 0, \tag{S181}$$

which holds for any weak-selection fitness function; this is the approximate-closure analogue of Eq. (S179).

As before, $(\rho - 1) \geq 0$, but the sign of the remaining term, $\rho(\beta_2\mu\rho - \beta_1)$, is not fixed in general: it pits a linear-selection contribution ($-\beta_1\rho$, typically positive since $\beta_1 < 0$) against a curvature contribution ($\beta_2\mu\rho^2$, typically negative since $\beta_2 < 0$). The approximate closure therefore admits pre-transposition/excision underdispersion under exactly the same qualitative condition as the negative-binomial closure: whenever the curvature term outweighs the (small) $(\rho - 1)$ and linear-selection terms and vice versa. This general condition should be distinguished from the Gaussian fitness function $w(n) = e^{-sn^2}$, for which $\beta_2\mu = \beta_1$ identically; that additional relation is a special case of Eqs. (S180)–(S181) above, not a property of the approximate closure in general.

#### S5.5 Equidispersion

Both closures admit pre-transposition/excision underdispersion once the curvature term in Eqs. (S179) and (S181) dominates the restoring (and, for the approximate closure, linear-selection) term. The exact boundary separating over- from underdispersion, equidispersion ($\rho_{m'} = 1$), is where these conditions hold with equality. For the negative-binomial closure, this fixes $\beta_2$ as

$$\beta_2 = -\frac{(\rho - 1)(1 + 2\beta_1\rho)}{\rho[3\rho^2 + (\mu - 4)\rho + 1]}, \tag{S182}$$

and for the approximate closure as

$$\beta_2 = \frac{\beta_1\rho - (\rho - 1)}{\mu\rho^2}. \tag{S183}$$

In both cases, curvature weaker than this boundary value ($\beta_2$ less negative) leaves $m'$ overdispersed, while curvature beyond it ($\beta_2$ more negative) drives underdispersion.

#### DPGP3 per-strain TE burden distributions (histograms)

This section gives empirical per-strain TE burden distributions for the DPGP3 Zambian *D. melanogaster* panel analysed in the main text. Supplementary Figure S1 shows histograms for every filtered family that passes the inclusion criteria; Supplementary Table S1 lists the corresponding sample sizes, mean, variance, dispersion ratio $\rho$, skewness, and excess kurtosis. The main text illustrates three representative families in Fig. 6; the appendix of the main document gives a compact summary table for those families (Section C).

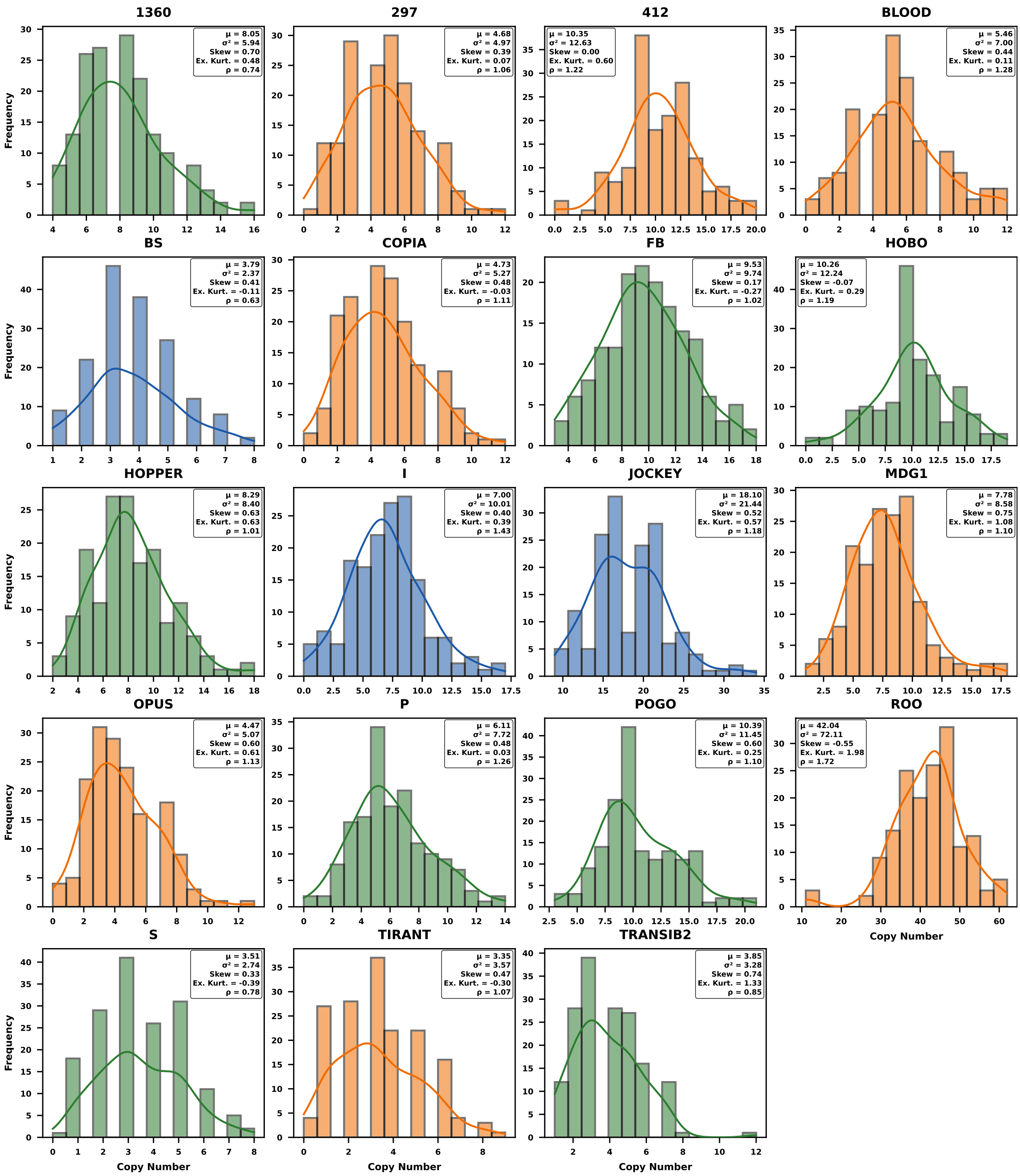


Figure S1: **Distribution of per-strain TE burdens across filtered families (DPGP3).** Shown are per-strain burden histograms for all $19$ TE families that pass the filtering criteria in the main text; Table S1 summarises their mean, variance, dispersion ratio $\rho$, skewness, and excess kurtosis.

| TE Family | Class | $n_{\text{present}}$ | Mean | Variance | Var./Mean ($\rho$) | Skewness | Excess kurtosis |
|---|---|---|---|---|---|---|---|
| **Overdispersed families** | | | | | | | |
| ROO | LTR | 6895.0 | 42.042683 | 72.114731 | 1.715274 | -0.547687 | 1.979038 |
| I | non-LTR | 1148.0 | 7.000000 | 10.012270 | 1.430324 | 0.401096 | 0.394768 |
| BLOOD | LTR | 896.0 | 5.463415 | 6.998653 | 1.281004 | 0.444032 | 0.108588 |
| P | DNA | 1002.0 | 6.109756 | 7.717941 | 1.263216 | 0.476073 | 0.028203 |
| 412 | LTR | 1697.0 | 10.347561 | 12.633061 | 1.220873 | 0.002909 | 0.602917 |
| HOBO | DNA | 1683.0 | 10.262195 | 12.243715 | 1.193089 | -0.066071 | 0.293218 |
| JOCKEY | non-LTR | 2968.0 | 18.097561 | 21.438276 | 1.184595 | 0.517010 | 0.567721 |
| OPUS | LTR | 733.0 | 4.469512 | 5.072684 | 1.134953 | 0.596543 | 0.610915 |
| COPIA | LTR | 775.0 | 4.725610 | 5.267806 | 1.114736 | 0.477319 | -0.025089 |
| MDG1 | LTR | 1276.0 | 7.780488 | 8.577286 | 1.102410 | 0.745779 | 1.081381 |
| POGO | DNA | 1704.0 | 10.390244 | 11.454137 | 1.102394 | 0.599592 | 0.251547 |
| TIRANT | LTR | 549.0 | 3.347561 | 3.565577 | 1.065127 | 0.472408 | -0.295699 |
| 297 | LTR | 768.0 | 4.682927 | 4.966332 | 1.060519 | 0.392393 | 0.071436 |
| FB | DNA | 1563.0 | 9.530488 | 9.735261 | 1.021486 | 0.167479 | -0.269287 |
| HOPPER | DNA | 1360.0 | 8.292683 | 8.404609 | 1.013497 | 0.628323 | 0.631285 |
| **Underdispersed families** | | | | | | | |
| BS | non-LTR | 622.0 | 3.792683 | 2.373934 | 0.625925 | 0.414527 | -0.110042 |
| 1360 | DNA | 1320.0 | 8.048780 | 5.936256 | 0.737535 | 0.697525 | 0.481536 |
| S | DNA | 575.0 | 3.506098 | 2.742294 | 0.782150 | 0.329700 | -0.385421 |
| TRANSIB2 | DNA | 632.0 | 3.853659 | 3.279066 | 0.850897 | 0.735486 | 1.329797 |

Table S1: Summary statistics for filtered TE families in the DPGP3 Zambian *D. melanogaster* dataset of Lee (2022). We report the number of presence calls summed across sites and strains ($n_{\text{present}}$), the mean and variance of per-strain TE burdens, the dispersion ratio $\rho$, and higher moments for each family.

#### Equilibrium moments: simulation versus theory

Table S2 summarises equilibrium mean, variance, skewness, and excess kurtosis for the stochastic Moran simulations and theoretical closures in Figs. 1 and 2 of the main text.

| | Mean | Var. | Skew. | Ex. Kurt. |
|---|---|---|---|---|
| Stochastic sim. | 92.99 | 100.29 | 0.12 | −0.02 |
| Theory (neg. bin.) | 91.66 | 99.37 | 0.12 | 0.02 |
| Theory (approx.) | 90.23 | 100.00 | — | — |
| Ch. & Ch. (1983) | 100.00 | 100.00 | 0.10 | 0.01 |

Table S2: **Comparison of stochastic simulation and theoretical values.** The table shows the mean, variance, skewness and kurtosis values corresponding to Fig. 1a, Fig. 1b, and Fig. 2 of the main text. The Charlesworth and Charlesworth (1983) values are computed from Eq. 8. The approximate closure values are computed using Eq. 6, and the parametric (negative binomial) closure values are computed from simulation outputs of Eq. 12. The stochastic model was simulated for 30,000 generations, and a single equilibrium value was obtained by averaging samples taken every 100 generations after a burn-in period of $2{,}000$ generations. The skewness and excess kurtosis for Charlesworth and Charlesworth (1983) are computed using $1/\sqrt{\mu_c}$ and $1/\mu_c$, respectively, from the Poisson distribution assumed by their model. The parametric closure skewness and excess kurtosis values are computed using Eqs. 10 and 11, while the approximate closure values are not reported as the higher moments are assumed zero.

#### Equilibrium moments across transposition rates

Tables S3–S4 report equilibrium mean, variance, dispersion ratio $\rho = \mathrm{Var}/\mathrm{Mean}$, skewness, and excess kurtosis at the pre-transposition/excision stage (i.e., $\rho_{m'}$, immediately after recombination but before transposition and excision act), and Tables S5–S6 report the same quantities at the post-transposition/excision stage (i.e., after a complete birth event), for a sequence of transposition rates $u$, in the same setting as the main-text comparison of theoretical predictions and stochastic simulations across transposition rates: diploid population size $N = 10^4$, selection coefficient $s = 1/N$, free (binomial) recombination, excision rate $v = 10^{-4}$, exponential fitness $w(n) = \exp(-sn^2)$, and $u$ ranging from $5 \times 10^{-4}$ to $0.04$. Together, these four tables list the numerical moment values for Fig. 3 of the main text. Dashes denote skewness and excess kurtosis not reported for the approximate closure or for Roze (2023).

| **Transposition rate** $u$ | $\mathbf{5 \times 10^{-4}}$ | $\mathbf{10^{-3}}$ | $\mathbf{1.5 \times 10^{-3}}$ | $\mathbf{2.2 \times 10^{-3}}$ | $\mathbf{3 \times 10^{-3}}$ | $\mathbf{4 \times 10^{-3}}$ | $\mathbf{5 \times 10^{-3}}$ | $\mathbf{6 \times 10^{-3}}$ |
|---|---|---|---|---|---|---|---|---|
| **Mean** ($\mu$) | | | | | | | | |
| Simulation | 1.81 | 3.70 | 6.51 | 9.99 | 13.75 | 18.68 | 23.47 | 28.43 |
| NB closure | 1.49 | 3.98 | 6.45 | 9.90 | 13.82 | 18.68 | 23.50 | 28.29 |
| Approx. closure | 2.00 | 4.48 | 6.96 | 10.41 | 14.33 | 19.19 | 24.01 | 28.80 |
| Roze (2023) | 2.00 | 4.50 | 7.00 | 10.50 | 14.50 | 19.50 | 24.50 | 29.50 |
| **Variance** ($\sigma^2$) | | | | | | | | |
| Simulation | 1.81 | 3.70 | 6.52 | 10.02 | 13.78 | 18.74 | 23.59 | 28.58 |
| NB closure | 1.50 | 3.98 | 6.46 | 9.93 | 13.86 | 18.76 | 23.62 | 28.46 |
| Approx. closure | 2.00 | 4.49 | 6.98 | 10.45 | 14.41 | 19.35 | 24.26 | 29.15 |
| Roze (2023) | 2.00 | 4.50 | 7.01 | 10.52 | 14.54 | 19.58 | 24.62 | 29.68 |
| **Dispersion ratio** ($\rho$) | | | | | | | | |
| Simulation | 1.0006 | 1.0004 | 1.0018 | 1.0025 | 1.0022 | 1.0030 | 1.0048 | 1.0053 |
| NB closure | 1.0007 | 1.0012 | 1.0017 | 1.0024 | 1.0032 | 1.0041 | 1.0051 | 1.0061 |
| Approx. closure | 1.0010 | 1.0020 | 1.0030 | 1.0044 | 1.0060 | 1.0081 | 1.0101 | 1.0121 |
| Roze (2023) | 1.0006 | 1.0011 | 1.0016 | 1.0023 | 1.0031 | 1.0041 | 1.0051 | 1.0061 |
| **Skewness** ($\gamma$) | | | | | | | | |
| Simulation | 0.75 | 0.52 | 0.39 | 0.32 | 0.27 | 0.23 | 0.21 | 0.19 |
| NB closure | 0.82 | 0.50 | 0.40 | 0.32 | 0.27 | 0.23 | 0.21 | 0.19 |
| **Excess kurtosis** ($\kappa - 3$) | | | | | | | | |
| Simulation | 0.56 | 0.27 | 0.16 | 0.10 | 0.07 | 0.06 | 0.04 | 0.03 |
| NB closure | 0.67 | 0.25 | 0.16 | 0.10 | 0.07 | 0.06 | 0.04 | 0.04 |

Table S3: **Equilibrium moments across transposition rates, pre-transposition/excision (Part 1).** Each column corresponds to a different transposition rate $u$. Values are measured immediately after recombination, before transposition and excision act (see the pre-transposition/excision mean and variance section of the main text). The stochastic model was simulated for 30,000 generations with averaging after a burn-in period of 2,000 generations.

| **Transposition rate** $u$ | $\mathbf{8 \times 10^{-3}}$ | $\mathbf{10^{-2}}$ | $\mathbf{1.2 \times 10^{-2}}$ | $\mathbf{1.6 \times 10^{-2}}$ | $\mathbf{2 \times 10^{-2}}$ | $\mathbf{2.6 \times 10^{-2}}$ | $\mathbf{3.2 \times 10^{-2}}$ | $\mathbf{4 \times 10^{-2}}$ |
|---|---|---|---|---|---|---|---|---|
| **Mean** ($\mu$) | | | | | | | | |
| Simulation | 37.80 | 47.40 | 56.57 | 74.87 | 92.68 | 118.58 | 143.71 | 175.79 |
| NB closure | 37.74 | 47.05 | 56.21 | 74.10 | 91.43 | 116.43 | 140.28 | 170.39 |
| Approx. closure | 38.26 | 47.56 | 56.71 | 74.57 | 91.85 | 116.70 | 140.32 | 169.95 |
| Roze (2023) | 39.50 | 49.50 | 59.50 | 79.50 | 99.50 | 129.50 | 159.50 | 199.50 |
| **Variance** ($\sigma^2$) | | | | | | | | |
| Simulation | 38.08 | 47.85 | 57.20 | 75.90 | 94.34 | 121.27 | 147.74 | 181.63 |
| NB closure | 38.04 | 47.51 | 56.87 | 75.23 | 93.15 | 119.20 | 144.29 | 176.25 |
| Approx. closure | 38.88 | 48.53 | 58.11 | 77.04 | 95.67 | 123.10 | 149.91 | 184.72 |
| Roze (2023) | 39.82 | 50.00 | 60.22 | 80.78 | 101.50 | 132.88 | 164.62 | 207.50 |
| **Dispersion ratio** ($\rho$) | | | | | | | | |
| Simulation | 1.0075 | 1.0096 | 1.0111 | 1.0138 | 1.0180 | 1.0227 | 1.0281 | 1.0332 |
| NB closure | 1.0080 | 1.0099 | 1.0117 | 1.0153 | 1.0188 | 1.0238 | 1.0285 | 1.0344 |
| Approx. closure | 1.0163 | 1.0204 | 1.0246 | 1.0331 | 1.0417 | 1.0548 | 1.0684 | 1.0869 |
| Roze (2023) | 1.0081 | 1.0101 | 1.0121 | 1.0161 | 1.0201 | 1.0261 | 1.0321 | 1.0401 |
| **Skewness** ($\gamma$) | | | | | | | | |
| Simulation | 0.17 | 0.15 | 0.14 | 0.12 | 0.11 | 0.09 | 0.09 | 0.08 |
| NB closure | 0.17 | 0.15 | 0.14 | 0.12 | 0.11 | 0.10 | 0.09 | 0.08 |
| **Excess kurtosis** ($\kappa - 3$) | | | | | | | | |
| Simulation | 0.03 | 0.02 | 0.02 | 0.01 | 0.01 | 0.01 | 0.01 | 0.00 |
| NB closure | 0.03 | 0.02 | 0.02 | 0.02 | 0.01 | 0.01 | 0.01 | 0.01 |

Table S4: **Equilibrium moments across transposition rates, pre-transposition/excision (Part 2).** Continuation of Table S3 for higher transposition rates.

| **Transposition rate** $u$ | $\mathbf{5 \times 10^{-4}}$ | $\mathbf{10^{-3}}$ | $\mathbf{1.5 \times 10^{-3}}$ | $\mathbf{2.2 \times 10^{-3}}$ | $\mathbf{3 \times 10^{-3}}$ | $\mathbf{4 \times 10^{-3}}$ | $\mathbf{5 \times 10^{-3}}$ | $\mathbf{6 \times 10^{-3}}$ |
|---|---|---|---|---|---|---|---|---|
| **Mean** ($\mu$) | | | | | | | | |
| Simulation | 1.81 | 3.70 | 6.52 | 10.01 | 13.79 | 18.76 | 23.59 | 28.60 |
| NB closure | 1.49 | 3.98 | 6.46 | 9.92 | 13.86 | 18.75 | 23.62 | 28.45 |
| Approx. closure | 2.00 | 4.48 | 6.96 | 10.41 | 14.33 | 19.19 | 24.01 | 28.80 |
| Roze (2023) | 2.00 | 4.50 | 7.00 | 10.50 | 14.50 | 19.50 | 24.50 | 29.50 |
| **Variance** ($\sigma^2$) | | | | | | | | |
| Simulation | 1.81 | 3.71 | 6.55 | 10.08 | 13.90 | 18.96 | 23.94 | 29.09 |
| NB closure | 1.50 | 4.00 | 6.49 | 9.99 | 13.99 | 18.98 | 23.98 | 28.97 |
| Approx. closure | 2.00 | 4.50 | 7.00 | 10.50 | 14.50 | 19.50 | 24.50 | 29.50 |
| Roze (2023) | 2.00 | 4.51 | 7.03 | 10.57 | 14.63 | 19.74 | 24.87 | 30.03 |
| **Dispersion ratio** ($\rho$) | | | | | | | | |
| Simulation | 1.0016 | 1.0024 | 1.0047 | 1.0068 | 1.0082 | 1.0110 | 1.0148 | 1.0171 |
| NB closure | 1.0017 | 1.0032 | 1.0047 | 1.0068 | 1.0092 | 1.0122 | 1.0152 | 1.0182 |
| Approx. closure | 1.0020 | 1.0040 | 1.0060 | 1.0088 | 1.0121 | 1.0161 | 1.0202 | 1.0243 |
| Roze (2023) | 1.0016 | 1.0031 | 1.0046 | 1.0067 | 1.0091 | 1.0121 | 1.0151 | 1.0181 |
| **Skewness** ($\gamma$) | | | | | | | | |
| Simulation | 0.75 | 0.53 | 0.40 | 0.32 | 0.27 | 0.24 | 0.21 | 0.19 |
| NB closure | 0.82 | 0.50 | 0.40 | 0.32 | 0.27 | 0.24 | 0.21 | 0.19 |
| **Excess kurtosis** ($\kappa - 3$) | | | | | | | | |
| Simulation | 0.56 | 0.28 | 0.16 | 0.10 | 0.07 | 0.06 | 0.05 | 0.04 |
| NB closure | 0.67 | 0.26 | 0.16 | 0.10 | 0.08 | 0.06 | 0.05 | 0.04 |

Table S5: **Equilibrium moments across transposition rates, post-transposition/excision (Part 1).** Each column corresponds to a different transposition rate $u$. Values are measured after a complete birth event, i.e., after recombination, transposition, and excision have all acted, and correspond directly to Fig. 3 and Table 1 of the main text. The stochastic model was simulated for 30,000 generations with averaging after a burn-in period of 2,000 generations.

| Transposition rate $u$ | $8 \times 10^{-3}$ | $10^{-2}$ | $1.2 \times 10^{-2}$ | $1.6 \times 10^{-2}$ | $2 \times 10^{-2}$ | $2.6 \times 10^{-2}$ | $3.2 \times 10^{-2}$ | $4 \times 10^{-2}$ |
|---|---|---|---|---|---|---|---|---|
| **Mean ($\mu$)** | | | | | | | | |
| Simulation | 38.09 | 47.87 | 57.24 | 76.06 | 94.52 | 121.65 | 148.29 | 182.81 |
| NB closure | 38.04 | 47.52 | 56.88 | 75.30 | 93.29 | 119.53 | 144.91 | 177.47 |
| Approx. closure | 38.26 | 47.56 | 56.71 | 74.57 | 91.85 | 116.70 | 140.32 | 169.95 |
| Roze (2023) | 39.50 | 49.50 | 59.50 | 79.50 | 99.50 | 129.50 | 159.50 | 199.50 |
| **Variance ($\sigma^2$)** | | | | | | | | |
| Simulation | 38.99 | 49.27 | 59.25 | 79.53 | 99.98 | 130.66 | 161.80 | 203.13 |
| NB closure | 38.96 | 48.95 | 58.94 | 78.93 | 98.91 | 128.88 | 158.85 | 198.81 |
| Approx. closure | 39.50 | 49.50 | 59.50 | 79.50 | 99.50 | 129.50 | 159.50 | 199.50 |
| Roze (2023) | 40.45 | 50.99 | 61.65 | 83.32 | 105.48 | 139.61 | 174.83 | 223.46 |
| **Dispersion ratio ($\rho$)** | | | | | | | | |
| Simulation | 1.0235 | 1.0294 | 1.0350 | 1.0456 | 1.0578 | 1.0740 | 1.0911 | 1.1112 |
| NB closure | 1.0242 | 1.0302 | 1.0362 | 1.0482 | 1.0602 | 1.0782 | 1.0962 | 1.1202 |
| Approx. closure | 1.0325 | 1.0408 | 1.0492 | 1.0661 | 1.0833 | 1.1097 | 1.1367 | 1.1739 |
| Roze (2023) | 1.0241 | 1.0301 | 1.0361 | 1.0481 | 1.0601 | 1.0781 | 1.0961 | 1.1201 |
| **Skewness ($\gamma$)** | | | | | | | | |
| Simulation | 0.17 | 0.15 | 0.14 | 0.12 | 0.11 | 0.10 | 0.09 | 0.08 |
| NB closure | 0.17 | 0.15 | 0.14 | 0.12 | 0.11 | 0.10 | 0.09 | 0.09 |
| **Excess kurtosis ($\kappa - 3$)** | | | | | | | | |
| Simulation | 0.03 | 0.03 | 0.02 | 0.02 | 0.01 | 0.01 | 0.01 | 0.00 |
| NB closure | 0.03 | 0.02 | 0.02 | 0.02 | 0.01 | 0.01 | 0.01 | 0.01 |

Table S6: **Equilibrium moments across transposition rates, post-transposition/excision (Part 2).** Continuation of Table S5 for higher transposition rates.

#### Equilibrium moments across map length $R$

Tables S7–S8 report equilibrium mean, variance, dispersion ratio $\rho = \mathrm{Var}/\mathrm{Mean}$, skewness, and excess kurtosis at a sequence of chromosome map lengths $R$ (expected number of crossovers per meiosis, Morgans). All values are recorded at the *pre-transposition/excision* stage, that is, immediately after recombination and before the current generation's transposition and excision. The simulation rows use the offspring copy number $m'$ at that point in the life cycle; the negative-binomial closure uses Eqs. (23) and (24) of the main text, and the approximate closure uses Eq. (S159) of the Supplementary Methods. For Roze (2023) we report two versions of the dispersion ratio. The first is the ratio of variance to mean, with the mean given by their Eq. (11) and the variance combining their Poisson baseline with linkage terms as in their Eqs. (5) and (13). The second is computed directly from their Eq. (14), which predicts $\rho$ without passing through the two moments. Dashes denote skewness and excess kurtosis not reported for the approximate closure or for Roze (2023).

| **Map length** $R$ | $\mathbf{10^{-5}}$ | $\mathbf{10^{-4}}$ | $\mathbf{10^{-3}}$ | $\mathbf{10^{-2}}$ | $\mathbf{10^{-1}}$ |
|---|---|---|---|---|---|
| | | | **Mean** ($\mu$) | | |
| Simulation | 5.66 | 5.58 | 5.64 | 6.23 | 7.78 |
| NB closure | 8.99 | 8.99 | 8.99 | 8.99 | 8.99 |
| Approx. closure | 9.42 | 9.42 | 9.42 | 9.42 | 9.42 |
| Roze (2023) Eq. 11 | 9.00 | 9.00 | 9.00 | 9.00 | 9.00 |
| | | | **Variance** ($\sigma^2$) | | |
| Simulation | 8.40 | 8.29 | 8.33 | 8.54 | 8.91 |
| NB closure | 9.08 | 9.08 | 9.08 | 9.08 | 9.08 |
| Approx. closure | 9.61 | 9.61 | 9.61 | 9.61 | 9.61 |
| Roze (2023) Eq. 5+13 | 11.47 | 11.47 | 11.43 | 11.14 | 10.14 |
| | | | **Dispersion ratio** ($\rho$) | | |
| Simulation | 1.4834 | 1.4872 | 1.4790 | 1.3723 | 1.1454 |
| NB closure | 1.0103 | 1.0103 | 1.0103 | 1.0103 | 1.0103 |
| Approx. closure | 1.0204 | 1.0204 | 1.0204 | 1.0204 | 1.0204 |
| Roze (2023) Eq. 11, 5+13 | 1.2750 | 1.2745 | 1.2705 | 1.2380 | 1.1265 |
| Roze (2023) Eq. (14) | 1.5498 | 1.5482 | 1.5324 | 1.4197 | 1.1643 |
| | | | **Skewness** ($\gamma$) | | |
| Simulation | 0.62 | 0.63 | 0.61 | 0.55 | 0.42 |
| NB closure | 0.34 | 0.34 | 0.34 | 0.34 | 0.34 |
| | | | **Excess kurtosis** ($\kappa - 3$) | | |
| Simulation | 0.45 | 0.44 | 0.43 | 0.34 | 0.20 |
| NB closure | 0.12 | 0.12 | 0.12 | 0.12 | 0.12 |

Table S7: **Equilibrium moments across map length $R$, pre-transposition/excision (Part 1: low recombination).** Each column corresponds to a different chromosome map length $R$ (Morgans). Moments are measured immediately after recombination, before transposition and excision.

| **Map length** $R$ | **1** | **10** | **100** | **1000** |
|---|---|---|---|---|
| | | **Mean** ($\mu$) | | |
| Simulation | 8.80 | 9.00 | 9.05 | — |
| NB closure | 8.99 | 8.99 | 8.99 | 8.99 |
| Approx. closure | 9.42 | 9.42 | 9.42 | 9.42 |
| Roze (2023) Eq. 11 | 9.00 | 9.00 | 9.00 | 9.00 |
| | | **Variance** ($\sigma^2$) | | |
| Simulation | 9.09 | 9.09 | 9.13 | — |
| NB closure | 9.08 | 9.08 | 9.08 | 9.08 |
| Approx. closure | 9.61 | 9.61 | 9.61 | 9.61 |
| Roze (2023) Eq. 5+13 | 9.30 | 9.13 | 9.10 | 9.10 |
| | | **Dispersion ratio** ($\rho$) | | |
| Simulation | 1.0331 | 1.0101 | 1.0082 | — |
| NB closure | 1.0103 | 1.0103 | 1.0103 | 1.0103 |
| Approx. closure | 1.0204 | 1.0204 | 1.0204 | 1.0204 |
| Roze (2023) Eq. 11, 5+13 | 1.0331 | 1.0139 | 1.0109 | 1.0106 |
| Roze (2023) Eq. (14) | 1.0352 | 1.0143 | 1.0111 | 1.0108 |
| | | **Skewness** ($\gamma$) | | |
| Simulation | 0.35 | 0.34 | 0.34 | — |
| NB closure | 0.34 | 0.34 | 0.34 | 0.34 |
| | | **Excess kurtosis** ($\kappa - 3$) | | |
| Simulation | 0.13 | 0.13 | 0.12 | — |
| NB closure | 0.12 | 0.12 | 0.12 | 0.12 |

Table S8: **Equilibrium moments across map length $R$, pre-transposition/excision (Part 2: high recombination).** Continuation of Table S7 for higher map lengths.

#### Equilibrium moments across selection strengths

Table S9 reports equilibrium mean, variance, dispersion ratio $\rho = \mathrm{Var}/\mathrm{Mean}$, skewness, and excess kurtosis for a sequence of selection coefficients $s$, in the same setting as the main-text comparison of theoretical predictions and stochastic simulations across selection strengths (Fig. 4): diploid population size $N = 10^4$, transposition rate $u = 0.01$, excision rate $v = 10^{-4}$, free (binomial) recombination, and exponential fitness $w(n) = \exp(-sn^2)$. For Roze (2023), the mean is their Eq. (11), $\bar{n} = (u - v)/\beta$ with $\beta = 2s$, and the variance is their Eq. (5) evaluated at the post-transposition/excision stage, $\sigma^2 = \bar{n}(1 + 4u) - \beta\bar{n}^2$, with $\rho$ taken as the ratio of the two.

| **Selection coefficient** $s$ | $\mathbf{3 \times 10^{-5}}$ | $\mathbf{10^{-4}}$ | $\mathbf{3 \times 10^{-4}}$ | $\mathbf{5 \times 10^{-4}}$ | $\mathbf{10^{-3}}$ | $\mathbf{2 \times 10^{-3}}$ |
|---|---|---|---|---|---|---|
| **Mean** ($\mu$) | | | | | | |
| Simulation | 160.37 | 47.80 | 15.56 | 9.13 | 4.29 | 1.88 |
| NB closure | 159.64 | 47.52 | 15.48 | 9.08 | 4.27 | 1.87 |
| Approx. closure | 158.53 | 47.56 | 15.85 | 9.51 | 4.76 | 2.38 |
| Roze (2023) | 165.00 | 49.50 | 16.50 | 9.90 | 4.95 | 2.48 |
| **Variance** ($\sigma^2$) | | | | | | |
| Simulation | 165.13 | 49.20 | 16.03 | 9.41 | 4.43 | 1.95 |
| NB closure | 164.45 | 48.95 | 15.95 | 9.35 | 4.40 | 1.93 |
| Approx. closure | 165.00 | 49.50 | 16.50 | 9.90 | 4.95 | 2.48 |
| Roze (2023) | 169.97 | 50.99 | 17.00 | 10.20 | 5.10 | 2.55 |
| **Dispersion ratio** ($\rho$) | | | | | | |
| Simulation | 1.0296 | 1.0293 | 1.0304 | 1.0302 | 1.0320 | 1.0331 |
| NB closure | 1.0301 | 1.0302 | 1.0304 | 1.0306 | 1.0311 | 1.0322 |
| Approx. closure | 1.0408 | 1.0408 | 1.0408 | 1.0408 | 1.0408 | 1.0408 |
| Roze (2023) | 1.0301 | 1.0301 | 1.0301 | 1.0301 | 1.0301 | 1.0301 |
| **Skewness** ($\gamma$) | | | | | | |
| Simulation | 0.08 | 0.15 | 0.26 | 0.35 | 0.51 | 0.76 |
| NB closure | 0.08 | 0.15 | 0.27 | 0.35 | 0.51 | 0.77 |
| **Excess kurtosis** ($\kappa - 3$) | | | | | | |
| Simulation | 0.00 | 0.02 | 0.07 | 0.13 | 0.28 | 0.61 |
| NB closure | 0.01 | 0.02 | 0.07 | 0.13 | 0.27 | 0.62 |

Table S9: **Equilibrium moments across selection strengths.** Each column corresponds to a different selection coefficient $s$. The table lists the numerical moment values for Fig. 4 of the main text. The stochastic model was simulated for 50,000 generations, and a single equilibrium value was obtained by averaging samples taken every 100 generations after a burn-in period of 9,000 generations. Theoretical predictions are computed as in Fig. 3; see the caption of Table 1 for details.

#### References

Charlesworth, B. and Charlesworth, D. The population dynamics of transposable elements. *Genetics Research*, 42(1):1–27, 1983. doi: 10.1017/S0016672300021455.

Lee, Y. C. G. Synergistic epistasis of the deleterious effects of transposable elements. *Genetics*, 220 (2):iyab211, 2022. doi: 10.1093/genetics/iyab211.

Roze, D. Causes and consequences of linkage disequilibrium among transposable elements within eukaryotic genomes. *Genetics*, 224(2):iyad058, 2023. doi: 10.1093/genetics/iyad058.
